# Single molecule footprinting measures low nucleosome occupancy in mature spermatozoa of mice and men

**DOI:** 10.64898/2026.06.30.735528

**Authors:** Laura Gaspa-Toneu, Haojun Shi, Evgeniy A. Ozonov, Mark. E. Gill, Christian De Geyter, Antoine H.F.M Peters

## Abstract

Nucleosomes are fundamental units of DNA packaging and gene regulation in eukaryotes. In mammalian sperm, most nucleosomes are replaced by protamines causing extreme chromatin compaction. Various epigenomic studies reported conflicting results on the distribution of residual nucleosomes in mammalian sperm^1–9^, questioning their potential role in mediating intergenerational inheritance of paternal epigenetic information^10,11^. Here we performed single-molecule footprinting through Nucleosome Occupancy and Methylome (NOMe) sequencing^12^ and applied the Bayesian statistical model nomeR^13^ to determine frequencies of nucleosome removal and retention at 103 specific genomic regions in thousands of developing haploid spermatids and mature spermatozoa of mice. While we readily detected footprints of nucleosomes and the transcription factor CTCF in round spermatids, chromatin became transiently highly accessible in elongating spermatids with loss of such footprints, indicating extensive chromatin reprogramming during spermiogenesis. In mature sperm, following nuclear decondensation with recombinant nucleoplasmin, we measured nucleosome occupancy frequencies ranging ∼1.2 to 1.7% at mouse loci. In human sperm, nucleosome occupancy varied between ∼2.3 to 4.5% at 163 genomic loci profiled. Contrasting mice, chromatin in ∼25% of human sperm was accessible upon reducing disulfide bonds between protamines arguing for species specific protamine packaging. Our findings support a stochastic rather than programmed potential role of residual nucleosomes in mammalian sperm in regulating paternal gene expression during ensuing embryonic development.

## Introduction

In mammals, parental genomes in early embryos are epigenetically distinct despite their genetic resemblance. The dimorphism in parental chromatin originates from sex specific chromatin formation during gametogenesis. In oocytes, genomes are packaged by nucleosomes, the fundamental unit of DNA packaging in eukaryotic cells. They carry oocyte specific histone variants and posttranslational modification (PTMs). Following fertilization, maternal chromatin becomes reprogrammed in a genomic location and species-specific manner^14–17^. In rodents, certain maternally inherited PTMs confer epigenetic information to offspring, e.g. in forming constitutive heterochromatin at pericentromeres^18,19^ or controlling imprinted X-inactivation^20^ and non-canonical genomic imprinting^21^.

In contrast, most nucleosomes are replaced by protamines by the end of spermatogenesis. Protamines are highly basic proteins that neutralize the negative charge of DNA thereby driving extreme compaction of sperm nuclei and protecting sperm DNA from damage until fertilization when they are exchanged by oocyte provided nucleosomes^22,23^. PTMs on protamines have been proposed to control the degree of packaging of mouse spermatozoa^24^ and their removal after fertilization^25^. Nonetheless, the histone-to-protamine exchange is incomplete in mammals, resulting in residual histone levels of ∼1-2% in mice and up to ∼10-15% in human sperm^1–3,26^. In human but not in mice, paternal inheritance of H3K9me3-modified nucleosomes was reported at pericentromeric and Y chromosome-linked Satellite II/III sequences via sperm to zygotes, as based on immuno-FISH experiments^18,27^. Beyond constitutive heterochromatin, it is unknown whether histones and their modifications function in paternal transmission of epigenetic information.

Early biochemical studies suggested that specific regions of the sperm genome are either preferentially packaged with histones or protamines^23,26,28–31^. Subsequent micrococcal nuclease-sequencing (MNase-seq) and chromatin immunoprecipitation-sequencing (ChIP-seq) experiments of sperm showed overrepresentation of nucleosomes at gene-regulatory regions consisting of CpG-rich, unmethylated sequences, particularly at promoters and exons, while being underrepresented at introns, intergenic regions, and repeat elements^1–3,31^. Since these nucleosomes also carried regulatory histone marks such as H3K4me3 and H3K27me3, they were proposed to possibly contribute to epigenetic inheritance and support gene expression post-fertilization^1–3^, a notion that was further investigated in follow-up studies^5–7,32–39^.

These studies have been challenged by another study reporting under-representation of nucleosomes at promoters and gene-dense regions^9^. Differences in solubilization pre-treatments of sperm nuclei, enzymatic digestions of sperm chromatin, and antibody pull-down efficiencies may underlie reported discrepancies^10,11^. Since then, alternative decondensation and solubilization methods were developed^33,40^. For example, recombinant nucleoplasmin (NPM) was used to effectively remove protamines and enhance chromatin accessibility^8,25,41,42^. NPM is highly abundant in oocytes and facilitates removal of protamines from sperm nuclei during fertilization allowing their replacement with maternally provided histones^43,44^. In NPM-decondensed sperm, histone H3 and H3K4me3 were over-represented at promoters of developmental and spermatogenic genes in ChIP-seq experiments and correlated with CpG content while histone H4 showed enrichment at intergenic loci, especially those overlapping repeats^8^. Other ChIP-seq studies reported CTCF binding to sperm chromatin of mice, with adjacent well-phased nucleosomes carrying active histone marks and chromatin loops forming topologically associated domains (TADs)^5,9,34,45–47^. In human sperm, however, CTCF binding is unclear, inferred by ATAC-seq but not detected by protein blot analyses, consistent with the reported absence of TADs^34,48,49^. Finally, a recent study called many reports on sperm chromatin into question, arguing that their findings may have been skewed by contamination from cell-free chromatin^4^.

Here, we employed the Nucleosome Occupancy and Methylome sequencing (NOMe-seq) assay^12^ to obtain single-DNA-molecule readout of nucleosome occupancy and CTCF binding footprints at selected gene regulatory and other loci in progenitor male germ cells undergoing histone-to-protamine replacement and in mature sperm. The method involves treating nuclei with the exogenous GpC methyltransferase M.CviPI, which methylates accessible GpC dinucleotides while protein-bound regions remain unmethylated, revealing single-molecule footprints (SMFs) after bisulfite conversion. The assay measures simultaneously endogenous DNA methylation at CpG dinucleotides on the same DNA strands, offering insights into possible relationships between DNA methylation and protein occupancy. The retention of nucleosomes and histone marks in mammalian sperm has been proposed to play a programmed role in supporting gene expression and development of the next generation^1–3,28,50^. Under this premise, most individual spermatozoa should retain nucleosomes at key loci. Our results, however, argue against a programmatic retention of nucleosomes in sperm, as they show overall low nucleosome occurrence in both mouse and human sperm, indicating limited potential for high-penetrance paternal transmission of nucleosome-encoded epigenetic information.

## Results

### Comprehensive chromatin remodeling during spermatid development

To assess possible contributions of genomic location and sequence, including CpG density, DNA methylation and histone modifications in modulating nucleosome eviction or retention during spermiogenesis, we selected 103 genomic regions, grouped into 14 amplicon categories (Fig. 1a-b, Ext Data Fig. 1a-c, Table S1)^1,3,5,9^. Amplicon categories included gene promoters (TSS +/- 500bp), intergenic regions, putative enhancers elements, CTCF binding sites and imprinted control regions (ICRs), representing the non-repetitive part of the genome. We further classified amplicons according to their CpG dinucleotide density into high, medium or low CpG groups and to their CpG methylation levels into hypomethylated (≤ 25%) or hypermethylated (> 75%) states. They represent regions marked by different histone variants and post-translational modifications such as H3K4me1, H3K4me3, H3K9ac, H3K9me3, H3K27ac and H3K27me3 in round spermatids (RSts) and sperm as reported before in MNase-and ChIP-seq. studies (Ext Data Fig. 1a-c, Table S1)^3,5,8,34,51–53^ and are located on different chromosomes of the genome (Ext. Data Fig. 1d). The average length of amplicons covered by informative GpC positions (GCH trinucleotides: GCA, GCC and GCT) was 474 bp (range: 384 – 532 bp), ensuring detection of at least one to two nucleosome(s) within a read. Amplicons contained on average one GCH trinucleotide every 12.6 bp (range: 8.5 - 21.8 bp) with an average maximum gap of 33.6 bp (range: 25 – 40 bp), thereby supporting high-resolution detection of nucleosomes and transcription factors like CTCF (Ext. Data Fig. 1e, Table S1).

**Fig. 1:**
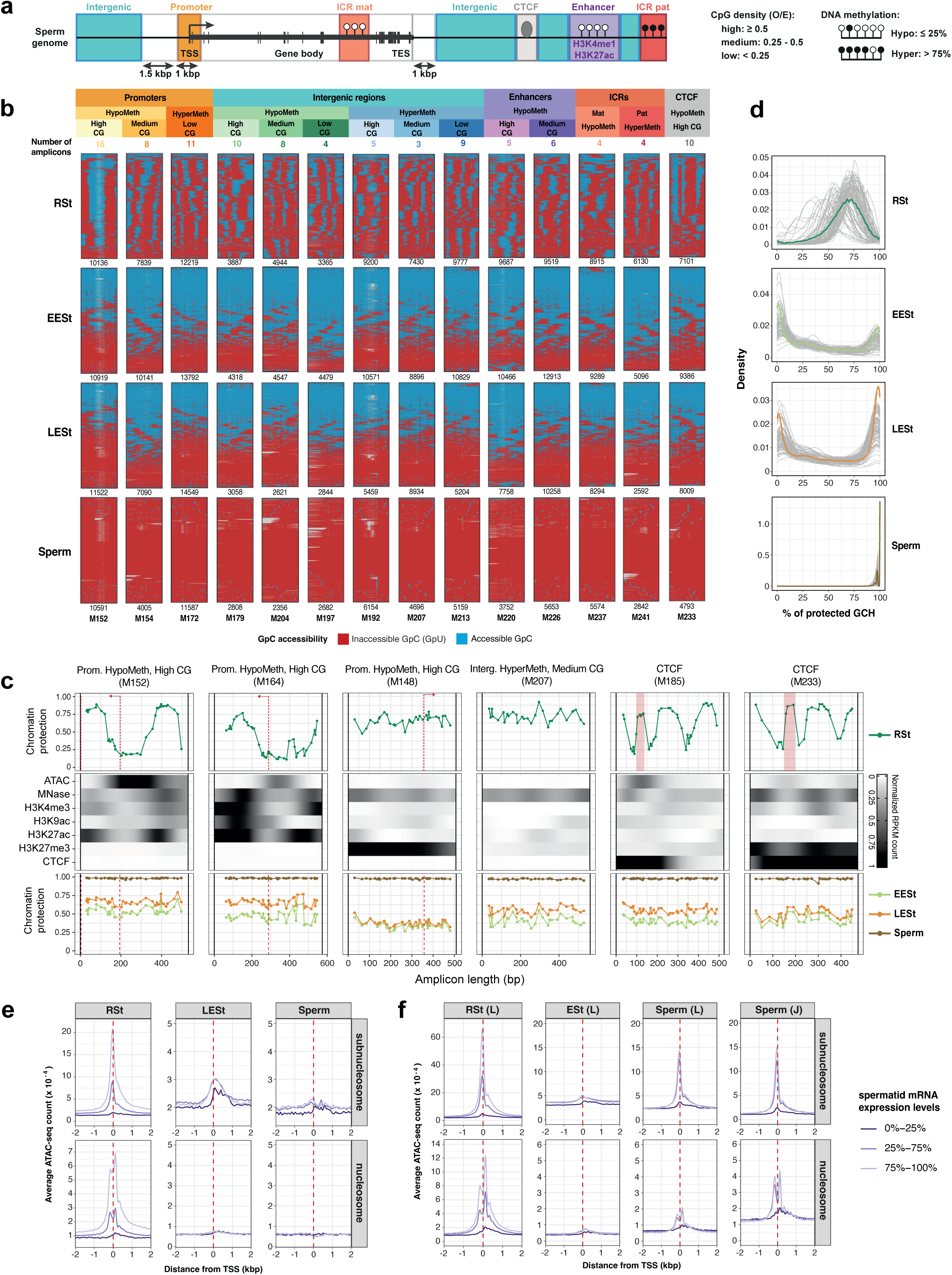
Extensive chromatin remodeling during spermatid development. **a.** Scheme illustrating genomic location, sequence and chromatin features of amplicon categories in mice. CpG density was defined by the ratio of observed to expected CpG dinucleotides (CpG-O/E). Endogenously methylated and non-methylated CpG dinucleotides are shown as black- and white-filled lollipops, respectively. TSS: transcription start site. TES: transcription end site. ICR: imprinting control region. **b.** Visualization of chromatin footprints in individual male haploid germ cells. Typical amplicons belonging to fourteen categories representing distinct genome locations and chromatin states are shown. Each row of a SMF-map corresponds to an individual fragment and each column to a cytosine within GCH context. Accessible (methylated) GCHs are shown in blue and inaccessible, protected (unmethylated) GCHs in red. Nucleotides between cytosines are not shown. Numbers of analysed reads and amplicon numbers (M152 etc) are indicated below plots. **c.** Average chromatin protection profiles at 6 representative amplicons in four cell types. Protection data are presented as the inverse of % GCH methylation (1 - % GCHme). ATAC-seq^58^, MNase-seq^51^ and ChIP-seq normalized counts are displayed for RSts (H3K4me3, H3K27me3^3^, H3K9ac, H3K27ac^52^, CTCF^53^). Position of TSS and CTCF motifs are indicated with dashed red lines and orange boxes, respectively. **d.** Density plots showing distributions of percentages of protected GCHs within molecules of individual (grey) and all (coloured) amplicons in indicated germ cell types. In RSts, most amplicons show ∼50-80% protection of GCHs, reflecting nucleosome protection and accessible linker DNA. The largely dual modal distributions in ESts indicate large fractions of cells with either fully accessible or fully inaccessible chromatin at all interrogated genomic regions. **e.** Averaged ATAC-seq-based chromatin accessibility at TSSs (+/- 2 kb) (in CPM) in RSts, LESts and sperm classified according to mRNA-seq-based expression quantiles in RSts or LESts respectively^87^. Subnucleosomal (<100 bp) and nucleosomal (160 - 250 bp) fragments were counted separately. **f.** Averaged ATAC-seq profiles measured in RSts, ESts and sperm^58^ and sperm^34^ classified according to mRNA-seq-based expression as shown in **e.**

To characterize nucleosome dynamics during spermiogenesis and position in mature sperm, we isolated mouse RSts, early and late elongating spermatids (EESts, LESts), and caudal epididymal sperm by FACS with 98.33 to 99.98% purity (Ext. Data Fig. 1e-f)^54^. We performed NOMe-seq on three independent biological replicates (Ext. Data Fig. 1e). At profiled amplicons, levels of GCH methylation reflecting chromatin accessibility correlated highly among biological replicates of RSts, moderately to highly among EESt and among LESt replicates but lowly between sperm samples (Ext. Data Fig. 1g). Chromatin accessibility varied extensively between the three different spermatid types and sperm, suggesting extensive chromatin remodeling during nuclear elongation. In contrast, endogenous WCG methylation correlated highly among cell types (Ext. Data Fig. 1h), consistent with reported stability of DNA methylation during spermatogenesis^52,55^.

In RSts, we reproducibly observed nucleosome-like protected footprints in amplicons profiled (Fig. 1b), consistent with findings in somatic cells^12,13,56^. At transcription start sites (TSS) of promoter regions, well-positioned nucleosome-like footprints overlapped with mono-nucleosomes mapped by MNase-seq and ChIP-seq of active histone marks, aligning with −1 and/or +1 nucleosomes adjacent to active TSS (Fig 1c, M152, M164) (Krebs et al., 2017). We occasionally observed at TSS shorter footprints co-localizing with ATAC-seq chromatin accessibility that may represent transcription factor or transcriptional machinery binding (Fig 1c, M164)^56–58^. Short footprints surrounded by well-positioned nucleosome-like footprints were also observed in genomic regions marked by high CTCF ChIP-seq enrichment (Fig. 1c, M185 and M233), suggesting nucleosome phasing following CTCF binding^53^. In contrast, loci marked by H3K27me3 (M148) or DNA methylation (M207) contained nucleosome-like footprints but lacked nucleosome positioning and had low ATAC-seq accessibility (Fig. 1c). Together, our NOMe-seq data recapitulates and extends chromatin occupancy data measured in RSts by conventional bulk profiling methodologies.

In contrast to RSts, nucleosome-like footprints were only observed in a relatively small fraction of reads in EESts and LESts. Instead, considerable proportions of single molecules were either fully accessible, lacking any distinctive large chromatin footprints, or were fully inaccessible (Fig. 1b). The proportions of fully accessible and inaccessible reads were similar between all amplicons (Fig. 1d), pointing to an overall similarity in chromatin remodeling states between the genomic regions profiled. Interestingly, positioning of nucleosomes at promoters previously active in RSTs was no longer observed (Fig. 1b). The loss of positioning could be due to nucleosome repositioning linked to global transcriptional shutdown in late spermiogenesis or to nucleosome eviction. Likewise, CTCF footprints and adjacent well-positioned nucleosomes were also not detectable (Fig. 1b-d), indicating that CTCF binding at these regions is largely lost, and nucleosomes are repositioned or evicted. In sperm, single molecules were generally completely inaccessible, without detectable nucleosome-like footprints (Fig. 1b-d).

The high frequency of fully accessible reads measured for all amplicons in elongating spermatids was surprising. To explore this observation further, we performed ATAC-seq on RSts, LESts- and sperm populations isolated by FACS-sorting as done for NOMe-seq. In parallel, we reanalyzed published ATAC-seq data of spermatids and sperm^34,58^. While multiple hypomethylated amplicon regions were accessible by ATAC-seq in RSts, we and others^58^ failed to measure such differential accessibility in LESts (Ext. Data Fig. 1i). Genome-wide analyses recapitulated high accessibility surrounding subnucleosomal and nucleosomal footprints at TSSs in RSts which related to their transcriptional states. In contrast, such accessibility was low or absent in LESts (Fig. 1e-f; Ext. Data Fig. 1j)^58^, arguing for either fully open or closed chromatin. Published ATAC-seq data reported regain of differential sub-/nucleosomal accessibility at loci in mature sperm samples, including TSS (Fig. 1f)^34,58^. Despite testing multiple protocols and conditions, we could not recapitulate these results and measured evenly low accessibility at TSSs in sperm, aligning with our NOMe-seq observations (Fig. 1b,1e). Our ATAC-seq results in LESts and sperm concur with our NOMe-seq findings and point to efficient widespread nucleosome eviction during nuclear elongation followed by tight compaction, presumably due to incorporation of protamines.

### Quantification of NOMe-seq footprints with a Bayesian probabilistic model

Visualization of NOME-seq data showed that genomic regions in elongating spermatids exist in three different chromatin states, i.e. carrying nucleosomes, being accessible or inaccessible (Fig. 1b). The data further indicates that the relative frequencies of chromatin states are comparable between genomic regions. To systematically call and quantify the different NOMe-seq chromatin footprints and detect potential differences between amplicon classes in an unbiased manner, we applied nomeR - a Bayesian probabilistic model that allows inference of footprint lengths and prediction of footprint locations within individual molecules^13^.

Based on calculated posterior probabilities, the model called four footprint types: (i) *nucleosome*: reads containing at least one nucleosome (100-150 bp) with high predicted probability; (ii) *fully accessible*: reads with mostly unprotected GCHs; (iii) *fully inaccessible*: reads with most GCHs protected; and (iv) *partially accessible*: reads lacking nucleosomes but potentially containing a TF footprint (30 – 80 bp) or other footprints starting at the edge of the amplicon window, with unknown lengths (Fig. 2a). Reads of the partially accessible class may represent boundaries between fully accessible and fully inaccessible regions within the genome or may contain nucleosomes at their edges. The class nucleosome therefore defines with high confidence and stringency the fraction of nucleosome-containing molecules for each locus and cell type analyzed.

**Fig. 2:**
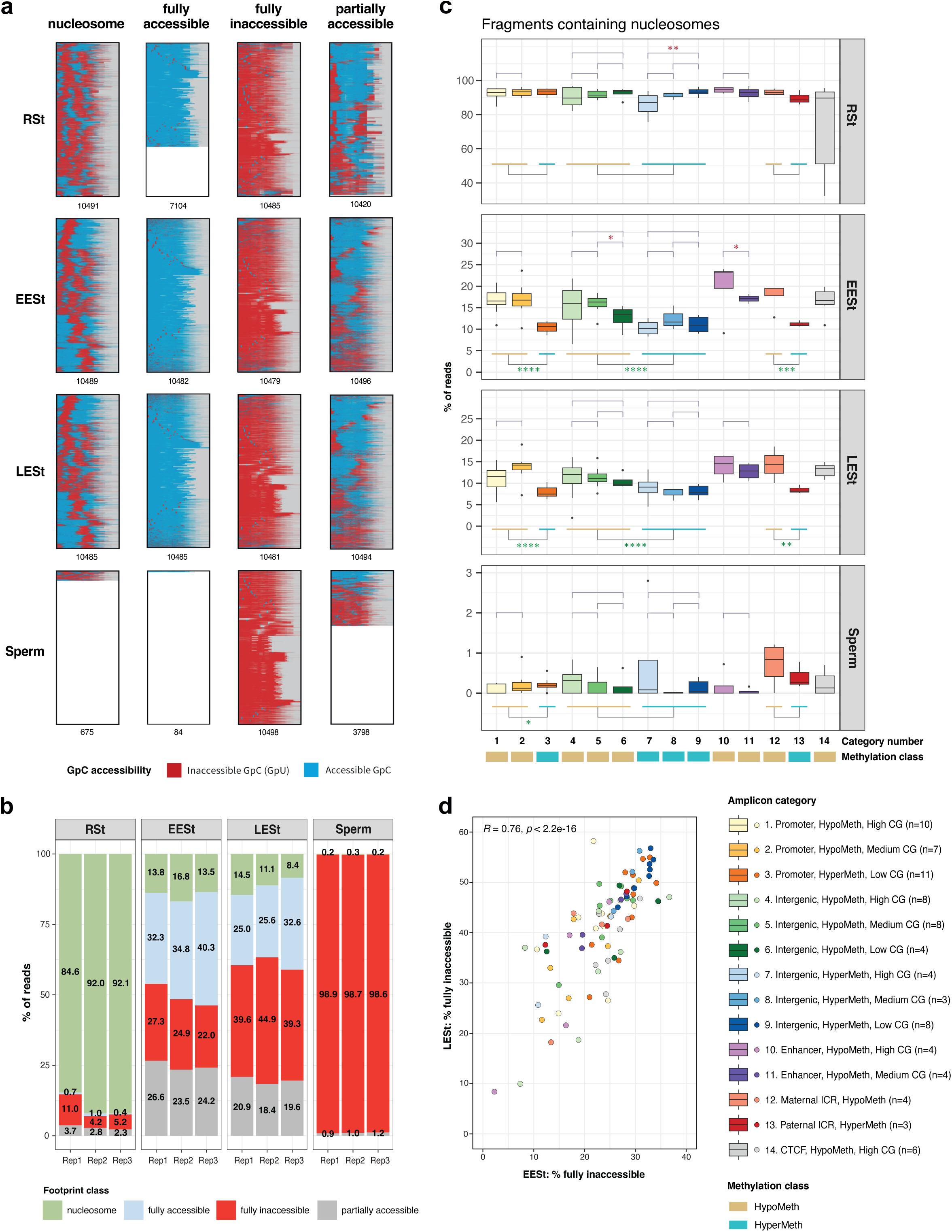
Native chromatin of mouse spermatozoa is inaccessible to exogenous GpC-methylation. **a.** NOMe-seq. SMF-maps visualizing single-molecule footprinting data reads classified as nucleosome, fully accessible, fully inaccessible or partially accessible according to the probabilistic NOMeR model. For each footprint class and cell type, a maximum of 10,500 fragments randomly selected from all amplicons (3500 fragments/replicate) are shown. Amplicon lengths and total number of sampled reads were equalized by the addition of NA values (grey and white, respectively). Each row of a SMF-map corresponds to a unique sequence read and each column to a cytosine within GCH context. Accessible GCHs (methylated) are shown in blue and protected GCHs (unmethylated) in red. **b.** Bar plots displaying the average percentage of fragments containing nucleosome, fully accessible, fully inaccessible or partially accessible footprints at all amplicons per cell type for each replicate sample. **c.** Box plots representing the distribution of nucleosome occupancy frequencies at amplicons belonging to 14 amplicon categories in four germ cell types. Boxes display, 25^th^, 50^th^ and 75^th^ percentiles and whiskers extend to 1.5 times the inter-quantile range. As indicated, statistical significance was evaluated using two-sided Wilcoxon rank-sum tests. Comparisons were performed among promoter, intergenic and enhancer amplicon categories with different GC levels. Likewise, between hypomethylated and hypermethylated amplicon regions for promoter and intergenic categories regardless of their CG levels, as well as between maternal and paternal ICRs. ns: p>0.05, *: p≤0.05, **: p≤0.01, ***: p≤0.001, ****: p≤0.0001. **d.** Scatterplot relating frequencies of fully inaccessible footprints in EESts and LESts across all amplicons. The corresponding Pearson correlation coefficient (*R*) and associated p-value are indicated in the upper left corner.

Footprint prediction revealed high abundances of nucleosome footprints in RSts, frequently with two nucleosomes per read (Fig. 2a, Ext. Data Fig. 2a-b). Shorter, discrete footprints were also observed, particularly at CTCF motif containing amplicons (e.g. M185, M233; Ext. Data Fig. 2b). In contrast, amplicons in ESts displayed markedly lower frequencies of nucleosome footprints, representing often single nucleosomes. Short footprints (<10 bp) and footprints larger than nucleosomes (>150 bp) were instead more prevalent in ESts (Fig. 2a, Ext. Data Fig. 2a-b). Thus, the molecule categorization generated by the model aligns well with earlier observations from footprint visualization (Fig. 1b).

### Dynamic chromatin remodeling during spermiogenesis

The prediction and categorization of footprints allowed us to quantify dynamic changes in footprint frequencies during spermiogenesis. In three biological replicate samples of RSts, ∼85% to 92% of all reads contained on average nucleosome footprints. During subsequent nuclear elongation, nucleosome footprint levels decreased 6- to 8-fold, to 13-17% of reads in EESts and 8-14% in LESts (Fig. 2b-c, Ext. Data Fig. 2a). Remarkedly, on average, ∼36% of reads in EESts and ∼28% in LESts represented fully accessible molecules which were barely present in RSts. We observed such extensive remodeling at all amplicons arguing for a genome wide process (Ext. Data Fig. 2c). The loss of nucleosomes was more extensive among low CG and CpG-methylated amplicons in ESts, suggesting earlier or more effective removal of nucleosomes from such sequences (Fig. 2c). By comparing percentages of fully inaccessible, closed reads measured in EESts and LESts, chromatin appeared to close with comparable rates at most amplicons (Fig. 2d, Ext. Data Fig. 2c). For those amplicons closing less efficiently in EESts, subsequent closing rates were higher (Ext. Data Fig. 2d).

Interestingly, in fully and partially accessible reads of EESts and LESts, we observed minute protected footprints of variable size scattered throughout amplicon regions (Ext. Data Fig. 2a). We speculate that these may reflect binding of single or small assemblies of proteins like transition proteins and/or protamines to DNA molecules. Fully inaccessible reads, which increased from an average of ∼7% in RSts to ∼25% in EESts, and ∼41% in LESts, reached over 99% in sperm (Fig. 2b-c; Ext. Data Fig. 2c). They may represent chromatin packaged by protamine molecules, getting progressively intra- and intermolecularly connected via oxidized disulfide bonds into compact, stable and inert structures.

Together, our NOMe-seq data point to a transient opening and subsequent closing of chromatin at 103 amplicon regions located throughout the genome during spermatid elongation and sperm formation lasting for days in mice. Since NOMe-seq reads represent chromatin states of individual haploid germ cells, the observed variabilities between reads within populations of EESts or LESts suggests that the remodeling from a nucleosome-to-protamine state initiates randomly at a given amplicon while it successively progresses in a manner comparable between amplicons during the process of nuclear elongation. Our data further suggest that the removal of nucleosomes and the repackaging by protamines into closed chromatin are two temporally separable processes.

### Nucleosome eviction in ESts is slightly advanced by endogenous DNA methylation

The absence of DNA methylation at CpG-rich sequences was previously associated with increased levels of nucleosome retention in mouse and human sperm^1–3,55^. To assess the possible impact of CpG density, endogenous DNA methylation and genomic locations on variability in nucleosome turnover dynamics during nuclear elongation in ESts, we compared nucleosome frequencies at amplicons belonging to the 14 categories previously described (Fig. 1b). In RSts, we measured an overall high frequency of nucleosomes in all amplicons categories. During elongation, we measured a significant 1.7-fold higher frequency of nucleosomes at hypomethylated CG-rich/intermediate promoter regions compared to hypermethylated CG-poor promoter regions (categories 1-3 in Fig. 2c; Ext. Data. Fig 2a-c). These data suggest that during elongation, higher CpG density may promote nucleosome retention, that endogenous DNA methylation may promote nucleosome eviction, or that both characteristics contribute to the process of nucleosome removal. To investigate these options, we compared nucleosome frequencies at intergenic regions with different CpG compositions and being either hypo- or hypermethylated at CpG dinucleotides (categories 4-6 vs. 7-9 in Fig. 2c; Ext. Data. Fig 2a-c). This latter comparison supports the view that DNA methylation promotes nucleosome eviction during spermatid elongation. Nucleosome frequencies at hypomethylated CG-rich/intermediate putative enhancer and maternal imprinting control regions versus hypermethylated paternal ICRs further support this conclusion (categories 10-13 in Fig. 2c; Ext. Data. Fig 2a-c). Based on these findings, we propose that the pace of nucleosome eviction is not uniform throughout the mouse genome and that it is somewhat modulated by endogenous DNA methylation.

### Nuclear decondensation with recombinant nucleoplasmin unmasks low abundance of nucleosome footprints in mouse sperm

While many elongating spermatids were open to chromatin footprinting, mouse spermatozoa were refractory. To overcome such inaccessibility, we explored methods to decondense sperm nuclei. We first employed heparin, a negatively charged glycosaminoglycan that was previously used to remove protamines from human and mouse sperm^29,59^. By microscopy, we observed efficient decondensation of FACS sorted mouse sperm following heparin treatment (Ext. Data Fig. 3a). By NOMe-seq, chromatin became fully accessible, but we observed no nucleosome footprints (Ext. Data Fig. 3b). Remarkedly, we failed to observe nucleosome footprints in RSts after heparin treatment, thus disqualifying heparin for use in native nucleosome footprinting experiments. Next, we permeabilized and decondensed sperm with detergents that we had previously applied in ChIP-seq and MNase-seq studies^60^. Unlike heparin, this method did not significantly affect histone stability. Unfortunately, even upon prolonged nuclear decondensation, many sperm molecules remained inaccessible to footprinting, thereby precluding a quantitative assessment of nucleosome frequencies in mouse sperm. For reads that gained accessibility, we did not detect nucleosome footprints (Ext. Data Fig. 3b).

Lastly, we tested reduction of protamine disulfide bonds with dithiothreitol (DTT) followed by treatment of chromatin with recombinant *Xenopus* nucleoplasmin (NPM)^8^. Amphibian NPM and mammalian NPM2 proteins are abundantly expressed in oocytes and facilitate protamine removal from sperm upon fertilization^43,44^. *In vitro*, treatment of permeabilized sperm with NPM leads to protamine displacement and decondensation while nucleosomes are retained^8,25^. We produced *Xenopus* NPM recombinantly and incubated mouse sperm with it for various durations (15, 45 and 90 minutes) after cell lysis and before M.CviPI-mediated methylation (Fig. 3a). Reduction of protamine disulfide bonds with DTT treatment itself did not decondense sperm nuclei nor did it significantly increase chromatin accessibility at sampled loci (Fig. 3a-d). Instead, additional NPM treatment caused sperm nuclei to decondense over time and chromatin gradually gained accessibility to footprinting unveiling the four chromatin states previously observed in ESt populations: nucleosomes, fully accessible, partially accessible and fully inaccessible (Fig. 3c-d). Hence, NPM-mediated decondensation was suitable to quantify nucleosome occupancy in decondensed sperm.

**Fig. 3:**
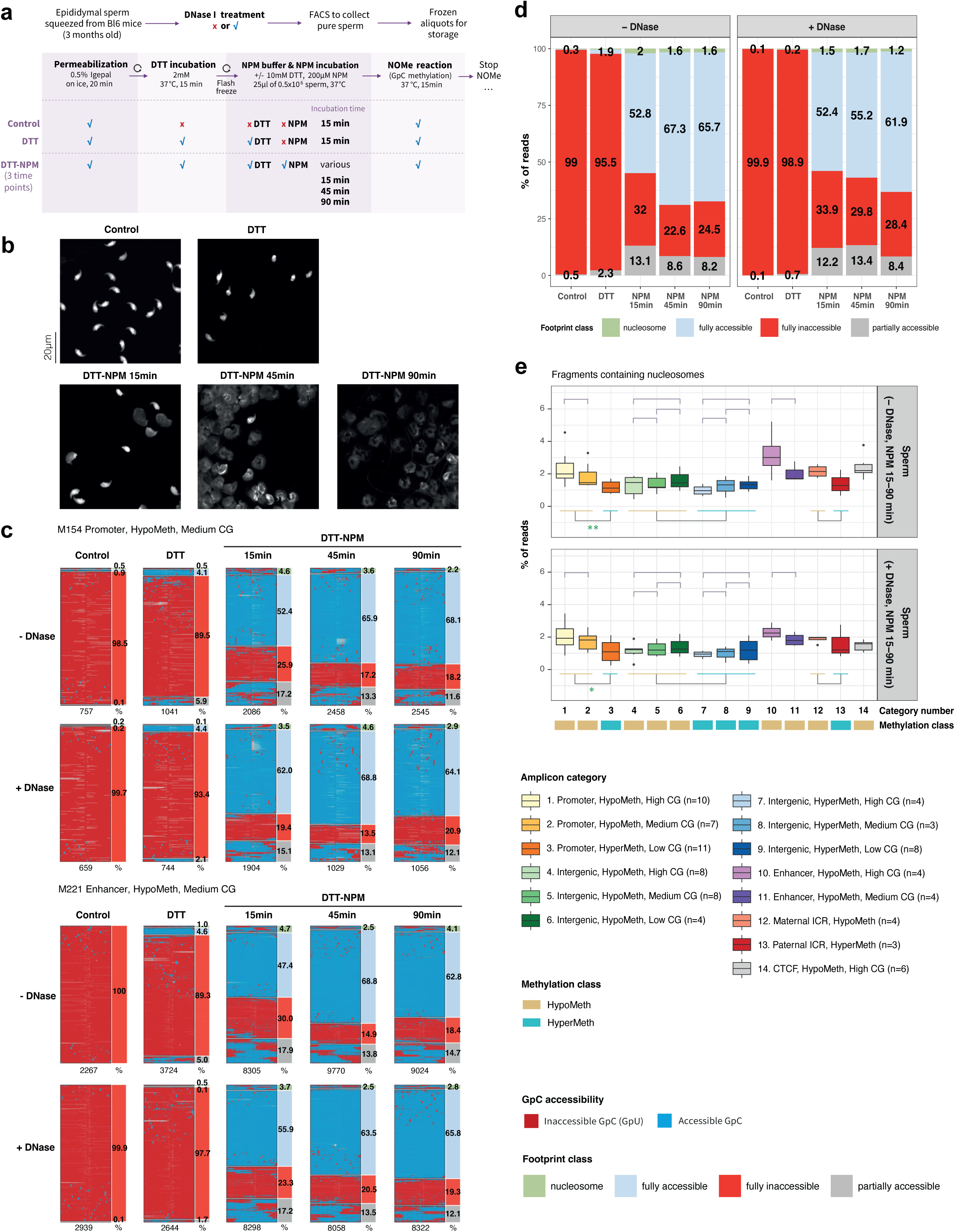
Low nucleosome occupancy at regulatory regions in decondensed mouse spermatozoa. **a.** Diagram outlining pretreatment of mouse spermatozoa without or with DNase I, FACS-purification, permeabilization and/or exposure to recombinant nucleoplasmin (NPM) prior to GpC methylation. **b.** Microscopy images of DAPI-stained mouse sperm nuclei subjected pretreatment conditions described in panel 3a. **c.** NOMe-seq. SMF-maps for 2 representative amplicons visualizing chromatin footprints at DNA molecules of individual spermatozoa, that had been classified according to the NOMeR model. Frequencies of predicted footprint classes are on the left of each SMF-map. Sperm was pretreated without or with DNAse I and further as indicated. Every row corresponds to a unique sequence read and each column to a cytosine in a GCH context. Nucleotides between cytosines are not shown. Each row of a SMF-map corresponds to an individual fragment and each column to a cytosine within GCH context. Accessible GCHs (methylated) are shown in blue and protected GCHs (unmethylated) in red. The number of analysed reads is shown at the bottom of each SMF-map. **d.** Bar plots displaying the average percentage of fragments containing nucleosome, fully accessible, fully inaccessible or partial accessible footprints at all amplicons in FACS-sorted control, DTT or DTT-NPM-exposed mouse spermatozoa, pretreated without or with DNase I. **e.** Box plots representing the distribution of nucleosome occupancy frequencies at amplicons belonging to 14 amplicon categories in FACS-sorted and DTT-NPM-exposed mouse spermatozoa, pretreated without or with DNase I. Data of all NPM duration conditions are shown. Boxes display, 25^th^, 50^th^ and 75^th^ percentiles and whiskers extend to 1.5 times the inter-quantile range. Statistical significance was evaluated as in Fig. 2C using two-sided Wilcoxon rank-sum tests. ns: p>0.05, *: p≤0.05, **: p≤0.01.

To ensure that the epididymal sperm we used to be free from somatic contamination, which could majorly distort nucleosome occupancy quantification^4^, we performed parallel NOMe-seq experiments using sperm treated with or without DNase I prior to FACS purification, cell lysis, NPM decondensation and footprinting (Fig. 3a, Ext. Data Fig. 3a). Quantification of footprint abundances with nomeR showed that DNAse I pre-treatment did not significantly change the frequency of the four footprint types, indicating high purity of the sorted sperm samples (Ext. Data Fig. 3c-d). While ∼22–28% of reads remained inaccessible to footprinting, ∼62-67% of molecules had fully accessible chromatin and were devoid of nucleosome footprints in both conditions (Fig. 3c-d; Ext. Data Fig. 3c-d). Nucleosomes were detected at very low occurrence, averaging to only ∼1.6-2.0% in untreated sperm and slightly lower in sperm treated with DNase I (∼1.2-1.7%) (Fig. 3d).

When comparing footprint frequencies across 14 amplicon categories in NPM-treated sperm (Fig. 3e), the small variabilities in nucleosome occupancies between amplicon classes that we had previously measured in EESts and LESts were further diminished in decondensed mature sperm, arguing for an almost complete removal of nucleosomes (Fig. 2c, 3e). We nonetheless observed slightly higher residual nucleosome occupancies at certain hypomethylated CG-rich amplicons, particularly in hypomethylated promoters, enhancers, maternal ICRs, and CTCF-containing amplicons (Fig. 3e). This underscores a minor yet lasting influence of endogenous DNA methylation on nucleosome retention during sperm formation.

Despite the overall low occurrence of nucleosomes among all amplicons in sperm, some amplicons reached up to ∼6% (Fig. 3e; Ext. Data Fig. 3e-g). To investigate the reproducibility of nucleosome occupancies, we calculated the mean and standard deviation of nucleosome occupancies for each amplicon across all NPM incubation conditions, DNase treatments, and replicates (Ext. Data Fig. 3h). We observed a few amplicons with nucleosome occupancies exceeding one standard deviation above the mean value across all amplicons. Such outliers, however, exhibited higher variability across samples, exceeding the standard deviations of most other amplicons. These data argue that outlier amplicons do not represent regions with programmed above-average nucleosome occupancy. Instead, the data highlights the largely stochastic nature of nucleosome eviction versus retention and does not support high-penetrance paternal transmission of nucleosome-encoded epigenetic information via typical gene regulatory sequences and ICRs.

### CTCF occupancy is early lost during nuclear elongation

While CTCF occupancy was readily detectable in RSts, we measured only infrequently short, protected footprints at some CTCF motifs in EESts, even less frequently in LESt and being undetectable in untreated or NPM-mediated decondensed sperm (Ext. Data. Fig 2b, 3e). Hence, we cannot recapitulate CTCF occupancy at tested loci in mouse sperm as was previously reported in MNase-ChIP-seq experiments (Ext. Data. Fig 1b)^34^.

### Nucleosome occupancy in human sperm is consistently low among donors

To investigate the prospect of paternal inheritance by sperm-borne nucleosomes in human, we extended our NOMe-seq analyses to sperm samples from three human donors with semen parameters within the range of fertile men (Ext. Data Fig. 4a). Spermatozoa of humans have been reported to contain higher levels of histones (∼10-15%) compared to those of mice (∼1-2%). Moreover, human sperm chromatin has been notably easier to decondense^1,2,26,60^. As for mouse, we designed NOMe-seq amplicon assays for 163 regions classified into 25 categories representing promoter, intragenic, intergenic, enhancer and ICR regions with distinct DNA hypo- and hypermethylation and histone PTM states, different CpG compositions and positioned on all autosomes and the X chromosome (Ext. Data Fig. 4b-d, Ext. Data Table 2). The average length of amplicons covered by informative GpC positions (GCH trinucleotides: GCA, GCC and GCT) was 501 bp (range: 337 – 532 bp). Amplicons contained on average one GCH trinucleotide every 11.1 bp (range: 7.7 - 18.4 bp) with an average maximum gap of 31.7 bp (range: 20 – 37 bp), ensuring detection of one to two nucleosome(s) within a read.

We performed microscopy and NOMe-seq experiments comparing spermatozoa from semen samples, that were washed with media, then either treated with or without DNase I and/or subjected to FACS purification and lastly lysed to enable access of the GpC methyltransferase M.CviPI to chromatin (Ext. Data Fig. 4e,f). We used nomeR to call chromatin states in human sperm^13^. In control sperm samples without NPM decondensation, we observed accessible chromatin in ∼12-21% of reads of washed sperm samples of the three donors that had neither been treated with DNase I nor purified by FACS sorting (Fig. 4b,c; Ext. Data Fig. 4h,i). Upon FACS sorting or DNAse I treatment, however, the frequency of such accessible reads reduced to ∼8-10% and ∼2-3%, respectively. Combinatorial pretreatment resulted in ∼2% accessible reads only. These findings argue that FACS sorting removes contaminating somatic or immature germ cells. Moreover, DNase I treatment prior to FACS sorting was required to effectively remove DNA from damaged cells and/or contaminating cell-free chromatin. Together, these experiments highlight the importance of thorough sperm purification for quantitative chromatin analyses.

**Fig. 4:**
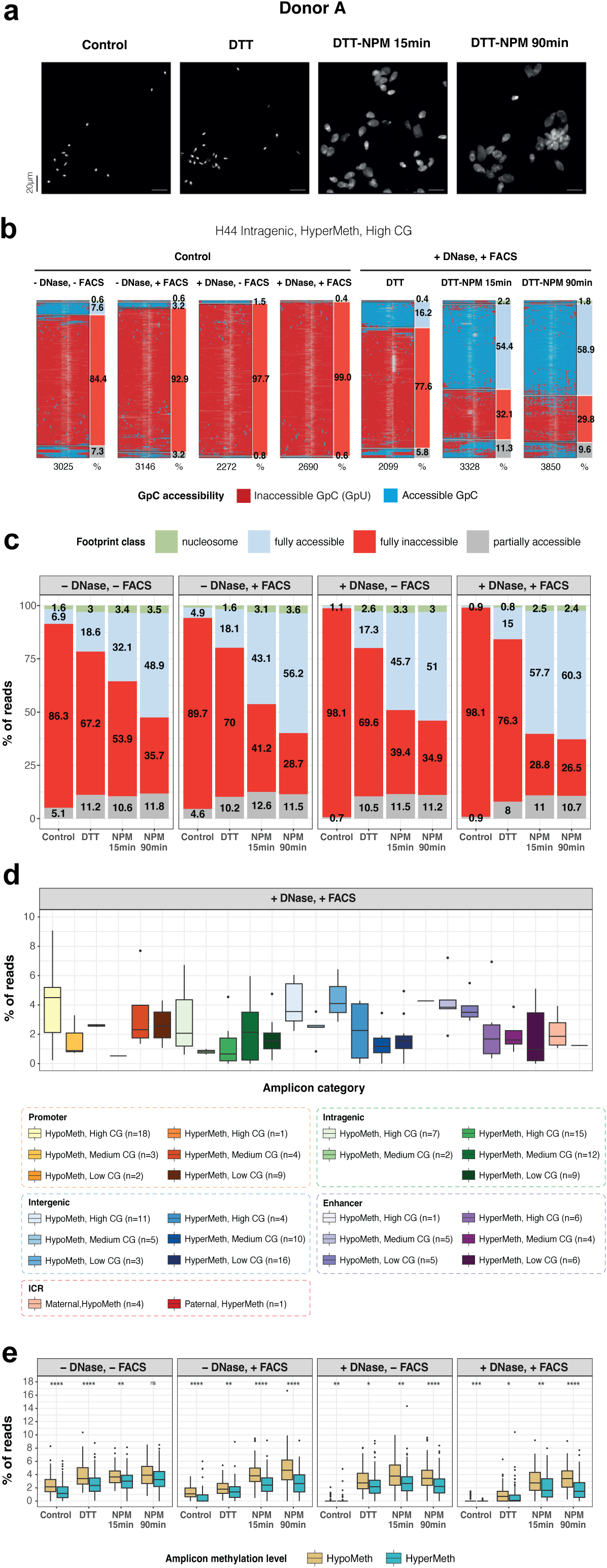
Limited accessibility with low nucleosome occupancy at genomic regions in human spermatozoa. **a.** Microscopy images of DAPI-stained sperm nuclei from human donor A. Sperm was pretreated with DNAse I, FACS-sorted and subjected to DTT and NPM treatment conditions as described in panel 3a. **b.** NOMe-seq. SMF-maps visualizing chromatin footprints at one representative amplicon for individual sperm molecules of human donor A, that had been classified according to the NOMeR model and annotated with frequencies of predicted footprint classes on the left of each SMF-map. SMF-maps corresponding to sperm pretreated without/with DNAse I, and without/with FACS sorting are illustrated for the control condition; SMF-maps corresponding to DNase I-treated, FACS-sorted sperm are further illustrated for DTT and NPM incubations as indicated. Every row corresponds to a unique sequence read and each column to a cytosine in a GCH context. Nucleotides between cytosines are not shown. Each row of a SMF-map corresponds to an individual fragment and each column to a cytosine within GCH context. Accessible GCHs (methylated) are shown in blue and protected GCHs (unmethylated) in red. The number of analysed reads is shown at the bottom of each SMF-map. **c.** Bar plots displaying the average percentage of fragments containing nucleosome, fully accessible, fully inaccessible or partially accessible footprints for all amplicons in control, DTT or DTT-NPM-exposed spermatozoa from human donor A, priorly treated without/with DNase I or without/with FACS sorting. **d.** Box plots representing the distribution of nucleosome occupancy frequencies at 163 amplicons belonging to 25 amplicon categories in DNase I-pretreated, FACS-sorted, DTT pretreated and 90 min NPM-exposed human spermatozoa of donor A. Boxes display, 25^th^, 50^th^ and 75^th^ percentiles and whiskers extend to 1.5 times the inter-quantile range. **e.** Box plots comparing the distribution of nucleosome occupancy frequencies at amplicons, grouped into hypomethylated and hypermethylated categories, in control, DTT or DTT-NPM-exposed spermatozoa from human donor A, priorly treated without/with DNase I or without/with FACS sorting. Boxes display, 25^th^, 50^th^ and 75^th^ percentiles and whiskers extend to 1.5 times the inter-quantile range. Statistical significance was evaluated using two-sided Wilcoxon rank-sum tests. ns: p>0.05, *: p≤0.05, **: p≤0.01, ***: p≤0.001, ****: p≤0.0001.

As in mice, reduction of protamine disulfide bonds with DTT by itself was not sufficient to decondense human spermatozoa (Fig. 4a; Ext. Data Fig. 4g). By contrast, DTT pretreatment of human spermatozoa did significantly increase chromatin accessibility to footprinting in ∼25% of reads, either becoming fully accessible (∼15%), partial accessible with undefined footprints (∼9%) or harboring ∼1-2% nucleosome footprints (Fig. 4b-d; Ext. Data Fig. 4h-j). Hence, upon reduction of disulfide bonds between protamines, a substantial part of the human genome unfolds and becomes accessible, revealing the absence of protamines at the profiled amplicon sites. These findings point to an extensive heterogeneity in chromatin composition and compaction of human spermatozoa, as previously noted^61^. This human data contrasts clearly to the uniformly compacted condition measured in spermatozoa of mice (Fig. 3c,d).

Subsequent treatment with NPM for 15 and 90 min expanded human sperm nuclei effectively by 10-20-fold (Fig. 4a) and the fraction of chromatin accessible to footprinting increased up to 75% with some variability between amplicons, duration of NPM treatment and donors, which presumably reflects intrinsic heterogeneity in chromatin packaging.

When profiling nucleosomes in doubly purified spermatozoa of three donors, we measured normal distributions of nucleosome occupancies over the 163 amplicons, with their overall mean ± standard deviation being 2.5±1.6%, 2.4±1.6% and 4.2±2.3% for donors A, B and C, respectively (Fig. 4d; Ext. Data Fig 4j,l,m). There were only a handful of amplicons with nucleosome frequencies outside the second standard deviation interval, suggesting that these are stochastic outliers. We observed variable nucleosome frequencies between the 25 different amplicon categories which also varied between donors (Fig. 4d; Ext. Data Fig 4j,l-n). Yet, grouping the amplicons according to endogenous DNA methylation levels showed that, like in mouse, hypomethylated regions had statistically significant ∼2-fold higher nucleosome frequencies than hypermethylated regions, even though remaining within the few percent range (Fig. 4e; Ext. Data Fig 4k). Together, our data unambiguously demonstrate that in mice and men, nucleosome frequences at gene regulatory and intergenic regions are low in sperm. Our data supports the view that nucleosomes are unlikely to function as a programmed means to transmit epigenetic information between generations.

## Discussion

In this study we answer the fundamental question regarding the occupancy of nucleosomes across genomic loci in individual mammalian spermatozoa. Using NOMe-seq we measured chromatin states and quantified nucleosome occupancy frequencies at 103 and 163 genomic sites located throughout the genomes of spermatozoa of mice and men, respectively. Profiling was done at deep coverage, representing hundreds to thousands of unique single molecules isolated from haploid spermatids and caudal epididymal sperm from mice and human ejaculated spermatozoa. The amplicon regions investigated represent promoter, intragenic, intergenic, possible enhancer and imprinting control regions, with different underlying CG-dinucleotide compositions, and differentially marked by DNA methylation, histone variants and posttranslational modifications as had previously been characterized by population-based MNase-/ChIP-seq experiments. Our experiments conclusively measure low nucleosome frequencies of just a few percent across all amplicons evaluated in both species. The mean percentages at amplicons ranged from ∼1.2 to 1.7% in mouse sperm and from ∼2.3 to 4.5% in human sperm samples from three individual fertile donors. All these samples had been subjected to DNAse I- and FACS-based purification and DTT-NPM treatment which enabled nuclear decondensation and gain of chromatin accessibility in over 70% of cells. These findings are supported by decreasing nucleosome frequencies measured in accessible regions in elongating spermatids, profiled without prior DTT-NPM treatment. We conclude that nucleosomes are unlikely to serve a deterministic regulatory role in paternal transmission of epigenetic information between and across generations of mice and humans.

The long-standing question of nucleosome occupancy in sperm gained relevance in recent years, due to a growing number of studies postulating that sperm borne modified nucleosomes may confer paternal inheritance of environmental cues experienced by the father that would affect health of offspring^62,63^. Moreover, various MNase-seq, ChIP-seq and ATAC-seq based studies reported discordant results regarding nucleosome occupancies in sperm that were likely due to variabilities in sperm purification protocols, methods for opening and enzymatically digesting sperm chromatin, and normalization approaches, also affecting the percentage of informative cells^10,11^. Indeed, we identified by NOMe-seq many inaccessible reads in mouse sperm decondensed with detergents previously used in MNase-seq and ChIP-seq^60^. By decondensing spermatozoa with NPM and performing NOMe-seq, we obtained quantitative measures of nucleosome occupancy in mouse and human male haploid germ cells.

Nonetheless, in line with our and other previous MNase-/ChIP-seq based findings^1–3,55^, we observed in mouse and human spermatozoa an up to 2-fold higher nucleosome occupancy frequency at certain CG-rich amplicons devoid of endogenous CpG methylation, such as promoters, enhancers, maternal ICRs, and CTCF containing amplicons, compared to methylated regions. We measured a comparable difference in elongating spermatids suggesting that endogenous DNA methylation directly or indirectly modulates the extent of nucleosome removal during chromatin remodeling and nuclear elongation. *In vitro*, DNA methylation enhances DNA condensation in the presence of polyamines^64^. The impact on protamines, however, remains unknown. TNP2’s preference for GC-rich DNA^65^ suggests a potential interaction with CpG methylation, but its role in histone replacement is uncharacterized. Functional studies are needed to clarify whether DNA methylation directly affects nucleosome eviction in sperm.

In mice, chromatin of mature sperm is highly compact and uniformly inaccessible to exogenous GpC methyltransferase activity at all regulatory and intergenic amplicons profiled (Fig. 2c). These sites remain inaccessible even upon reduction of disulfide bonds between protamines by DTT treatment (Fig. 3c-e; Ext. Data Fig. 3g). The few residual nucleosomes were only detectable upon NPM-mediated removal of protamines and unfolding of sperm chromatin (Fig. 3c-e; Ext. Data Fig. 3e,g). Hence, these data support the view that chromatin reprogramming in elongating spermatids of mice is remarkably efficient, resulting in a comprehensive and tight packaging of the genome by protamines in mature sperm, at least as measured at promoter proximal and distal regulatory sequences studied here. Like the few residual nucleosomes, any sporadic open chromatin at regulatory sequences is unlikely to serve instructive roles in paternal epigenetic inheritance in mice, e.g. in specifying *de novo* chromatin formation upon fertilization, in contrast to endogenous DNA methylation, as we have previously shown^55^. Along similar lines, we failed to detect footprints for CTCF in elongating spermatids, even in accessible chromatin reads, and in decondensed mature spermatozoa while they were readily measurable at high frequencies in RSts. These data point to efficient removal of CTCF from chromatin during nuclear elongation and a rather limited potential for paternal transmission of such TF-based bookmarking, alike reduced CTCF occupancy during mitosis in differentiated cells^66^. Likely, maternally provided CTCF binds newly to the paternal genome in one-cell embryos^67^. Hence, our findings contrast previous conclusions drawn from ATAC-seq based accessibility studies and warrant further in-depth quantitative SMF analyses^5,34,68^.

In human, chromatin in mature sperm is also highly compacted and uniformly inaccessible to exogenous GpC methyltransferase activity, as in mice (Fig. 4b-c; Ext. Data Fig. 4h-i). However, upon DTT-mediated reduction of protamine disulfide bonds, we observed this time a gain in accessibility at almost all amplicon regions, up to ∼25% of unique reads, resulting in full or partial accessibility and a low frequency of nucleosomes (Fig. 4b-c; Ext. Data Fig. 4h-i,4l-m). Thus, contrary to mice, chromatin repackaging of the human genome during spermiogenesis seems incomplete, leaving many regions devoid of protamines. A comparable finding was recently reported for human sperm pretreated with high concentrations of DTT^69^. These findings concur with ∼18% lower protamine levels quantified in individual spermatozoa of human compared to those of mouse, hamster, bull and horse^70^. Our data argues that many of such regions are nevertheless folded into inaccessible configurations in mature sperm, protected through oxidation of nearby protamines. It remains to be determined whether such inefficient protamine packaging is a general feature of human spermatogenesis resulting in stochastic heterogeneity in genome-wide chromatin states among spermatozoa and/or whether it reflects aberrant genome-wide chromatin remodeling in a subset of spermatozoa. In case of the former option, transmission of open chromatin may have only a probabilistic but not deterministic impact on chromatin formation in the early human embryo. For the latter option, it remains open whether major improperly condensed spermatozoa would support embryogenesis.

Our NOMe-seq data are in line with a recent imaging-based chromatin accessibility study of human sperm, in which fluorescently labeled nucleotides had been enzymatically incorporated in accessible DNA sequences^71^. This study investigated sperm of normozoospermic fertile and infertile men and reported the existence of sperm subpopulations which varied in DNA accessibility that related to varying reproductive outcomes^71^. Aberrantly high accessibility may hinder embryonic development. Another study described varying levels of protamines and redox states of their thiols in spermatozoa of normozoospermic men^61^, indicating heterogeneity in sperm chromatin composition and compaction.

In mice, we also observed cellular heterogeneity, not in the degree of chromatin compaction in mature sperm, but likely in the timing of chromatin remodeling during spermiogenesis. By comparing round spermatids and populations of early and late elongating spermatids, differing in nuclear sizes and the extent of Tnp2 and Protamine1 labeling, we observed a notable loss of nucleosomes and a drastic opening of chromatin followed by a progressive decrease in nuclear accessibility during spermiogenesis. Even though the two populations of elongating spermatids are developmentally assorted, each covering several days of development, our results indicate that while the initiation of remodeling is variable between amplicon regions and individual elongating spermatids, the rate of subsequent remodeling leading to inaccessibility appears comparable between most amplicons. Such heterogeneity may reflect inherent cell-to-cell variability in the nucleosome-to-protamine remodeling process that occurs along the anterior to posterior axis of spermatid heads during their nuclear elongation^72^. In line with our NOMe-seq and ATAC-seq observations, loss of nucleosomes starting early during elongation has also recently been reported in deep ATAC-seq analyses of developmentally synchronized or FACS sorted spermatids^73,74^. Nonetheless, the detection of residual nucleosome-sized fragments later during nuclear elongation may point to a certain level of coordination of timing of remodeling according to genome compartmentalization existing afore in RSts^73^. Hada and colleagues also described chromatin hyper-relaxation and subsequent PRM1 deposition consolidation at gene-rich type A compartments prior to at type-B compartments during spermiogenesis^74^. The quantitative extent of such regulation awaits further analysis. Providing a time frame for bulk nucleosome removal, an independent quantitative imaging study recently showed that replication-dependent H3 histones are effectively removed just prior to stage XII, concurrently with abundant Tnp2 presence^75^. Intriguingly, we observed variably short footprints in accessible regions of elongating spermatids. Since protamines bind directly to DNA, covering ∼11 to 15 bp^70^, we hypothesize that the short footprints may reflect individual and/or small assemblies of nearby TNP and/or protamine proteins that have not yet compacted the chromatin. Alternatively, some may reflect binding of subnucleosomal particles containing the histone variant H2AL2 and TNPs and covering ∼60 bps of DNA^51^. Eventually, larger amounts of protamines will be needed to densely package the chromatin fiber, followed by their oxidation enabling three-dimensional compaction, as noted above for human sperm.

Finally, our study underscores the importance of removing contaminating somatic and/or immature germ cells and of extracellular chromatin originating from damaged and/or lysed nuclei prior to quantitative determination of chromatin states in mouse and human spermatozoa^4^. The quantitative assessment of accessible chromatin states in sperm is particularly sensitive to extracellular contamination as well as to experimental approaches like DTT treatment used to increase accessibility.

## Methods

### Biological sample collection

#### Mice

Animal housing, handling and experimental procedures with animals were performed in accordance with the Swiss Animal Protection Ordinance (licenses 2670 and 3183) and in compliance with FMI institutional guidelines. Mice were housed in Type II long cages containing aspen bedding on IVC racks in rooms with 15–20 air changes per hour and a 12h light/dark cycle. Temperature was maintained within 20–24 °C with a relative humidity within 45–65%. Food and water were provided ad libitum. Mouse spermatids and spermatozoa were obtained from C57BL/6JRj male mice aged 3-7 months.

#### Human sperm samples

Semen samples were provided by healthy donors of known fertility at the Reproductive Medicine and Gynecological Endocrinology (RME) clinic of the University Hospital of Basel. Research was approved by the Ethics Commission of Northwest- and Central Switzerland (EKNZ)^76^ Sperm were washed in Modified Ham’s F-10 (MHF-10 media), centrifuged at 300g for 15 minutes and stored at −80°C until further use.

#### Isolation of mouse spermatids by cell sorting

Mouse round spermatids and early and late elongating spermatids were isolated as described^54^. Testes were dissected into Gey’s balanced salt solution (GBSS) (G9779, Sigma). After removing the tunica albuginea, seminiferous tubules were transferred into clean GBSS and were dispersed apart slightly. To remove interstitial cells, tubules from two testes were transferred into a 15 ml Falcon tube containing 200U/ml Type I Collagenase (LS004196, Worthington Biochemical) and 5 μg/ml DNAse I (DN25, Sigma) in GBSS. After incubating 5 minutes in a shaking water bath at 32°C, tubules were carefully pipetted with a disposable plastic Pasteur pipet and further incubated for an additional 5 minutes. Tubules were allowed to settle at the bottom of the tube and were transferred into a new Falcon with 200U/ml Type I Collagenase, 5 μg/ml DNAse I and 0.025% Trypsin (2505-014, Gibco) in GBSS. To obtain a single cell suspension, the seminiferous tubules were disaggregated by extensive pipetting followed by 10 minutes incubation at the shaking water bath, supplementation with additional 0.025% Trypsin and a second round of pipetting and 10 minutes incubation. FBS was added to inactivate the Trypsin, and cells were transferred to a fresh Falcon tube through a 40 μm sterile CellTrics® filter (04-004-2327, Sysmex). Staining was performed with 22 μg/ml of Hoechst 33342 (H3570, ThermoFisher) and 1.3 μM SYTO-16 (S7578, ThermoFisher) for 1 hour at room temperature protected from light. Cells were centrifuged at 250g and the pellet was resuspended in GBSS containing 3 μM DRAQ7 (DR70250, BioStatus) and 10 μg/ml DNAse I. After filtering through a 30μm sterile CellTrics® filter cells were sorted into PBS with a BD FACSAria^TM^ III Cell Sorter on the basis of DNA content, STYO-16 fluorescence, forward scatter (FSC) and side scatter (SSC) as described^54^.

#### DNase I treatment and isolation of mouse cauda sperm by cell sorting

Mouse cauda epididymides were dissected into a Petri dish and fat patches were removed with forceps and scissors. Each epididymis was carefully squeezed into 100 µl of EmbryoMax® Human Tubal Fluid (HTF) media (MR-070-D, Sigma) with the help of two forceps. Sperm was transferred into an Eppendorf and was incubated at 500 rpm for 1 hour at 37°C. For DNase I treatment, DNase I stock was added into the sperm suspension to a final concentration of 200µg/ml and incubated at room temperature for 10 minutes. DNase I-treated sperm were centrifuged at 6’000g, washed and resuspended in PBS. Cells were stained with 2 μl/ml Hoechst 33342 (H3570, ThermoFisher) shaking at 500 rpm for 1 hour at 25°C and protected from light. To break sperm tails, cells were briefly sonicated with a Brandson Tip digital sonicator with 10% amplitude and 3 cycles of 0.5 second ON / 2 seconds OFF. Sperm were filtered through a 50 μm sterile CellTrics® filter prior to sorting into PBS with a BD FACSAria^TM^ III Cell Sorter. Cells were sorted according to DNA content, forward scatter (FSC) and side scatter (SSC). Sorted cells were aliquoted, flash-frozen and stored at −80°C.

#### DNase I treatment and isolation of human donor sperm by cell sorting

Frozen semen samples were thawed, washed and resuspended in PBS containing 2.5mM MgCl_2_ and 0.5mM CaCl_2_. For DNase I treatment, DNase I stock was added into the sperm suspension to a final concentration of 200µg/ml and incubated at room temperature for 10 minutes. DNase I-treated sperm were centrifuged at 6000g, washed and resuspended in PBS without MgCl_2_ and CaCl_2_. To break sperm tails, the sperm suspension in closed Eppendorf tubes was sonicated using a Diagenode Bioruptor® Pico sonication device with easy mode setting for 25 cycles of 1 second ON / 30 seconds OFF. Cells were stained with 2 μl/ml Vybrant^TM^ DyeCycle^TM^ Ruby (V10309, ThermoFisher) shaking at 500 rpm for 1 hour at 25°C and protected from light. Sperm were filtered through a 50 μm sterile CellTrics® filter prior to sorting into PBS with a SONY MA900 Multi-Application Cell Sorter. Cells were sorted according to DNA content, forward scatter (FSC) and side scatter (SSC). Sorted cells were aliquoted, flash-frozen and stored at −80°C.

#### Quality control of FACS isolated spermatid and sperm populations

Sorted cells were first manually counted using a hemocytometer. Next, the purity of sorted cell populations was evaluated through the morphological assessment of ∼10,000 – 20,000 nuclei fixed with 4% paraformaldehyde in PBS on 10-well slides (631-1371, VWR) and stained with DAPI in VectaShield mounting media (Vector Labs H-1200).

### Sperm decondensation

#### Decondensation of mouse sperm with detergents

Sperm was decondensed using detergents as described^60^. Fresh or frozen mouse sperm was pre-treated with 50 mM DTT at room temperature for 2 hours and supplemented with 100 mM pre-warmed NEM (E3876, Sigma-Aldrich) for 30 minutes. After a PBS wash, 500’000 sperm were resuspended in 10-50 ul of Complete Buffer 1 (CB1: 15 mM Tris-HCl (pH 7.5), 60 mM KCl, 5 mM MgCl_2_, 0.1 mM EGTA, 300 mM Sucrose, 0.5 mM DTT, 0.25% Igepal, 0.5% DOC) and were decondensed at room temperature for 5, 10 or 30 minutes.

#### Decondensation of mouse sperm with heparin

Mouse fresh sperm was decondensed by incubation with 10 mM DTT, 0.1% Igepal and 1 mg/ml heparin in PBS for 22 minutes at 15°C.

#### NPM purification

The plasmid pNP029-pET24a-6His-SMT3-xNPM149 encoding amino acids 1-149 of *Xenopus laevis* NPM was introduced into the E. coli strain BL21 (DE3)-CodonPlus-RIL (230245, Agilent). Cells were grown in LB with kanamycin (50 µg/ml) and chloramphenicol (35 µg/ml) at 37°C and allowed to grow until an OD_600_ of 0.8. 200 μM IPTG was added and after ∼30 minutes the temperature was decreased to 20°C to induce protein expression. Cells were collected after 20h and resuspended in His-Lysis buffer (50 mM Na-phosphate pH 7.4, 150 mM NaCl, 10 mM imidazole) at 20 ml per 1L culture, and snap-frozen in liquid nitrogen. Thawed cell lysates were topped up with His-Lysis buffer to 50ml per 1L culture, supplemented with 1x cOmplete™ EDTA-free protease inhibitors (11873580001, Sigma), lysozyme (1 mg/ml) (L6876-5G, Sigma), MgCl_2_ (2.5 mM) and PierceTM Universal Nuclease (10 µl per 50 ml). Lysates were homogenized and lysed by sonication, followed by centrifugation (30’000g, 4°C, 30 minutes) and filtering (0.22 µm), and were subsequently loaded onto a HisTrap^TM^ HP 5 ml column (GE Healthcare) prepacked with Ni Sepharose®. After washing the column with His-Wash buffer (50 mM Na-phosphate pH 7.4, 500 mM NaCl, 50 mM imidazole), recombinant NPM with His-SUMO-tag was eluted with His-Elution buffer (50 mM Na-phosphate pH 7.4, 500 mM NaCl, 250 mM imidazole). Peak eluent fractions were pooled and diluted to 3X with 50 mM Na-phosphate pH 7.4 and incubated with ULP1 SUMO-protease for 1 hour at 4°C to liberate the His-SUMO tag. The resultant digestant was further diluted with 2X eluent volumes of 50mM Na-phosphate pH 7.4, filtered (0.22 µm), and applied to the HisTrap^TM^ HP 5 ml column to retain the His-SUMO tag. The flow-through was subsequently loaded onto a HiTrap® Q HP 5 ml column (GE Healthcare), and the column was developed with a linear gradient of NaCl (100-820 mM) and constant Tris (20mM pH8.0). Peak factions were pooled and dialyzed against KH buffer (20 mM HEPES-KOH pH7.7, 100 mM KCl). The final dialysate was concentrated with a Vivaspin® Turbo 4 10 kDa (Sartorius) concentrator tube.

#### Decondensation of mouse sperm with recombinant Xenopus NPM

Mouse sperm was decondensed *in vitro* with recombinant NPM as described^8^ with several modifications. Frozen sperm pellets were thawed and resuspended in 100µl of Lysis Buffer (10mM Tris pH7.4, 10mM NaCl, 3mM MgCl_2_, 0.1mM EDTA, 0.5% Igepal) per aliquot, and cells were incubated on ice for 20 minutes. After spinning at 6’000g and 4°C for 5 minutes, cells were washed with 200µl cold KH buffer (100 mM KCl, 20 mM Hepes-KOH, pH 7.7) and incubated in 100µl KH buffer with or without 2 mM DTT for 15 minutes at 37°C. Sperm were centrifuged at 6’000g, 4°C for 5 minutes and 85µl supernatant was discarded. The remaining 15µl containing the sperm pellet was mixed well, flash-frozen and stored at −20°C for at least 2 hours. After thawing, NPM-decondensation was performed in 25 µl reactions containing 500’000 sperm, 200 μM of purified NPM, 20 mM Hepes-KOH pH 7.7, 100 mM KCl, 2.5 mM MgCl_2_, 5 mM Sodium Butyrate, 10 mM DTT and 1x cOmplete™ EDTA-free protease inhibitors (11873580001, Sigma). Cells were incubated at 37°C with 600 rpm for the indicated time.

#### Decondensation of human sperm with recombinant Xenopus NPM

Human sperm was decondensed in vitro with recombinant NPM as described^8^ with several modifications. Frozen sperm pellets were thawed and resuspended in 100µl of Lysis Buffer (10mM Tris pH7.4, 10mM NaCl,3mM MgCl_2_, 0.1mM EDTA, 0.5% Igepal) per aliquot, and cells were incubated on ice for 20 minutes. After spinning at 6’000g and 4°C for 5 minutes, cells were washed with 200µl cold PBS and incubated with 2 mM DTT in 50µl PBS for 15 minutes at 37°C. Reduced spermatozoa were centrifuged at 6’000g, 4°C for 5 minutes and washed once with 100µl KH buffer (100 mM KCl, 20 mM Hepes-KOH, pH 7.7). NPM-decondensation was performed in 25 µl reactions containing 500’000 sperm, 100 μM of purified NPM, 20 mM Hepes-KOH pH 7.7, 100 mM KCl, 2.5 mM MgCl_2_, 5 mM Sodium Butyrate, 10 mM DTT and 1x cOmplete™ EDTA-free protease inhibitors (11873580001, Sigma). Cells were incubated at 37°C with 600 rpm for the indicated time.

### Design, selection and validation of NOMe-seq primers

#### Defining NOMe-seq amplicon categories

The following 26 categories of amplicons were defined based on genomic location (“Promoter” vs. “Intergenic”), CpG density (“High CG”: ≥ 0.5 observed/expected CG; “Medium CG”: <0.5 and ≥ 0.25 observed/expected CG, and “Low CG”: <0.25 observed/expected CG), containing a CpG Island “CGI” or not “nonCGI”, DNA methylation level (“Hypomethylated”: DNA methylation ≤ 25%; “Hypermethylated”: DNA methylation > 75%) as well as chromatin status and function (i.e. overlapping an enhancer, a CTCF binding site or an imprinted control region (ICR)): 1) Promoter, Hypomethylated, High CG; 2) Promoter, Hypomethylated, Medium CG; 3) Promoter, Hypomethylated, Low CG; 4) Promoter, Hypermethylated, High CG; 5) Promoter, Hypermethylated, Medium CG; 6) Promoter, Hypermethylated, Low CG; 7) Intragenic, Hypomethylated, High CG; 8) Intragenic, Hypomethylated, Medium CG; 9) Intragenic, Hypermethylated, High CG; 10) Intragenic, Hypermethylated, Medium CG; 11) Intragenic, Hypermethylated, Low CG; 12) Intergenic, Hypomethylated, High CG; 13) Intergenic, Hypomethylated, Medium CG; 14) Intergenic, Hypomethylated, Low CG; 15) Intergenic, Hypermethylated, High CG; 16) Intergenic, Hypermethylated, Medium CG; 17) Intergenic, Hypermethylated, Low CG; 18) Enhancer, Hypomethylated, High CG; 19) Enhancer, Hypomethylated, Medium CG; 20) Enhancer, Hypomethylated, Low CG; 21) Enhancer, Hypermethylated, High CG; 22) Enhancer, Hypermethylated, Medium CG; 23) Enhancer, Hypermethylated, Low CG; 24) Maternal ICR (Hypomethylated); 25) Paternal ICR (Hypermethylated); 26) CTCF (Hypomethylated, High CG). For mouse, primers were designed for categories 1, 2, 6, 12-19, and 24-26. Human amplicons were generated for categories 1-25.

#### Generation of input coordinates for primer design

Gene annotations were retrieved from *TxDb.Mmusculus.UCSC.mm10.knownGene* and *TxDb.Hsapiens.UCSC.hg38.knownGene* for mouse and human, respectively. Promoter coordinates were defined as TSS +/- 500 bp of reference genes. Coordinates of intergenic regions were generated by tiling mouse (BSgenome.Mmusculus.UCSC.mm10) or human (BSgenome.Hsapiens.UCSC.hg38) genome assemblies into 1 kbp sequences and discarding tiles overlapping gene coordinates, promoters of all annotated transcripts (+/- 2.5 kbp from the TSS), 1kbp downstream of the 3’ end of genes, enhancers and imprinted control regions (ICRs) (Fig. 1a). Mouse and human enhancer coordinates were downloaded from the Fantom5 project (https://fantom.gsc.riken.jp/5/datafiles/latest/extra/Enhancers/; *“mouse_permissive_enhancers_phase_1_and_2”*and *“human_permissive_enhancers_phase_1_and_2”)*. Enhancers predicted for mouse testes^77^ were also included (http://chromosome.sdsc.edu/mouse/download.html, *“testes.enhancer.txt”,* resized to 1kbp) and amplicons overlapping both datasets were preferentially selected. Coordinates for CGI (“*cpgIslandExt*”) and repeat annotations (RepeatMasker, *“rmsk”*) were downloaded from UCSC Table Browser and mouse ICR coordinates were taken from^78^. CTCF binding and DNA methylation status were determined after primer design (see below). Blacklisted regions from ENCODE (accessions ENCFF547MET and ENCFF001TDO) and^79^ were filtered out. LiftOver was used to convert coordinates between genome assemblies. Genomic coordinates were assigned to amplicon categories with the following order of priority: CTCF > ICR > Promoter > Enhancer > Intragenic > Intergenic.

#### Primer design

NOMe-seq primers for each amplicon category were designed with in house scripts wrapping Primer3 with minor modifications and were kindly provided by the Schübeler group^57^. Genomic sequences for primer hybridization lacked CpG and GpC dinucleotides. Amplicon size was set between 450 bp and 580 bp and Tm between 50°C and 60°C.

#### Amplicon filtering, characterization and selection

To ensure an optimal distribution of GCH trinucleotides (GCH) for high resolution nucleosome footprinting, amplicons with >40 bp separating any consecutive GCH were discarded. Amplicons with less than 17 GCH and more than 65 GCH were also removed. Each primer pair was required to contain at least 3 cytosines to ensure specificity for bisulfite converted DNA. Amplicons with the smallest gaps between GCHs and the maximum length covered by GCHs were preferentially selected. When possible, amplicons were further filtered by minimum and maximum Tm of 54°C and 57.5°C respectively. Amplicons overlapping repeats were avoided. For certain amplicon categories in mouse or human genomes, none of the few primer pairs obtained passed filtering criteria and those categories were left out. Promoters of genes with known function were preferentially selected. Gene names were obtained with AnnotationDbi R package^80^.

Average CpG methylation levels of filtered amplicons were quantified with QuasR^81^ using published whole genome bisulfite sequencing data of mouse and human spermatozoa^52^. Amplicons were accordingly classified as hypomethylated (≤ 25% CpG methylation) or hypermethylated (>75%). Amplicons with less than 10 reads were discarded. Yet, a lower number of reads was accepted for categories with scarce numbers of filtered amplicons. Several ATAC-seq, MNase-seq and ChIP-seq published were downloaded, processed and quantified at amplicon coordinates as described below. CTCF amplicons were selected based on high CTCF ChIP-seq enrichment in round spermatids^53^ and/or in sperm^5^. Amplicons overlapping putative enhancers were further subselected on the basis of high levels of H3K4me1 and H3K27ac in sperm. Those with lower levels of H3K4me1 and H3K27ac were reclassified as intergenic.

#### Lambda phage DNA, other control amplicons and primer validation

Amplicons to amplify spiked in Lambda phage DNA were similarly designed from NC_001416.1 NCBI sequence. Additional shorter amplicons were also included as internal control between mouse experiments. Longer distances between consecutive GCHs were allowed for control and lambda DNA amplicons. Selected primers (Supplementary Tables 1 & 2) were commercially synthesized, and amplification products were validated by agarose gel electrophoresis.

### Chromatin methods

#### Standard procedure of Nucleosome Occupancy and Methylome amplicon sequencing (NOME-seq)

NOMe-seq was performed as described^12,82^ with minor modifications. 500’000 haploid cells or 250’000 diploid cells were permeabilized with 250µl ice-cold Lysis Buffer (10 mM Tris pH 7.4, 10 mM NaCl, 3 mM MgCl_2_, 0.1 mM EDTA, 0.5% Igepal) for 10 minutes on ice. 750 µl of Wash buffer (10 mM Tris pH 7.4, 10 mM NaCl, 3 mM MgCl_2_, 0.1 mM EDTA) was added and nuclei were centrifuged for 5 minutes at 4°C. Cultured cells were centrifuged at 600g, round spermatids at 1’600g, elongating spermatids at 3’000g, mouse and human sperm at 6’000g unless indicated otherwise. Nuclei were resuspended in 250 µl of Wash Buffer, centrifuged and further resuspended in 94.5 µl of 1x GC Reaction Buffer (M0227L, NEB). Footprinting was performed with the addition of 150U of M.CviPI (M0227L, NEB) in a final concentration of 1 mM S-adenosylmethionine (SAM) (M0227L, NEB) and 300 mM sucrose. 2.6 ng of lambda DNA (SD0011, Thermofisher) was spiked in per reaction. After 15 minutes of incubation at 37°C, the reaction was stopped with 10 mM Tris pH 7.4, 200 mM NaCl, 5 mM EDTA, 0.5% SDS and 200 ug/ml Proteinase K. After an overnight incubation at 55°C and 1’200 rpm, samples were RNAse A treated and DNA was purified with Phenol:Chloroform:Isoamyl Alcohol (15593031, ThermoFisher) and Ethanol precipitation. DNA pellets were resuspended in H_2_O and quantified with Qubit 1x dsDNA High Sensitivity kit (Invitrogen). 600 ng to 1’400 ng of DNA per reaction were typically recovered.

DNA was bisulfite converted with EpiTect Bisulfite Kit (59104, QIAGEN) according to the standard protocol. Amplicons were individually amplified using KAPA HiFi HotStart Uracil+ ReadyMix Kit (07959079001, Roche) in a total volume of 12.5 µl with 0.4 mM primers and 15-30 ng of input DNA. Different annealing temperatures ranging from 54°C to 60°C were used for groups of amplicons with similar Tm. After an initial denaturation at 98°C for 5 minutes, samples were amplified with 35 cycles at 98°C for 30 seconds, 54-60°C for 30 seconds, and 72°C for 30 seconds. According to amplification yield, 2 µl, 4 µl or 8 µl of each amplicon were pooled per sample. PCRs pools were purified using 0.8X AMPureXP beads (A63881, Beckman Coulter) and quantified with Qubit 1x dsDNA High Sensitivity kit. Libraries were prepared from 100-300 ng of amplified DNA with NEBNext® Ultra™ II DNA Library Prep Kit (E7645S, NEB) and NEBNext Multiplex Oligos (E7335, E7500, E7710, E7730, NEB) according to kit instructions. Quality of libraries was assessed with Fragment Analyzer (Agilent) and sequenced paired-end 300×300 bp with an Illumina MiSeq sequencer.

Three independent biological replicates were performed for round spermatids, early and late elongating spermatids and mouse sperm. Each replicate was obtained by pooling sorted cells from 2 mice. A negative control sample was generated from mouse sperm with a mock NOMe-seq reaction lacking M.CviPI.

#### NOMe-seq of NPM-decondensed mouse and human sperm

At indicated timepoints, 500’000 mouse or human sperm incubating in 25µl of NPM decondensation reaction (see above) were complemented with 69.5µl 1x GC Reaction Buffer and footprinted for 15 minutes at 37°C with 150U of M.CviPI (M0227L, NEB), 2.6 ng of lambda DNA (SD0011, Thermofisher), a final concentration of 1 mM SAM (M0227L, NEB) and 300 mM sucrose. Samples were further processed as described in *Standard procedure.* Two independent biological replicates were performed for mouse sperm. NOMe-seq for each timepoint of decondensation was carried out independently for sperm samples from mice and human donors A, B and C.

#### NOMe-seq of samples decondensed with detergents

Decondensation reactions performed in small volumes (<20 µl) were directly resuspended in 1x GC Reaction Buffer to a total volume of 94.5 µl. Alternatively, CB1 buffer without detergents was added to the decondensation reaction prior to spinning at 6’000 – 10’000g for 5 minutes in at 4°C. Supernatant was removed, cell pellets were resuspended in 94.5 µl of 1x GC Reaction Buffer and footprinting was performed as described above in *Standard procedure*.

#### Assay for transposase-accessible chromatin (ATAC-seq)

ATAC-seq was performed as previously described^83^ with minor modifications. Cells in PBS were counted with a hemocytometer and 50’000 diploid or 100’000 haploid cells were aliquoted per reaction. After spinning for 5 minutes at 4°C cell pellets were lysed in 50 µl of 10mM Tris-HCl, pH 7.4, 10mM NaCl, 3mM MgCl_2_ and 0.5% Igepal (CA-630, Sigma) for 5 minutes on ice. 5 volumes of lysis buffer without detergent were added and cells were centrifuged for 10 minutes at 4°C. The supernatant was carefully removed and nuclei were resuspended in tagmentation reaction mix containing 25 µl of TD buffer (FC-121-1030, Illumina Nextera Kit), 2.5 µl of transposase (TDE1, FC-121-1030, Illumina Nextera Kit) and 22.5 µl of nuclease-free water. Nuclei were tagmented for 30 minutes at 37°C and 600 rpm. DNA was immediately purified with Qiagen MinElute PCR purification Kit (28004, QIAGEN) and stored at −20°C. Libraries were PCR amplified with Q5 High-Fidelity DNA Polymerase (M0541, New England Biolabs) and 0.5 µM indexed primers as designed^83^. The following cycling conditions were used: 72°C for 5 minutes; 98°C for 30 sec; and 10–11 cycles at 98°C for 10 sec, 63°C for 30 sec, and 72°C for 1 minute. The optimal number of PCR cycles were estimated by qPCR. Libraries were purified with AmpureXP beads (A63881, Beckman Coulter) and sequenced paired-end with a NextSeq500 instrument (Illumina)^60,84^.

### NOMe-seq data quantification and statistical analysis

#### Data pre-processing and alignment

NOMe-seq raw sequencing reads were pre-processed with Trim Galore^85^ to remove Illumina adaptors and low quality bases (parameters: --paired --illumina --quality 20 --length 125 --clip_R1 5 --clip_R2 5). Trimmed reads were aligned with QuasR (Bowtie aligner, parameters: “-X 700 -k 2 --best --strata”, bisulfite mode “undir”)^81^ to a custom genome containing amplicon genomic sequences retrieved from BSgenome.Mmusculus.UCSC.mm10, BSgenome.Hsapiens.UCSC.hg19 and NC_001416.1 for mouse, human and Lambda phage genomes, respectively.

#### Methylation calling and filtering

Cytosine methylation calling and data retrieval from BAM files were performed on individual sequenced fragments (i.e. single DNA molecule) with R package *fetchNOMe*^13^ by analysing C to T and G to A substitutions resulted from bisulfite conversion. Cytosines were split into the following groups based on genomic context:

1. Cytosines in “GCH” context were used to retrieve information about footprints and included GCA, GCC and GCT.
2. Cytosines in “WCG” context were used to retrieve information about endogenous CpG methylation and included ACG and TCG.
3. Cytosines in “bisC” context were used for filtering fragments with failed bisulfite conversion and included ACA, ACC, ACT, TCA, TCC and TCT.

Fragments with >10% “bisC” methylation were considered as incompletely bisulfite converted and were filtered out. A minimum of 10 “bisC” positions was required to enter evaluation of bisulfite conversion efficiency. Fragments with the same start and end position and identical methylation states of all cytosines were considered PCR duplicates and only one unique fragment was retained for further analysis. Only amplicons with >=100 filtered reads were kept for analysis.

#### Single-molecule heatmaps of methylation status

R functions were used to generate single-molecule WCG or GCH heatmaps by hierarchically clustering individual sequenced fragments based on WCG or GCH methylation, respectively. For improved visualization, dendrograms were further reordered using as weights the percentage of WCG or GCH methylation of each read but maintaining the constraints of the dendrogram.

#### Prediction of footprints and classification of NOMe-seq molecules

Quantification of footprint abundances in NOMe-seq data has been performed using R package “nomeR” (version 0.7.0)^13^ that provides functionality for i) footprint spectral analysis −inference of representative footprint lengths in NOMe-seq datasets and ii) footprint prediction −determining positions of footprints with different lengths in individual NOMe-seq molecules. Briefly, the footprint prediction and classification of NOMe-seq fragment were performed following these three steps.

i. ***Footprint spectral analysis:*** Footprint spectral analysis for spermatogenesis data has been done according to the following procedure. First, we merged biological replicates for each amplicon and cell type. Second, we noticed that partial footprints at edges of amplicons introduce spurious peaks in footprint spectra. Therefore, we clipped all protected GCHs at edges of amplicons and only focused on footprints which are fully contained within amplicons. Third, we performed footprint spectral analysis using function *infer_footprints_stan_vb* from the nomeR package using non-informative prior distribution for footprint abundances. To obtain prior hyperparameters for noise levels we use control NOMe-seq data for spike-in lambda DNA and fit Beta-binomial distribution to observed frequencies of number of protected GCHs per fragment. Resulting shape parameters of Beta-binomial distribution we use as hyperparameters for background noise prior distribution. Similarly, for footprint noise prior distribution, we use control NOMe-seq data for lambda DNA without M.CviPI for fitting Beta-binomial distribution and obtaining hyperparameters for footprint noise prior distribution.
ii. ***Prediction of footprint locations:*** To determine exact locations of nucleosome footprints within single molecules we used the function *predict_footprints* from the nomeR package. To construct footprint models needed for predicting footprints we selected ranges in footprint spectra obtained by footprint spectral analysis based on typical peaks observed for the data. The following footprint ranges were chosen for construction of footprint models:

- 2 - 5 bp (“Short”) correspond to short footprints and presumably reflect correlated background noise for which the assumption of statistical independence between position is broken.
- 30 - 80 bp (“TF”) corresponds to typical footprints from CTCF binding.
- 100 – 150 bp (“Nucleosome”) corresponds to typical nucleosome footprints.
- 151 – 200 bp (“Inaccessibility”) aims to identify fragments which are fully inaccessible. After determining prior coverages we collapsed the “Inaccessibility” footprint models into only a single model of length 200. After running prediction for all footprint models and each fragment using the function *predict_footprints* from the nomeR package, we aggregated coverage posterior probability profiles for all footprint models with lengths belonging to each footprint range, e.g. “Short”, “TF”, etc. Given calculated coverage profiles for “Nucleosome” and “Inaccessibility” for each position in each fragment we choose a set of cutoffs to assign NOMe-seq fragments to several footprint classes.
iii. ***Classification of NOMe-seq fragments:*** Based on coverage posterior probabilities for individual fragments, one of the following 4 categories was correspondingly assigned: “Nucleosome”, “Fully Accessible”, “Fully Inaccessible” or “Partially Accessible”. Classification was performed as follows: fragments with >= 90% of inaccessible GCHs and/or predicted mean coverage for inaccessibility >= 0.5 were classified as “Fully Inaccessible”. Fragments below the thresholds were further quested for accessibility. When predicted mean coverage for accessibility was >=0.9, class “Fully Accessible” was assigned. If <0.9, fragments were classified as “Nucleosome” when maximum predicted coverage for nucleosome was >= 0.8. Fragments not categorized as: “Nucleosome”, “Fully Accessible” or “Fully Inaccessible” were assigned to “Partially Accessible”. For simplification and because the current study mostly focused on the analysis of nucleosome occupancy, fragments containing TF footprints (30 - 80 bp) but devoid of nucleosomes were also grouped into “Partially Accessible”.

To select the cutoffs above, multiple thresholds were evaluated beforehand for optimal footprint classification. The criteria followed for cutoff selection was: i) Inaccessibility: >99.9% of fragments from an unfootprinted sample (i.e. negative control not treated with M.CviPI methyltransferase) were classified as Inaccessible (data not shown). ii) Accessibility: chosen threshold allowed to exclude most TF footprints as well as highly accessible reads with short footprints originating from elongating spermatids. iii) Nucleosome: the percentage of fragments categorized as Nucleosomes did not change significantly between tested thresholds.

#### ATAC-seq data processing

Published ATAC-Seq datasets were downloaded from NCBI GEO repository. Nextera adapters were trimmed using Trim Galore with parameters “--nextera --paired --stringency 3”. Reads were aligned to mouse (BSgenome.Mmusculus.UCSC.mm10) or human (BSgenome.Hsapiens.UCSC.hg19) genome assemblies with QuasR (Bowtie aligner, parameters: “-300 maxHits --best –strata --maxins 1500”) prior removal of the last base with the same package^81^. Potential PCR duplicates were discarded using samtools^86^.

#### MNase-seq and ChIP-seq data processing

Adaptors of published and custom MNase-seq and ChIP-seq datasets were removed with Trim Galore^85^ with parameters “--illumina. --stringency 3”, and “—paired” when processing paired-end data. Processed reads were aligned to mouse (BSgenome.Mmusculus.UCSC.mm10) or human (BSgenome.Hsapiens.UCSC.hg19) genome assemblies using STAR with parameters “--alignIntronMin 1 --alignIntronMax 1 --alignEndsType EndToEnd --alignMatesGapMax 1000 --outFilterMatchNminOverLread 0.85 --outFilterMultimapNmax 300 --outMultimapperOrder Random --outSAMmultNmax 1)”. Samtools^86^ were used to remove putative PCR duplicates.

#### ATAC-seq, MNase-seq and RNA-seq data quantification

Enrichments were quantified with the qCount function of QuasR^81^. Log2 RPKMs were calculated and input datasets were used for normalization when available. To quantify genome accessibility with paired-end ATAC-seq data only subnucleosome-sized reads (30-100 bp) were selected and counted at read start position.

RNA-seq gene-level read counts were obtained from a previous study^87^ and processed with the same framework using QuasR. Biological replicates were merged by summing raw counts per cell type. Expression levels were normalized to Reads Per Kilobase per Million mapped reads (RPKM), followed by log2 transformation with a pseudocount of 1. For each cell type, genes were classified into three expression categories based on the distribution of log2(RPKM + 1) values: 0–25%, 25–75%, and 75–100%, reflecting relative expression within each sample.

## Data availability

The NGS datasets used in this study are available from the Gene Expression Omnibus (GEO) database.

## Code availability

Computational codes and scripts used in this study for fast retrieval of NOMe-seq data from BAM files (fetchNOMe) are available at https://doi.org/10.5281/zenodo.8402785 and https://github.com/fmi-basel/compbio-nomeR.

## Acknowledgements

We gratefully acknowledge Dr. Yuki Okada for providing advice on NPM chromatin decondensation methology. We thank staff of the RME for helping in preparation of human semen samples. We thank FMI colleagues Hubertus Kohssler (cell sorting), Dr. Sirisha Aluri, Eliza Pandini Figueiredo Moreno, Dr. Sebastien Smallwood (Functional Genomics), Sandra Muehlhaeusser (Structural Biology) for their excellent support. We thank Dr. Nhuong Nguyen and other members of the Peters laboratory for critical support for the project. This research was supported by the Novartis Research Foundation, the Swiss National Science Foundation (31003A-172873, 320030-189264) and the European Research Council (ERC) under the European Union’s Horizon 2020 research and innovation program (grant agreement ERC-AdG 695288 – Totipotency).

## Author contributions

L.G.-T., H.S. and A.H.F.M.P. designed experiments. L.G.-T. designed, developed, performed and interpreted NPM-driven decondensation experiments of human sperm and amplicon-based NOMe-seq experiments of mouse spermatid and sperm samples. H.S. produced recombinant NPM, further optimized NPM decondensation protocols for mouse and human sperm, developed DNAse I digestion and FACS procedures and performed and interpreted NOMe-seq experiments for mouse and human sperm. E.A.O conceived the statistical SMF analysis model, developed R packages *fetchNOMe* and *nomeR*, and interpreted NOMe-seq data. M.E.G. co-developed and supported FACS sorting experiments. C.D.G. organized human samples. L.G.-T., H.S. and A.H.F.M.P. wrote the manuscript with input from all authors. A.H.F.M.P. conceived and supervised the overall project and acquired funding.

## Competing interests

The authors declare no competing interests.

## Notes

### Competing Interest Statement

The authors have declared no competing interest.

