## Supplemental Figures and Tables for "Single molecule footprinting measures low nucleosome occupancy in mature spermatozoa of mice and men"

**Extended Data Figures & Tables**

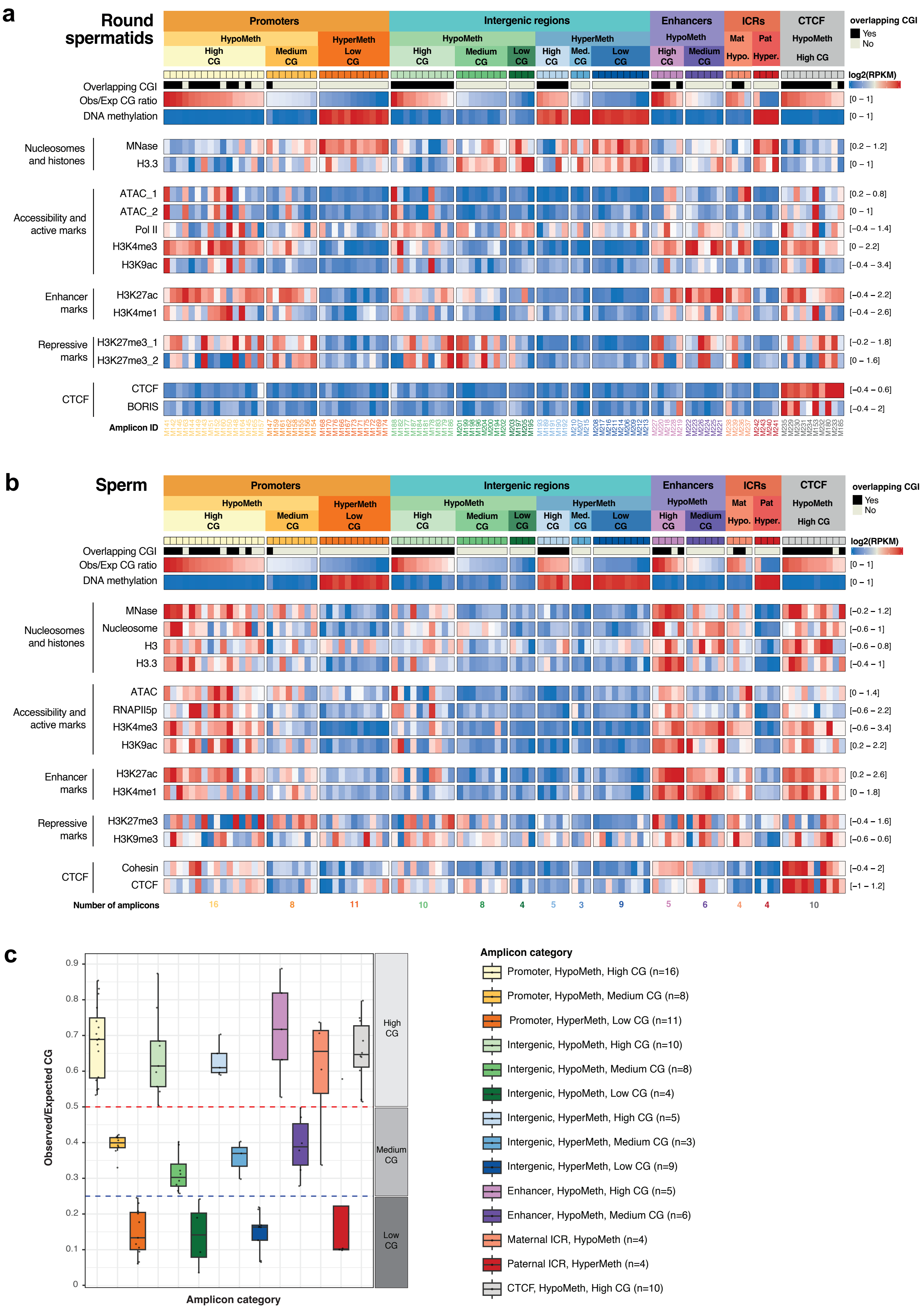


### Extended Data Figure 1

d

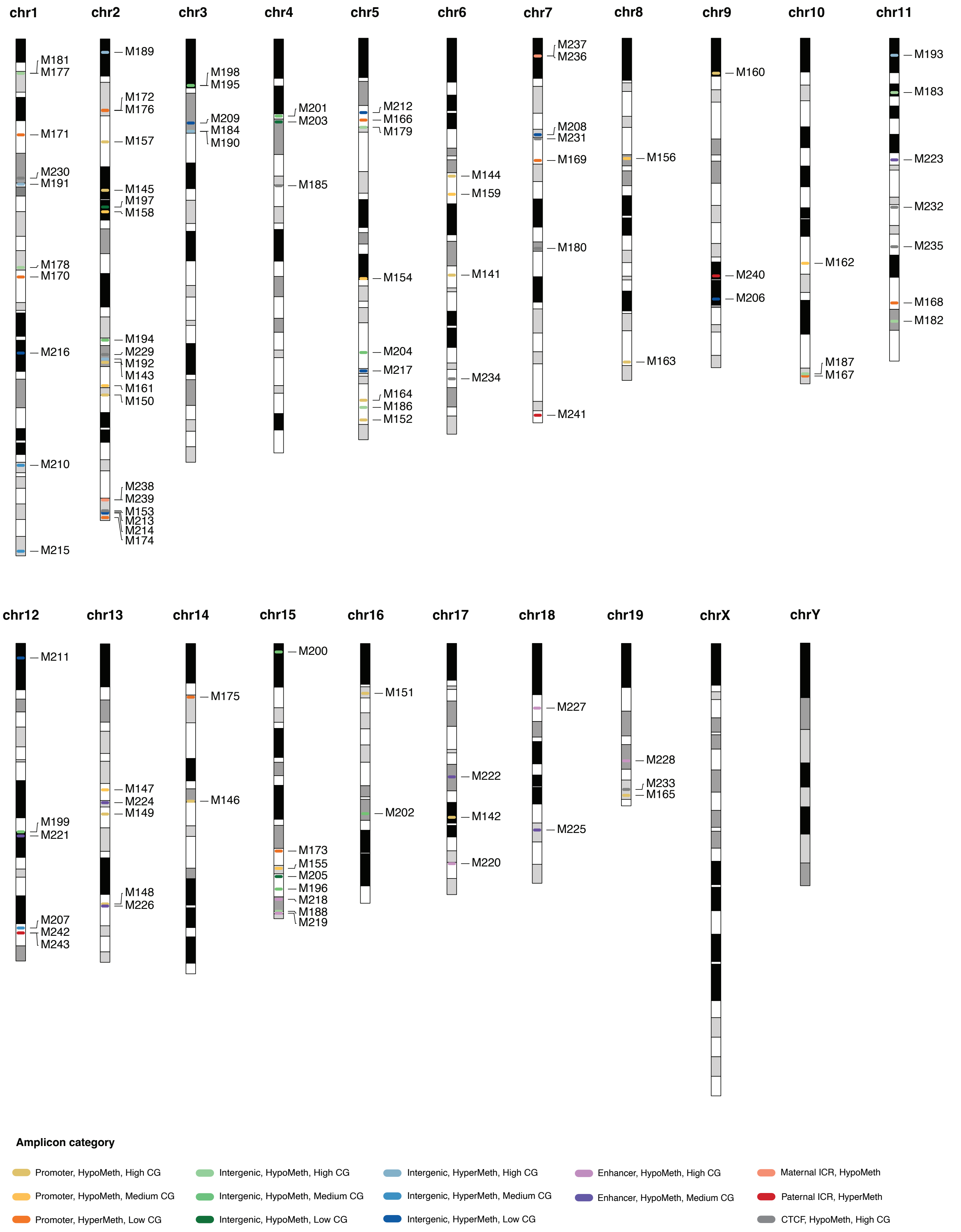


**Extended Data Figure 1 (cont.)**

**e**

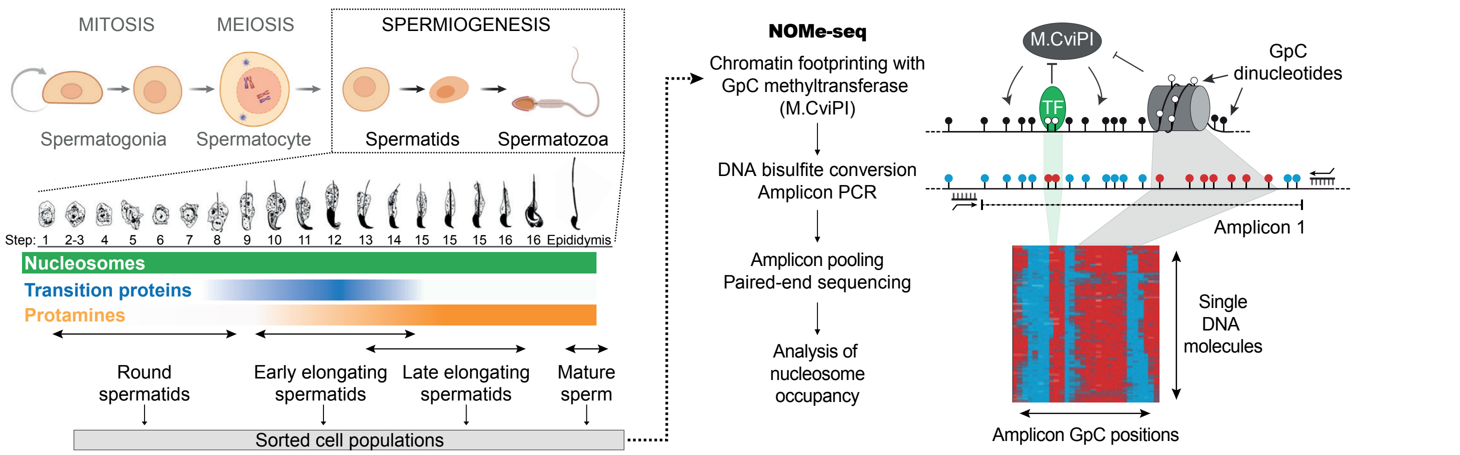


**f**

| Sorted cell population | Replica | Total cell number | % RSt | % ESt | % cauda sperm | % other | Average purity (%) |
|---|---|---|---|---|---|---|---|
| RSt | Rep 1 | 1450 | **98.55** | 1.24 | 0.00 | 0.21 | **98.33** |
| | Rep 2 | 2335 | **98.12** | 1.20 | 0.00 | 0.68 | |
| | Rep 3 | 473 | **98.31** | 1.69 | 0.00 | 0.00 | |
| EESt | Rep 1 | 2393 | 0.08 | **99.79** | 0.00 | 0.13 | **99.49** |
| | Rep 2 | 2955 | 0.24 | **99.46** | 0.00 | 0.30 | |
| | Rep 3 | 389 | 0.51 | **99.23** | 0.00 | 0.26 | |
| LESt | Rep 1 | 951 | 0.11 | **98.95** | 0.00 | 0.94 | **99.47** |
| | Rep 2 | 2054 | 0.05 | **99.46** | 0.00 | 0.49 | |
| | Rep 3 | 260 | 0.00 | **100.00** | 0.00 | 0.00 | |
| Sperm | Rep 1 | 1303 | 0.00 | 0.00 | **100.00** | 0.00 | **99.98** |
| | Rep 2 | 1618 | 0.00 | 0.00 | **99.94** | 0.06 | |
| | Rep 3 | 242 | 0.00 | 0.00 | **100.00** | 0.00 | |

**i**

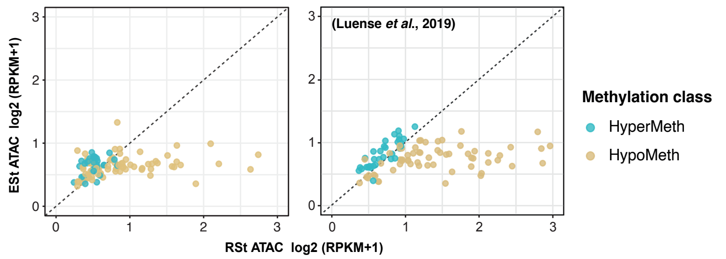


**j**

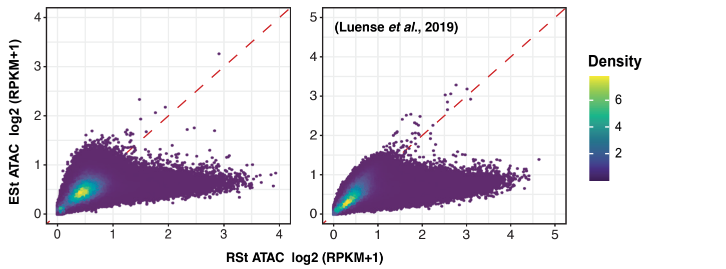


**g**

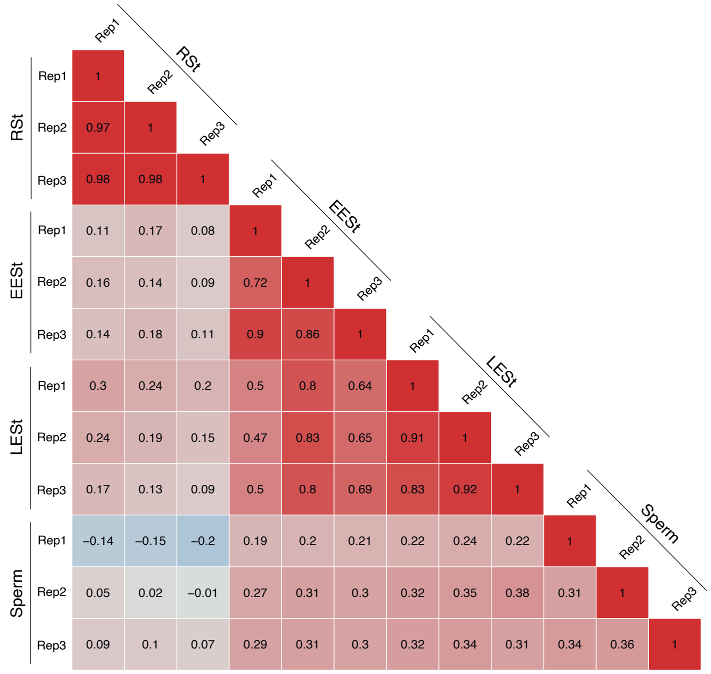


**h**

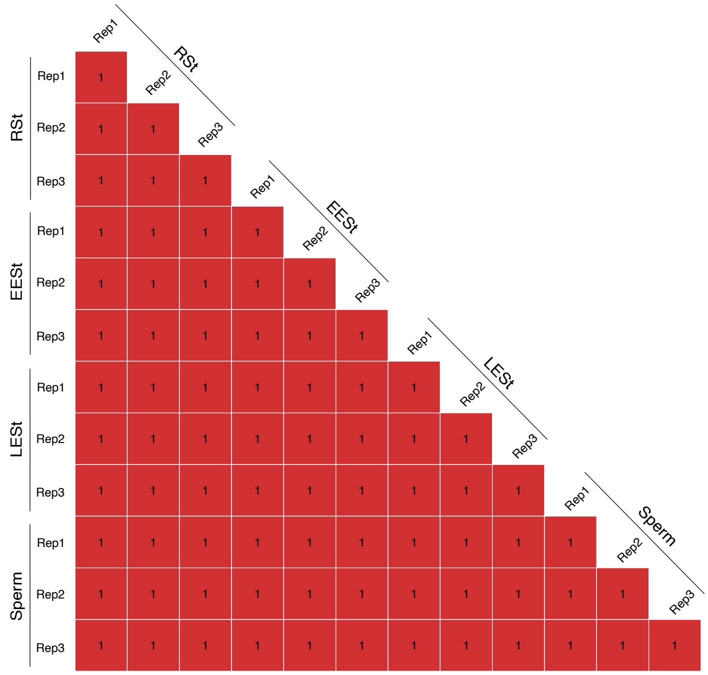


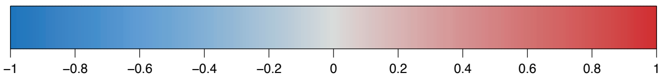

**Extended Data Figure 1 (cont.)**

**Extended Data Fig. 1: Experimental conditions for SMF profiling of male germ cells of mice.**

**a.** and **b.** Scheme showing chromatin characteristics at amplicons that had previously been measured by MNase-/ChIP-seq in RSts and sperm as follows: DNA methylation in RSts and sperm, Pol II, H3K9ac, H3K27ac, H3K4me1, and H3K27me3_1 in RSts[1]; MNase in RSts[2]; MNase in sperm, H3.3 in RSts and sperm, H3K4me3 and H3K27me3_2 in RSts[3]; CTCF and BORIS in RSts[4]; H3K9me3 in sperm[5]; RNAPII5p, H3K9ac, H3K27ac, H3K4me1, Cohesin, and CTCF in sperm; ATAC in RSts and sperm[6,7]; ATAC in RSts, ESts and sperm (this study). Amplicon IDs are shown at the bottom of the plot. Most CG-high regions overlap with CpG islands (CGIs). In RSts, nucleosome and histone occupancies generally correlated inversely to Observed/Expected (Obs/Exp) CG ratio, which has been attributed to differential nucleosome turnover at non-methylated CG-high sites versus methylated CG-low sites[3]. In sperm, however, nucleosome and histone occupancies were relatively enriched at unmethylated CG sites suggesting selective nucleosome retention during sperm development[3,5]. Many CG-high and several CG-medium promoter, intergenic, enhancer and maternal ICR regions showed also enrichment for activity-related post-translational histone modifications such as H3K4me3, H3K9ac, H3K27ac and even H3K4me1 in RSts and/or sperm. Repressive H3K27me3 and H3K9me3 modifications were enriched at certain selected CG-high and CG-medium regions while CTCF showed enrichment at CTCF-motif containing sequences in RSts and sperm.

**c.** Box plot showing Observed/Expected CG ratios of mouse amplicons per amplicon category as defined in panels a/b. Horizontal dashed lines indicate thresholds used to categorize amplicons other than ICRs into High, Medium and Low CG classes.

**d.** Karyogram showing the positions of selected amplicons on mouse chromosomes, colored according to their respective amplicon categories.

**e.** Nucleosome occupancy data has been generated at defined genomic locations (amplicons) in sorted populations of mouse round spermatids, early and late elongating spermatids and cauda epididymal sperm by chromatin footprinting with the M.CviPI GpC methyltransferase (NOMe-seq). GpC dinucleotides are represented as lollipops colored in black (methylated), white (unmethylated), red (bisulfite converted, i.e. GpU) or blue (unconverted). Colors of the heatmap depict protection (red) or accessibility (blue) at GpC positions of an amplicon.

**f.** Purity of FACS-sorted populations. Immunofluorescence staining for TNP2 confirmed a higher abundance of cells at earlier stages of differentiation in EESts compared to LESts populations[8].

**g.** and **h.** Pearson correlation coefficients of amplicon GCH methylation (**g**) and endogenous CpG (WCG) methylation (**h**). The percentage of GCH and WCG methylation was calculated for each amplicon and sample replicate.

**i.** and **j.** ATAC-seq shows variability in chromatin accessibility in RSts, particularly at hypomethylated regions, while it was uniform in ESts[9]. The data are shown for individual amplicons (**i**) or for 1 kb genomic tiles (**j**). Normalized RPKM counts were calculated using all fragment sizes.

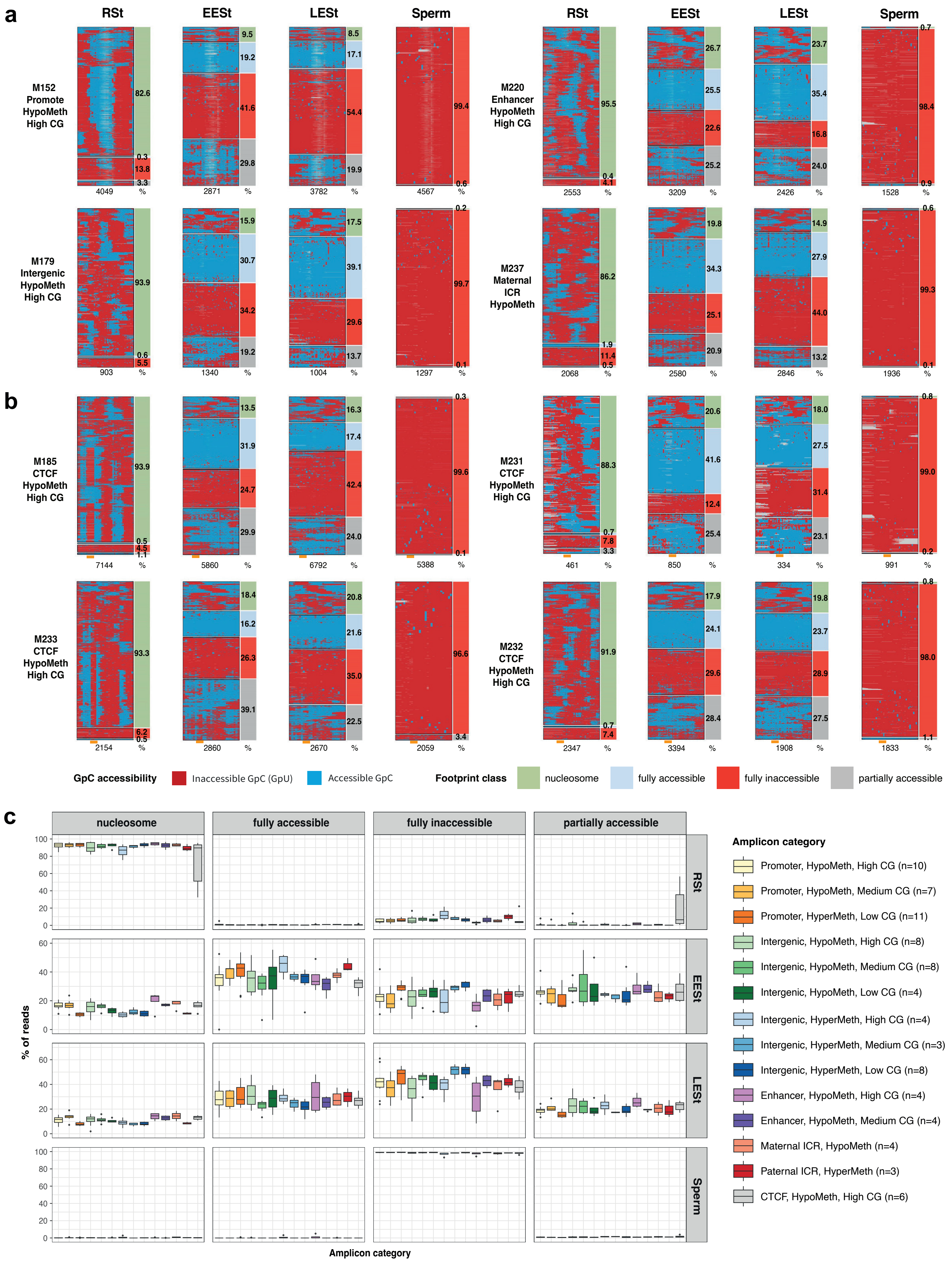


**Extended Data Figure 2**

**d**

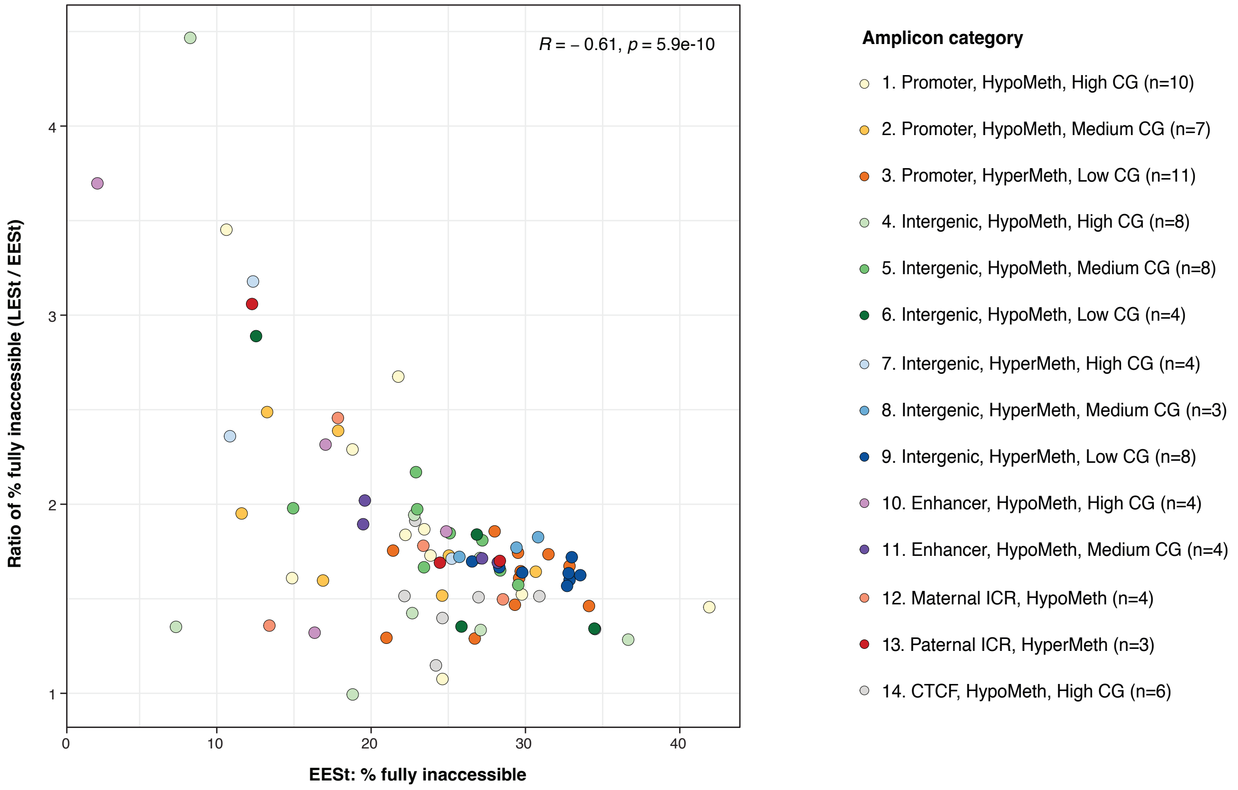


**Extended Data Figure 2 (cont.)**

**Extended Data Fig. 2: Extensive chromatin remodeling during spermatid development in mice.**

**a.** and **b.** NOMe-seq. data at representative amplicons for promoter, intergenic, enhancer and ICR regions (**a**) and four CTCF-motif containing amplicons (**b**) in haploid male germ cells. SMF-maps visualize chromatin footprints at DNA molecules of individual spermatozoa that had been classified according to the NOMeR model. Frequencies of predicted footprint classes are on the left of each SMF-map. Each row of a SMF-map corresponds to an individual fragment and each column to a cytosine within GCH context. Nucleotides between cytosines are not shown. Accessible GCHs (methylated) are shown in blue and protected GCHs (unmethylated) in red. The number of analyzed reads is shown at the bottom of each SMF-map. For CTCF amplicons (**b**), the position of the CTCF motif is annotated as an orange box below each SMF-map. Amplicons M185 and M233 display prominent CTCF binding with nearby phased nucleosome in RSts while amplicons M231 and M232 do not.

**c.** Box plots presenting frequency distributions of four footprint types at amplicons belonging to the 14 amplicon categories in four germ cell types (extending Fig. 2c). Boxes display, 25$^{th}$, 50$^{th}$ and 75$^{th}$ percentiles and whiskers extend to 1.5 times the inter-quantile range.

**d.** Scatterplot showing the ratio of fully inaccessible footprint frequencies in LESts relative to EESts (LESt/EESt; y axis) as a function of the frequency of fully inaccessible footprints in EESt (x axis) across all amplicons. This representation, analogous to an MA plot, illustrates changes in inaccessibility from EESts to LESts in relation to the baseline frequency in EESts. The Pearson correlation coefficient (*R*) and associated p-value are indicated in the upper right corner.

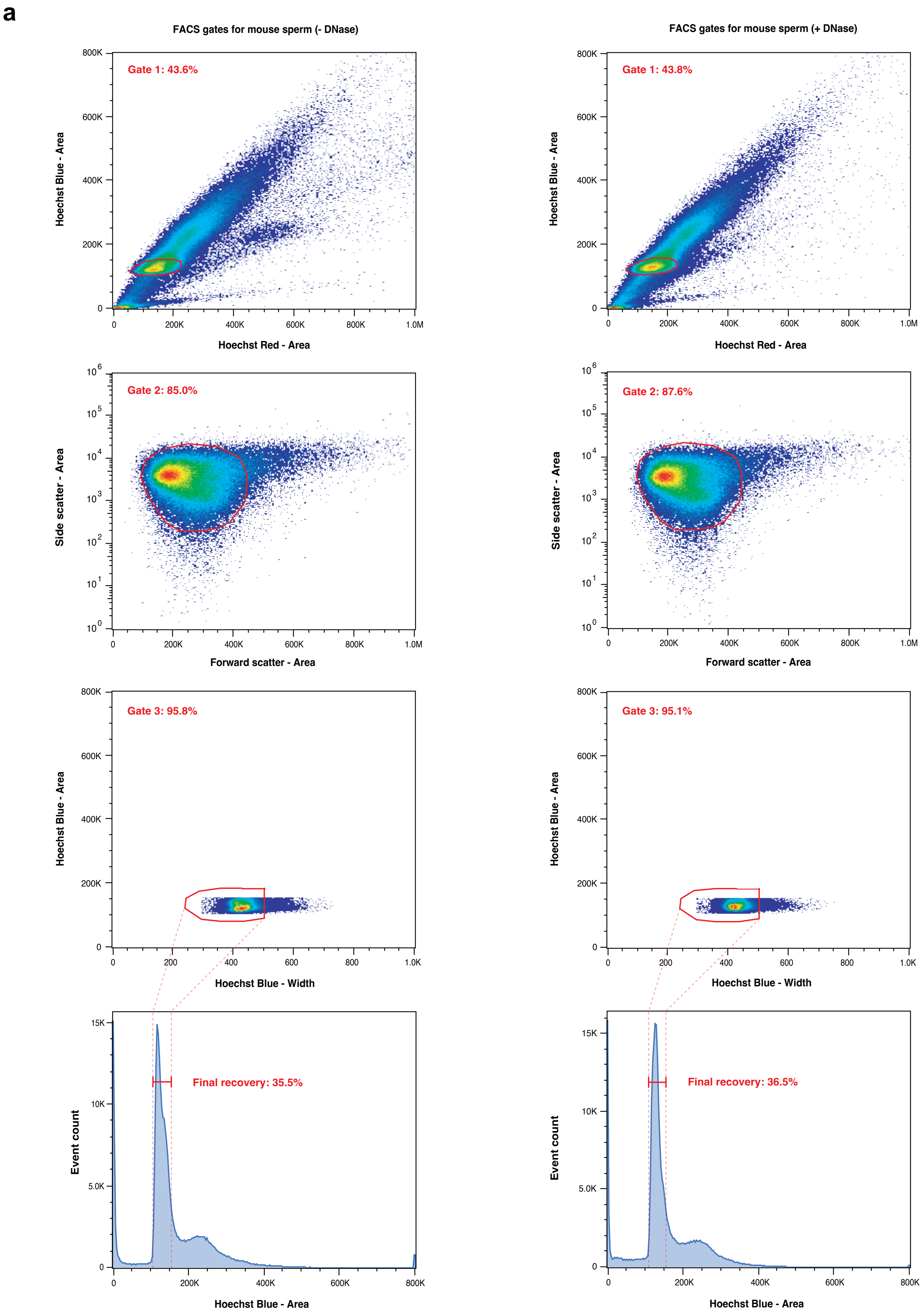


### Extended Data Figure 3

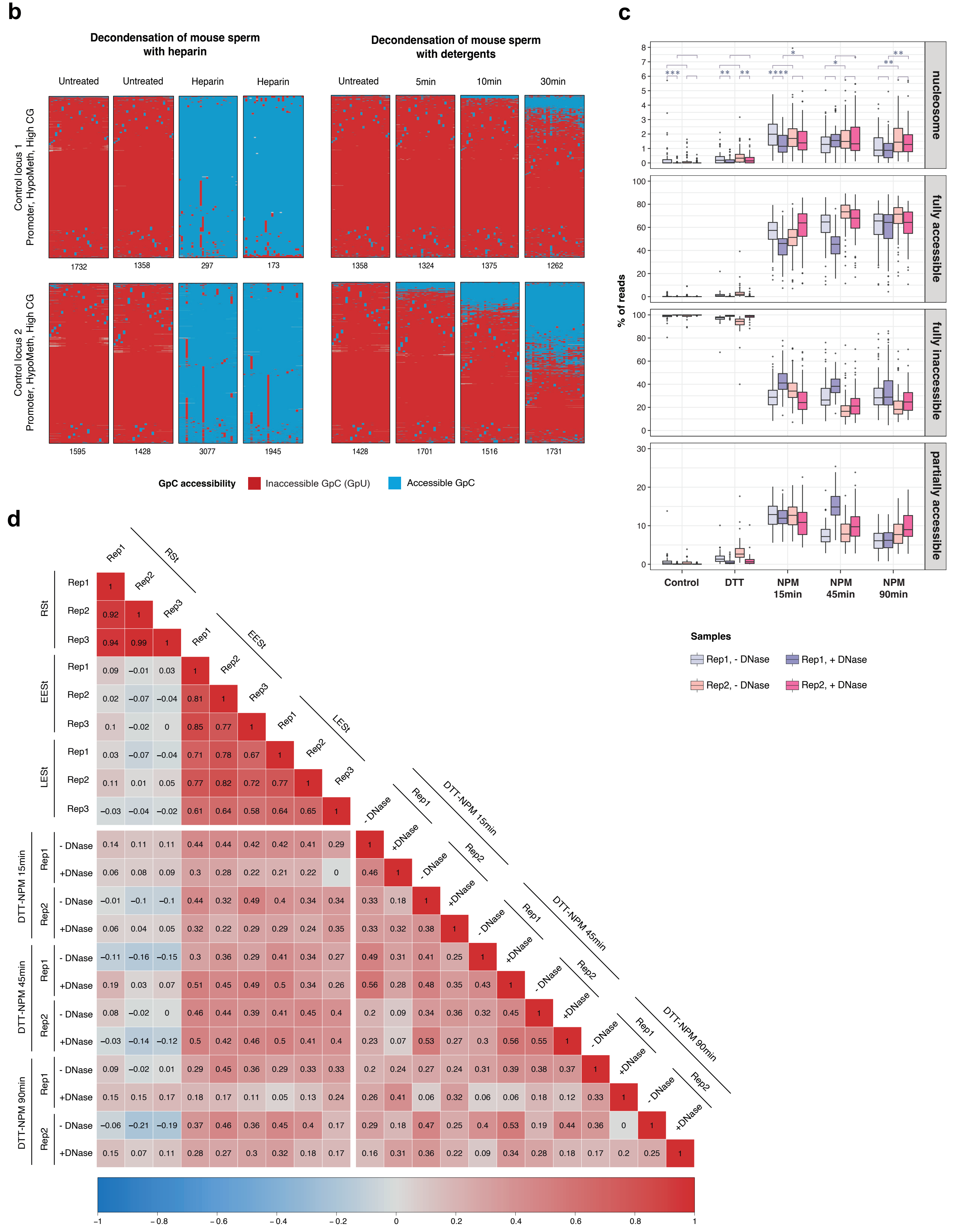


**Extended Data Figure 3 (cont.)**

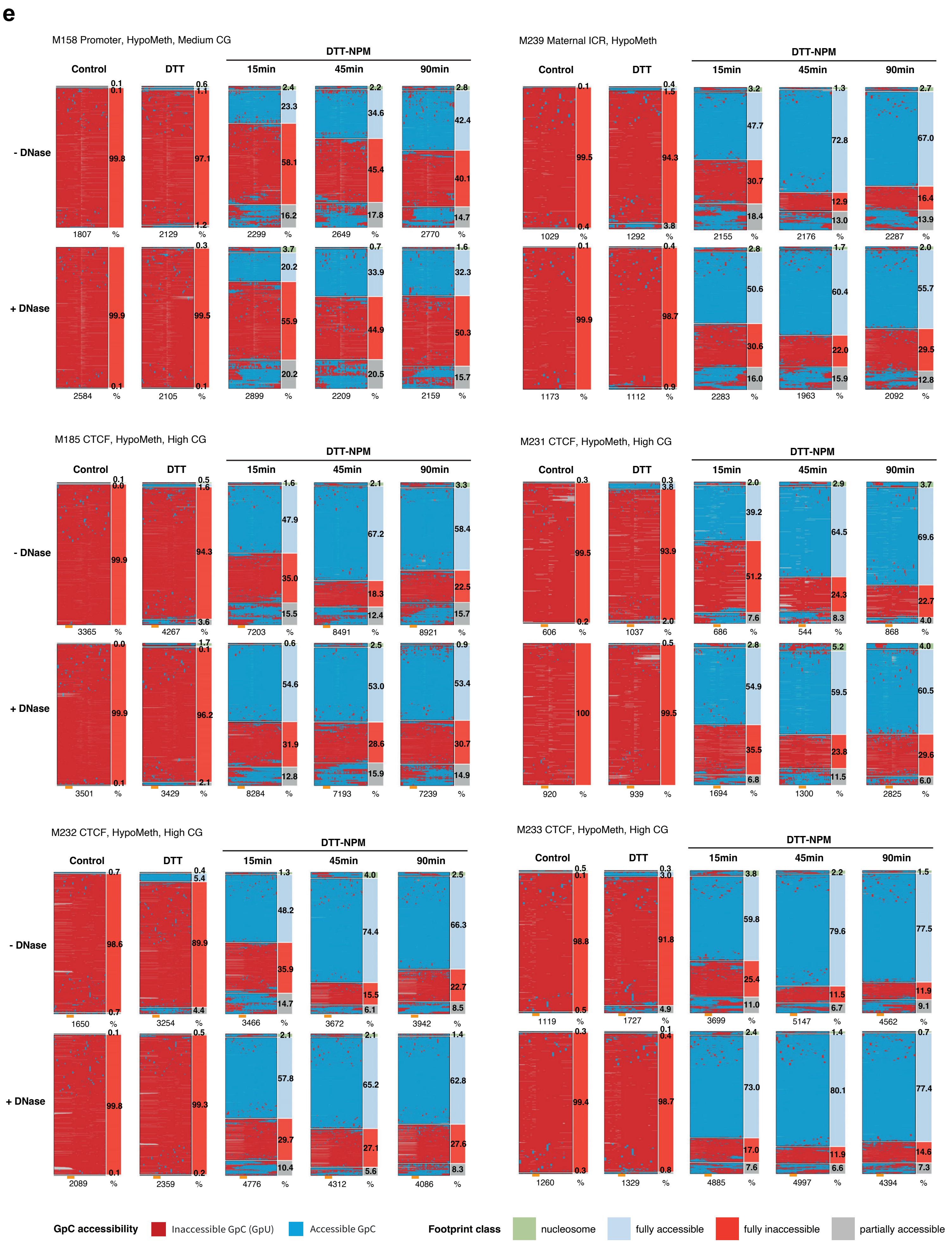


**Extended Data Figure 3 (cont.)**

**f**

**Frequency of footprints per amplicon category (- DNase)**

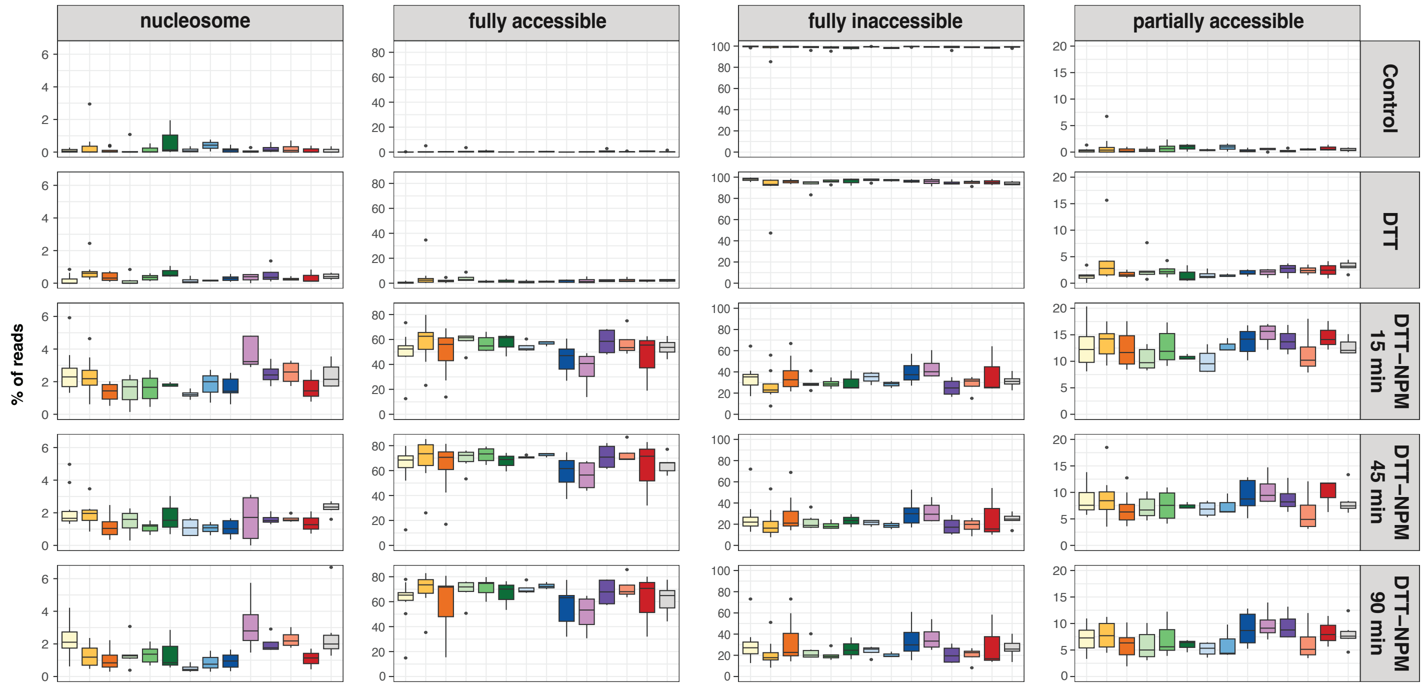


**g**

**Frequency of footprints per amplicon category (+ DNase)**

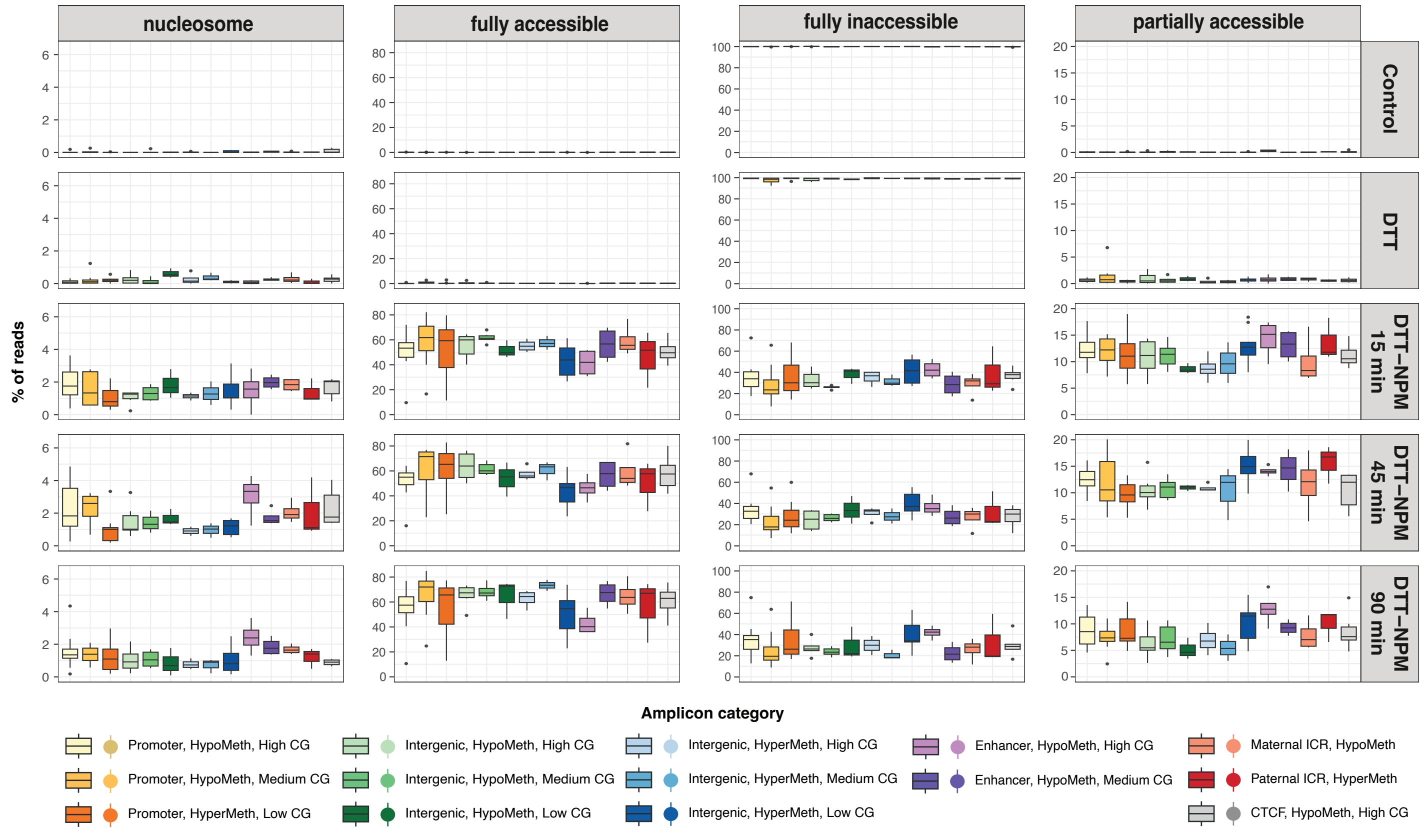


**h**

**Statistics for nucleosome occupancy across replicas and NPM incubations**

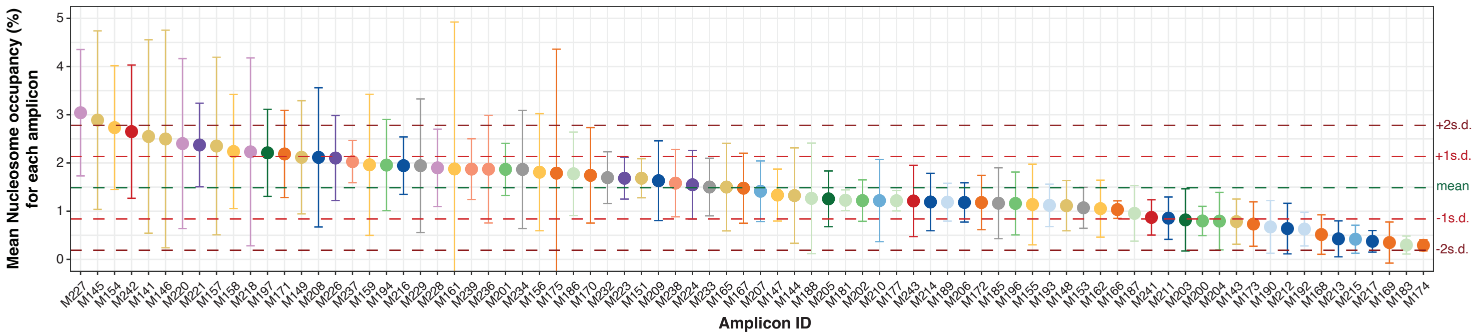


### Extended Data Figure 3 (cont.)

**Extended Data Fig. 3: NPM-mediated decondensation enables SMF of mouse spermatozoa.**

**a.** FlowJo-generated FACS sorting gates for mouse sperm without (left column) or with (right column) prior DNase I treatment. Spermatozoa were first selected for staining with the nuclear dye Hoechst according to its red and blue spectra (Gate 1) and further selected by forward and side scatter signals (Gate 2). A final sorting gate refined the selection by Hoechst blue width (Gate 3), visualized as event counts on the pre-selection axis of Hoechst blue by area and showing the percentage of final recovery (4th plot). Polygons and bars in red represent the selected sorting gates. Dashed red lines represent the corresponding boundaries of gates between plots.

**b.** Chromatin footprinting of mouse sperm decondensed with heparin (left) and detergents over several timepoints (5min, 10min and 30min) (right). SMF-maps displaying protected (red, unmethylated) or accessible (blue, methylated) GCHs at 2 control regions.

**c.** Box plots comparing the frequency distributions of four footprint types across all amplicons between the two replicates and conditions without or with DNase I treatment in FACS-sorted mouse spermatozoa (extending Fig. 3e). Data of all NPM duration conditions are shown. Boxes display 25$^{th}$, 50$^{th}$ and 75$^{th}$ percentiles and whiskers extend to 1.5 times the inter-quantile range. Differences in nucleosome occupancy between samples were assessed using pairwise two-sided Wilcoxon rank-sum tests. ns: $p>0.05$, *: $p\leqslant0.05$, **: $p\leqslant0.01$, ***: $p\leqslant0.001$, ****: $p\leqslant 0.0001$. Nucleosome footprints showed overlapping distributions with similar, statistically not significant, variabilities between DNase I conditions at the prolonged timepoints.

**d.** Pearson correlation coefficients of nucleosome footprint prediction for RSts, EESts, LESts and FACS-sorted, DTT-NPM-exposed spermatozoa treated with or without DNAse I. Nucleosome occupancy frequencies were calculated for all amplicons and sample replicates. Unlike high correlations of nucleosome occupancies at amplicons among RSts replicates and all ESt replicate samples, respectively, nucleosome frequencies at amplicons showed only moderate positive correlations among NPM-treated sperm samples across NPM timepoints and replicates (Pearson correlation coefficients <0.6), being indifferent between DNase I treatments.

**e.** NOMe-seq. data at representative promoter and maternal ICR regions and four CTCF-motif containing amplicons in mouse spermatozoa (comparable to Fig. 3c). SMF-maps visualize chromatin footprints at DNA molecules of individual spermatozoa, that had been classified according to the NOMeR model. Frequencies of predicted footprint classes are shown on the left of each SMF-map. Sperm was pretreated without or with DNAse I and further as indicated. Each row of a SMF-map corresponds to an individual fragment and each column to a cytosine within GCH context. Accessible GCHs (methylated) are shown in blue and protected GCHs (unmethylated) in red. Nucleotides between cytosines are not shown. The number of analyzed reads is shown at the bottom of each SMF-map. For CTCF amplicons, the position of CTCF motif is annotated as an orange box below each SMF-map.

**f.** and **g.** Box plots presenting frequency distributions of four footprint types at amplicons belonging to the 14 amplicon categories in FACS-sorted mouse spermatozoa, pretreated without (**f**) or with DNase I (**g**), followed by further DTT-NPM incubation. Data of all NPM duration conditions are shown. Boxes display, $25^{th}$, $50^{th}$ and $75^{th}$ percentiles and whiskers extend to 1.5 times the inter-quantile range (extending Fig. 3e).

**h.** Plot showing the mean value of nucleosome occupancies of each amplicon across 3 NPM duration conditions for both replicates in DNase I-treated, FACS-sorted spermatozoa. The error bars represent the standard deviation of nucleosome occupancy values across all aforementioned samples for each amplicon to reflect its variability across samples. The mean value across all amplicons is indicated by the green dashed line. The values of one and two standard deviation(s) above and below the mean are indicated by red and brown dashed lines, respectively.

**a**

| Human donor | Volume (ml) | Concentration (million/ml) | Concentration (million/ejaculate) | Normal morphology (%) | Round cell (million/ml) | Progressive motility (%) | DNA fragmentation TUNEL (ejaculate %) |
|---|---|---|---|---|---|---|---|
| **Donor A** | 2.3 | 35.43 | 81.49 | 7.7 | 0.2 | 59 | / |
| **Donor B** | 2.7 | 99.95 | 269.87 | 10.0 | 0.0 | 61 | 2.8 |
| **Donor C** | 3.2 | 50.08 | 160.26 | 19.9 | 0.6 | 42 | / |
| **Reference value** | ≥ 1.5 | ≥ 15 | ≥ 39 | ≥ 4.0 | < 5 | ≥ 32 | / |

**b**

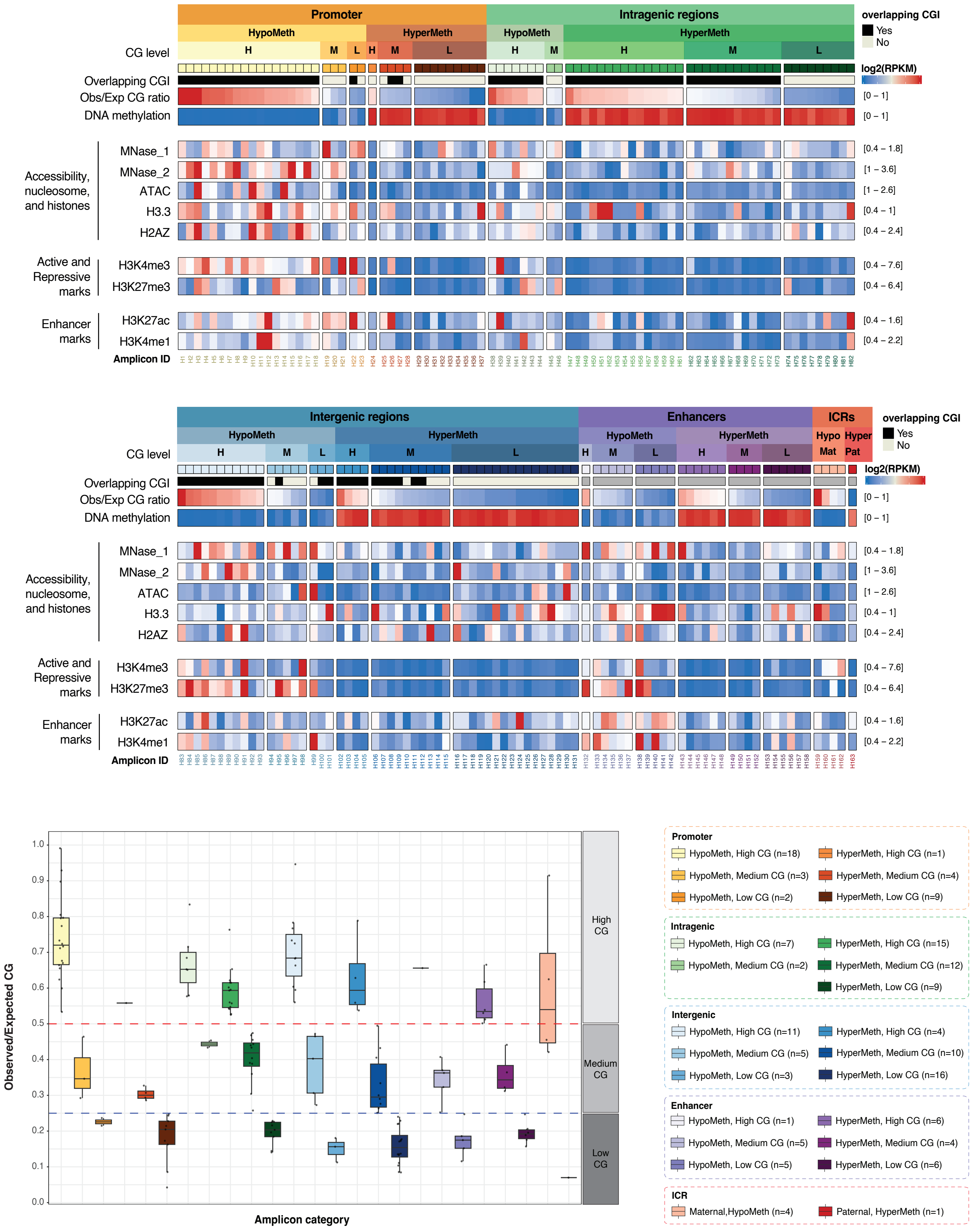


**Extended Data Figure 4**

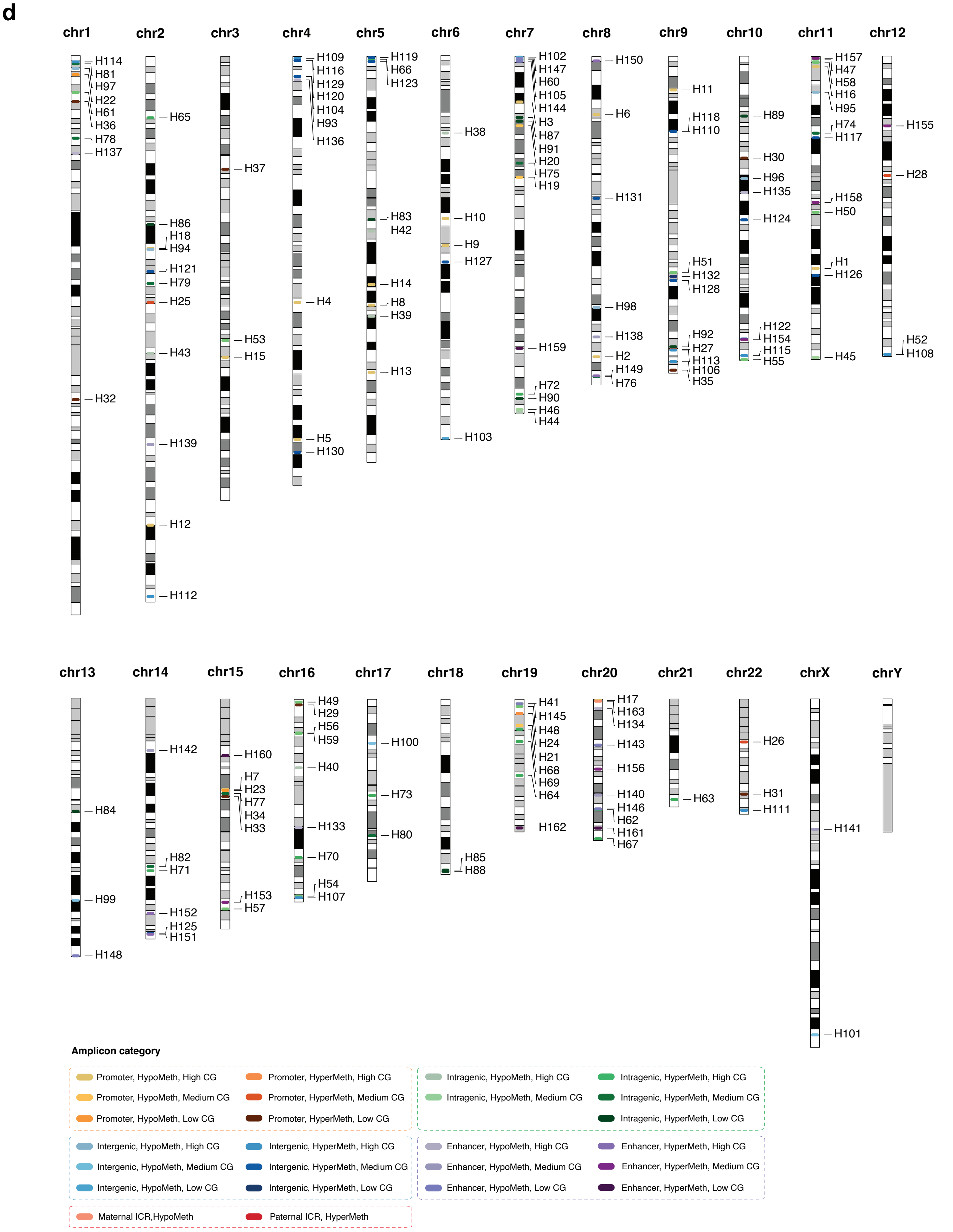


**Extended Data Figure 4 (cont.)**

e

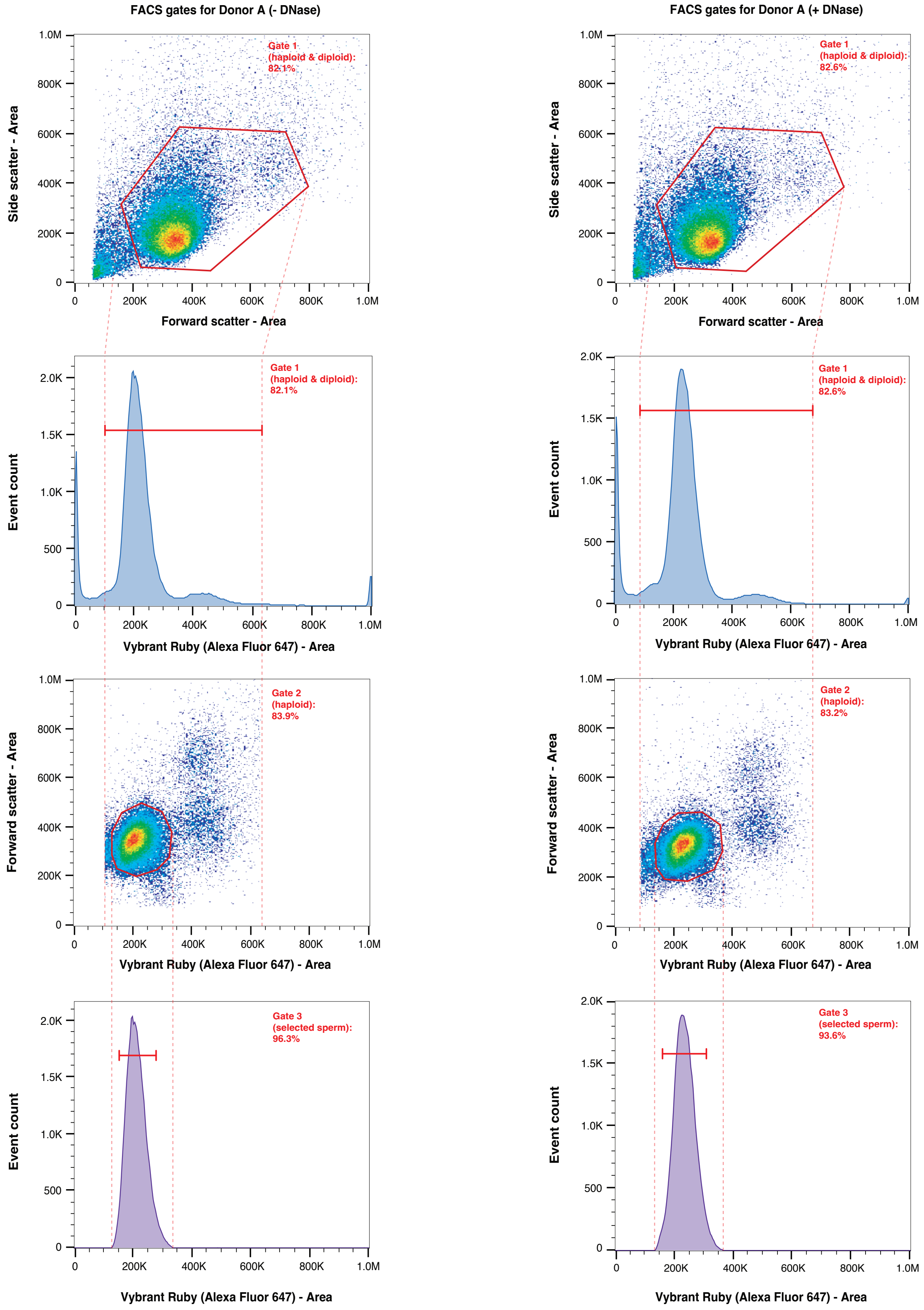


f

| Human donor | - DNase | | | | + DNase | | | |
|---|---|---|---|---|---|---|---|---|
| | Gate 1 | Gate 2 | Gate 3 | Final recovery | Gate 1 | Gate 2 | Gate 3 | Final recovery |
| Donor A | 82.1% | 83.9% | 96.3% | 66.3% | 82.6% | 83.2% | 93.6% | 64.3% |
| Donor B | 81.1% | 84.9% | 90.9% | 62.6% | 84.1% | 89.8% | 91.1% | 68.8% |
| Donor C | 66.0% | 88.0% | 96.3% | 55.9% | 66.8% | 86.9% | 92.8% | 53.8% |

**Extended Data Figure 4 (cont.)**

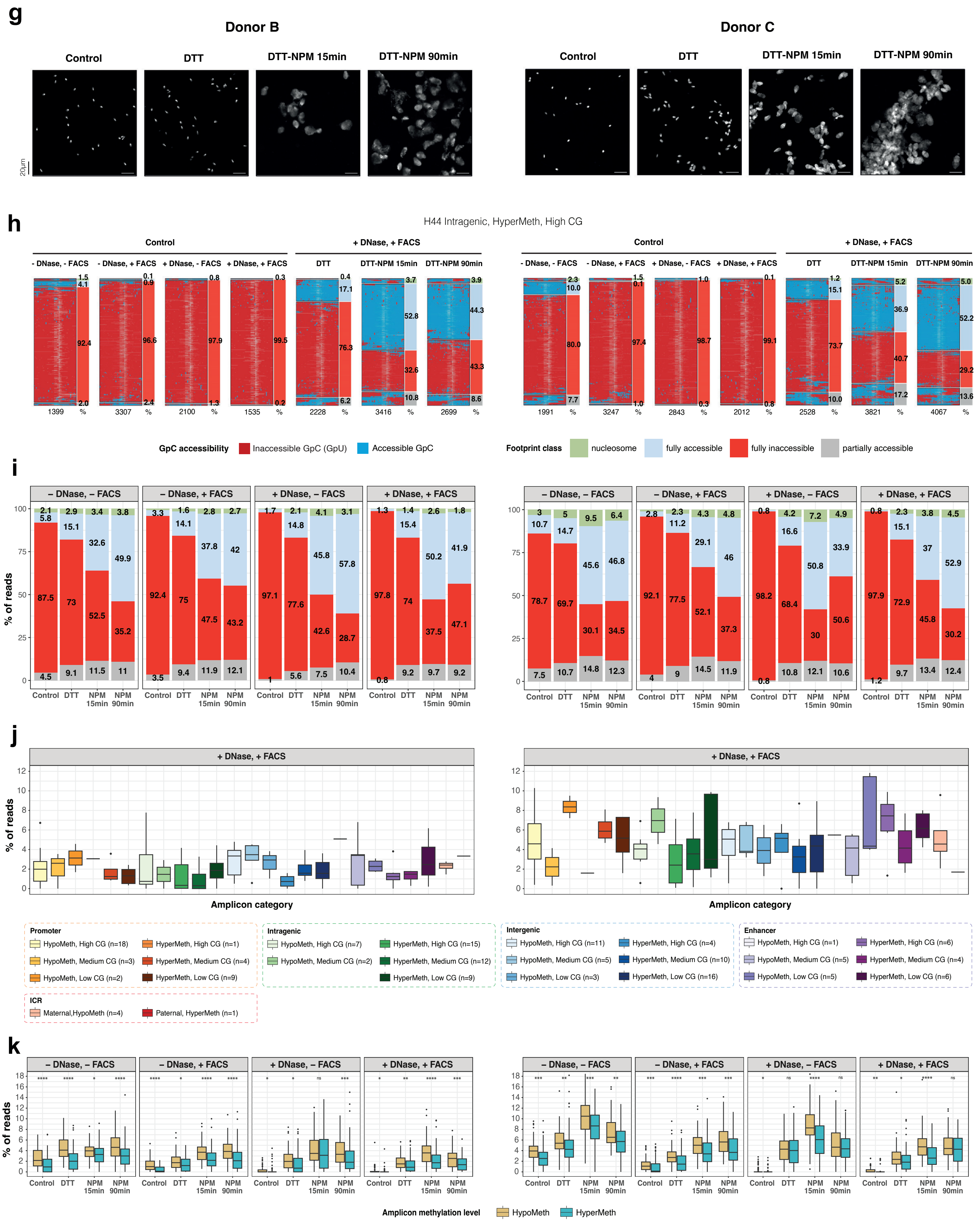


### Extended Data Figure 4 (cont.)

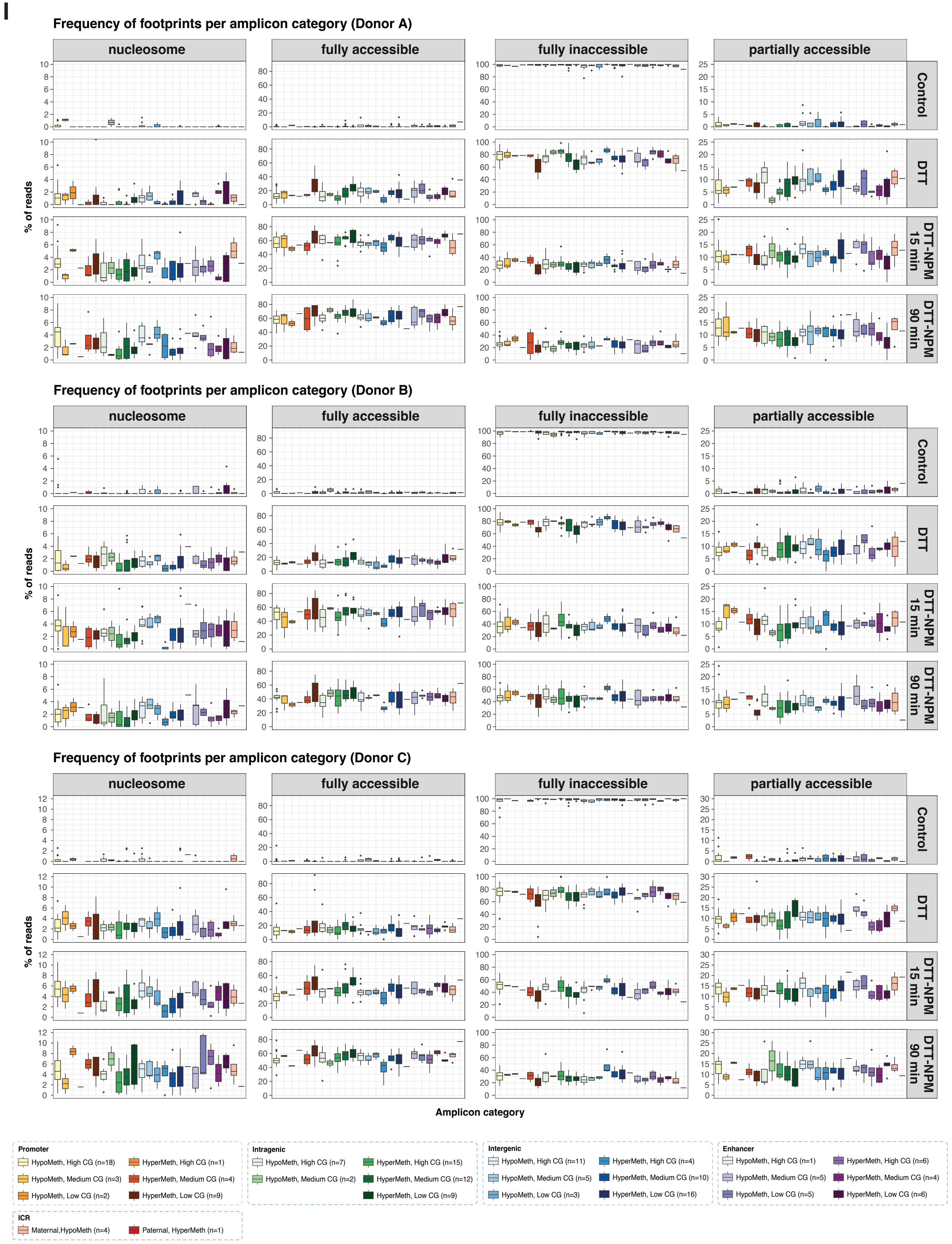


**Extended Data Figure 4 (cont.)**

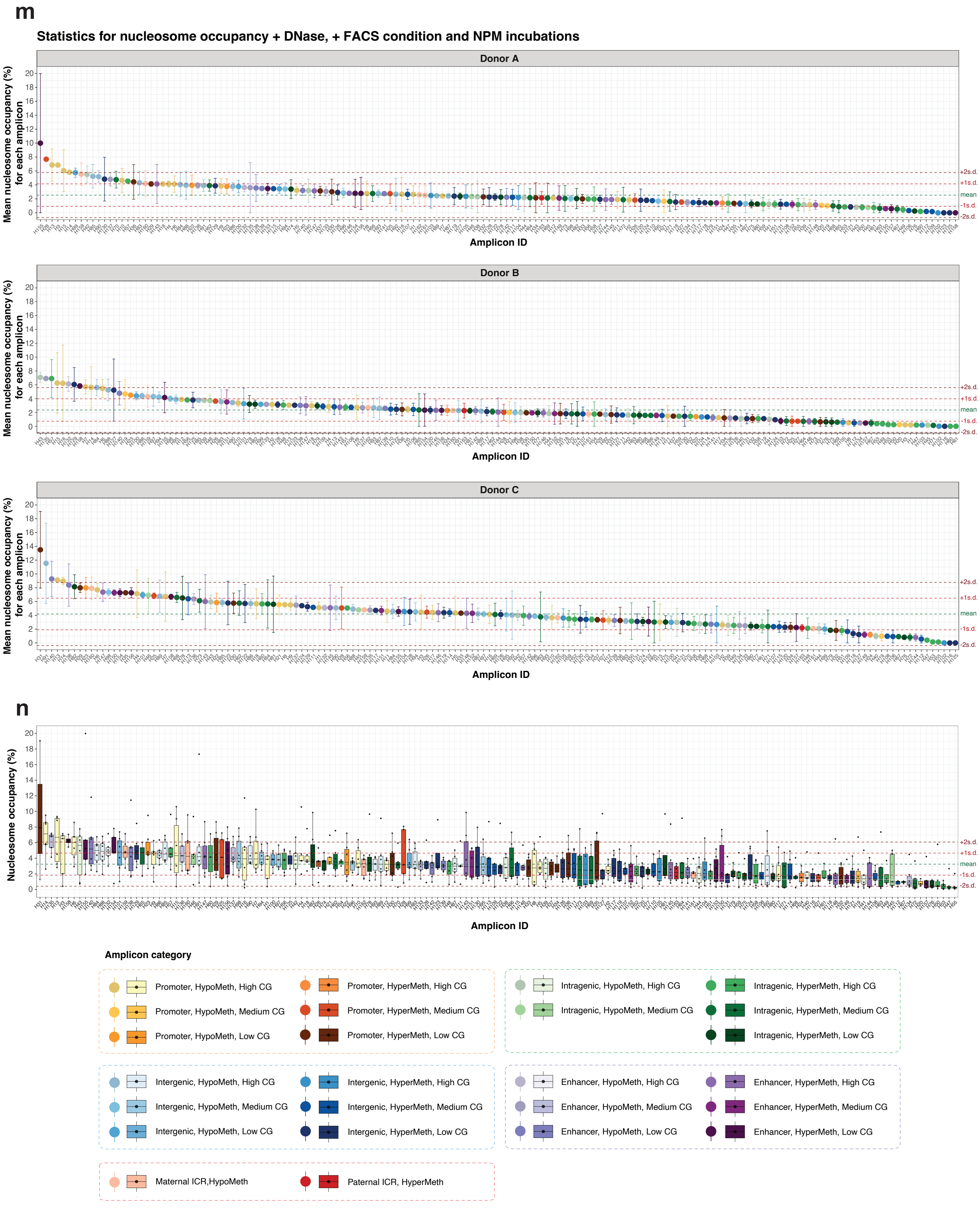


**Extended Data Figure 4 (cont.)**

**Extended Data Fig. 4: SMF profiling of human spermatozoa: experimental conditions and quantifying nucleosome occupancies and levels of local accessibilities.**

**a.** Spermiogram parameters of human donors A, B and C, and the reference value for fertile men according to WHO guideline[10].

**b.** Scheme showing chromatin characteristics at amplicons that had previously been measured by MNase-/ChIP-seq in human spermatozoa as follows: DNA methylation, H3.3, H2AZ, H3K4me3, H3K27me3, H3Kme1, H3K27ac , MNase_2[1]; ATAC[7]; MNase_1[11]. Amplicon IDs are shown at the bottom of the plot.

**c.** Box plot showing Observed/Expected CG ratios of human amplicons per amplicon category defined in panels b. Horizontal dashed lines indicate thresholds used to categorize amplicons other than ICRs into High, Medium and Low CG classes.

**d.** Karyogram showing the positions of selected amplicons on human chromosomes, colored according to their respective amplicon categories.

**e.** FlowJo-generated FACS sorting gates for Donor A sperm without (left column) or with (right column) prior DNase I treatment. Spermatozoa were selected based on forward and side scatters signals (Gate 1), then visualized as event counts on the Y-axis based on staining with the nuclear dye Vybrant Ruby (2nd plot). Gate 1 included cells with large forward and side scatter areas since the orientation of sperm with retained tails may be detected as larger in scatter areas. Gate 1 retained, however, a population of diploid cells (2nd plot). Haploid cells were selected based on Vybrant Ruby staining (Gate 2 in the 3rd plot), which are visualized as event counts (4th plot). A final sorting Gate 3 refined the selection by Vybrant Ruby. Polygons and bars in red represent the selected sorting gates. Dashed red lines represent the corresponding boundaries of gates between plots.

**f.** Summary table for sorting percentages after each sorting gate and final recovery rates for sperm from donors A, B, and C, both with and without prior DNase I treatment.

**g.** Microscopy images of DAPI-stained sperm nuclei from human donors B and C. Sperm were pretreated with DNAse I, FACS-sorted and subjected to DTT and NPM treatment conditions as described in Fig. 3a.

**h.** NOMe-seq. SMF-maps visualizing chromatin footprints at one representative amplicon for individual sperm molecules of human donors B and C, that had been classified according to the NOMeR model and annotated with frequencies of predicted footprint classes on the left of each SMF-map. SMF-maps corresponding to sperm pretreated without/with DNAse I, and without/with FACS sorting are illustrated for the control condition; SMF-maps corresponding to DNase I-treated, FACS-sorted sperm are further illustrated for DTT and NPM incubations as indicated. Every row corresponds to a unique sequence read and each column to a cytosine in a GCH context. Nucleotides between cytosines are not shown. Each row of a SMF-map corresponds to an individual fragment and each column to a cytosine within GCH context.

Accessible GCHs (methylated) are shown in blue and protected GCHs (unmethylated) in red. The number of analyzed reads is shown at the bottom of each SMF-map.

**i.** Bar plots displaying the average percentage of fragments containing nucleosome, fully accessible, fully inaccessible or partially accessible footprints for all amplicons in control, DTT or DTT-NPM-exposed spermatozoa from human donors B and C, priorly treated without/with DNase I or without/with FACS sorting.

**j.** Box plots representing the distribution of nucleosome occupancy frequencies at 163 amplicons belonging to 25 amplicon categories in DNase I-pretreated, FACS-sorted, DTT pretreated and 90min NPM-exposed human spermatozoa of donors B and C. Boxes display, $25^{th}$, $50^{th}$ and $75^{th}$ percentiles and whiskers extend to 1.5 times the inter-quantile range.

**k.** Box plots comparing the distribution of nucleosome occupancy frequencies across all amplicons, grouped into hypomethylated and hypermethylated categories, in control, DTT or DTT-NPM-exposed spermatozoa from human donors B and C, priorly treated without/with DNase I or without/with FACS sorting. Boxes display, $25^{th}$, $50^{th}$ and $75^{th}$ percentiles and whiskers extend to 1.5 times the inter-quantile range. Statistical significance was evaluated using two-sided Wilcoxon rank-sum tests. ns: $p>0.05$, *: $p\leqslant0.05$, **: $p\leqslant0.01$, ***: $p\leqslant0.001$, ****: $p\leqslant0.0001$.

**l.** Box plots representing the frequency distributions of four footprint types at 163 amplicons belonging to 25 amplicon categories in DNase I-pretreated, FACS-sorted, DTT pretreated and NPM-exposed human spermatozoa of donors A, B and C. Data of both NPM duration conditions are shown. Boxes display, $25^{th}$, $50^{th}$ and $75^{th}$ percentiles and whiskers extend to 1.5 times the inter-quantile range.

**m.** Plots showing the mean value of nucleosome occupancies of each amplicon across both NPM duration conditions in DNase I-treated, FACS-sorted human spermatozoa of donors A, B and C. The error bars represent the upper and lower values of nucleosome occupancies at the two NPM duration conditions. The mean value across all amplicons is indicated by the green dashed line. The values of one and two standard deviation(s) above and below the mean are indicated by red and brown dashed lines, respectively.

**n.** Box plot representing the distribution of nucleosome occupancies of each amplicon across both NPM duration conditions in DNase I-treated, FACS-sorted human spermatozoa of all three donors A, B and C. The mean value across all amplicons is indicated by the green dashed line. The values of one and two standard deviation(s) above and below the mean are indicated by red and brown dashed lines, respectively.

### Supplementary Table 1

| Organism | ID | Class | Chromosome | Start | End | Strand | Fwseq | Rvseq | Full frag. length | GCH inform. length | GCH number | GCH MaxGap | WCG number | obs/exp CG | o/e CG level | CGI class | Methylation class | Promoter SYMBOL | Edges |
|---|---|---|---|---|---|---|---|---|---|---|---|---|---|---|---|---|---|---|---|
| Mouse | M141 | Promoter_HypoMeth_highCG | chr6 | 89595208 | 89595773 | + | TGAATAAGATTGGAAATGGAAGAATA | CACAAATACCCCTACCTCAAAAA | 565 | 504 | 49 | 35 | 16 | 0.85385 | highCG | CGI | HypoMeth | Chchd6 | FALSE |
| Mouse | M142 | Promoter_HypoMeth_highCG | chr17 | 65772234 | 65772811 | - | TATAAGTTAGATGTTGGGTGGTTATT | TATCCTCTAATCCTCAACAATCCC | 577 | 494 | 50 | 34 | 11 | 0.83094 | highCG | CGI | HypoMeth | Rab31 | FALSE |
| Mouse | M143 | Promoter_HypoMeth_highCG | chr2 | 122298637 | 122299216 | - | TTTTAGGGAGAATTTTGAATATTTTT | AAAATCTTCACCCTTCCTAAAT | 579 | 522 | 34 | 33 | 11 | 0.70436 | highCG | CGI | HypoMeth | Duox2 - Duoxa2 | FALSE |
| Mouse | M144 | Promoter_HypoMeth_highCG | chr6 | 52158289 | 52158819 | + | AGGATTAGAGAAGAAAAATGATATGAA | AATATCTATTAAATTCCCACCCTTCC | 530 | 457 | 38 | 29 | 7 | 0.73985 | highCG | CGI | HypoMeth | Hotairm1 - Hoxa1 | FALSE |
| Mouse | M145 | Promoter_HypoMeth_highCG | chr2 | 57237531 | 57238090 | + | TTTAGGATTTAAAGTGGGGTTTTT | TATAATATTAAATTTCTATCCCTCCCCA | 559 | 487 | 41 | 37 | 8 | 0.55028 | highCG | CGI | HypoMeth | Gpd2 | FALSE |
| Mouse | M146 | Promoter_HypoMeth_highCG | chr14 | 59625185 | 59625735 | + | ATATTTTTGGTGAAGGAAAGTGAAGG | TTTCCCCCTCTATTCACTTCCC | 550 | 500 | 59 | 35 | 9 | 0.81687 | highCG | CGI | HypoMeth | Shisa2 | FALSE |
| Mouse | M148 | Promoter_HypoMeth_highCG | chr13 | 98353884 | 98354390 | + | AGTTGTTTTTAGTGGGATGATTTTTA | TAACCTTTTCCACTATAATCCCTTTA | 506 | 451 | 33 | 35 | 7 | 0.58315 | highCG | CGI | HypoMeth | Foxd1 | FALSE |
| Mouse | M149 | Promoter_HypoMeth_highCG | chr13 | 64370044 | 64370614 | + | GAGTTATATGATTTGATGAAGATTT | ACATTTTAATCTACAACTTTCATACA | 570 | 504 | 37 | 36 | 10 | 0.72254 | highCG | CGI | HypoMeth | Ctsl | FALSE |
| Mouse | M150 | Promoter_HypoMeth_highCG | chr2 | 134643821 | 134644400 | - | TTTGTTAATATGAAATGGGGGATG | AAATAAATAAAAACACTCACAATTTCA | 579 | 518 | 50 | 36 | 8 | 0.65579 | highCG | CGI | HypoMeth | Tmx4 | TRUE |
| Mouse | M151 | Promoter_HypoMeth_highCG | chr16 | 18836403 | 18836946 | - | AGTTTGGTATTTAGGAGAGAAATA | AAAAACTAACCAAAACTCCTATTTTT | 543 | 476 | 42 | 34 | 13 | 0.69050 | highCG | CGI | HypoMeth | Hira - 2510002D24Rik | FALSE |
| Mouse | M152 | Promoter_HypoMeth_highCG | chr5 | 144357919 | 144358448 | + | GATTTGAGATTGAGTGGTTTTTG | CCATCAATCCATTTCAAATAAAAA | 529 | 467 | 36 | 33 | 10 | 0.68749 | highCG | CGI | HypoMeth | Dmrt1i - Baiap2l1 | TRUE |
| Mouse | M157 | Promoter_HypoMeth_highCG | chr2 | 38931772 | 38932331 | - | TTTGATTTTAGGAGAGTTTAGATTAAGT | AATCCCTCATTTCACAAAACCTAA | 559 | 476 | 51 | 27 | 6 | 0.53319 | highCG | nonCGI | HypoMeth | Olfml2a | FALSE |
| Mouse | M160 | Promoter_HypoMeth_highCG | chr9 | 13246785 | 13247349 | - | TAGAAAGATAAGGGTTTTAGGAAAGA | ACTTAACTCTATCTTCTCTAACCCTA | 564 | 505 | 45 | 29 | 8 | 0.67459 | highCG | nonCGI | HypoMeth | Ccdc82 | FALSE |
| Mouse | M163 | Promoter_HypoMeth_highCG | chr8 | 122460534 | 122461113 | - | TTGTTTATTTATTTTAGAGATTGTTTTATT | CTACCTCTCAATATCTCCAATTTC | 579 | 465 | 43 | 34 | 14 | 0.77765 | highCG | nonCGI | HypoMeth | Snai3 | TRUE |
| Mouse | M164 | Promoter_HypoMeth_highCG | chr5 | 136907867 | 136908438 | - | TTTTGAGATTTGGGTATTTTAATATTTTT | ACCAAAATCACAATACTATTTTCCAA | 571 | 483 | 40 | 32 | 8 | 0.57440 | highCG | nonCGI | HypoMeth | Ift22 | TRUE |
| Mouse | M165 | Promoter_HypoMeth_highCG | chr19 | 57452699 | 57453267 | + | AGAAAGGAAGTGATTTTATTTTAAAT | AAAAACTAAAAATAAAAATCCCAATA | 568 | 462 | 48 | 33 | 7 | 0.54712 | highCG | nonCGI | HypoMeth | Trub1 | FALSE |
| Mouse | M147 | Promoter_HypoMeth_mediumCG | chr13 | 55209454 | 55210006 | + | TTTTTAATGGATTGTGGGTTTTG | TCCATTCCTCAAAAACAATAAAAA | 552 | 484 | 34 | 36 | 8 | 0.42182 | mediumCG | CGI | HypoMeth | Nsd1 | FALSE |
| Mouse | M154 | Promoter_HypoMeth_mediumCG | chr5 | 90903672 | 90904251 | + | GGATATTTTAGGGTTTTATAGTGGAA | AAAACACTCACATAACTTCTATCTAA | 579 | 525 | 44 | 32 | 7 | 0.32984 | mediumCG | nonCGI | HypoMeth | Cxcl2 | FALSE |
| Mouse | M155 | Promoter_HypoMeth_mediumCG | chr15 | 85016955 | 85017531 | - | GGTTATTATAGGATTGGGTGAGTTT | TCTCTAACAACACTATAAAAACCTCA | 576 | 525 | 46 | 35 | 5 | 0.38855 | mediumCG | nonCGI | HypoMeth | Upk3a | FALSE |
| Mouse | M156 | Promoter_HypoMeth_mediumCG | chr8 | 45507726 | 45508288 | + | ATTAGATGAAGAGGGTTTAAAGATTT | TCATCCTTCCTATCCTATATTCTTAA | 562 | 496 | 44 | 33 | 7 | 0.37701 | mediumCG | nonCGI | HypoMeth | Sorbs2 | FALSE |
| Mouse | M158 | Promoter_HypoMeth_mediumCG | chr2 | 65363784 | 65364361 | - | GTAGGGGGAGGAATATTTTGTAAA | TCAAATTAAATCAAATCTATCCCTTTT | 577 | 501 | 46 | 31 | 3 | 0.39228 | mediumCG | nonCGI | HypoMeth | Mir6337 - Slc38a11 | FALSE |
| Mouse | M159 | Promoter_HypoMeth_mediumCG | chr6 | 59024136 | 59024715 | + | TATAAAAGTGTTAGTGAAAGAGAAGT | CAAACTTATCTAAAACCTCTCAAAAA | 579 | 512 | 46 | 35 | 6 | 0.41572 | mediumCG | nonCGI | HypoMeth | Fam13a | FALSE |
| Mouse | M161 | Promoter_HypoMeth_mediumCG | chr2 | 131127159 | 131127721 | + | GGTTTTTGAGGGTTTGGAATTTTT | TTCTCCAAACTAAAACTCCAATAAAA | 562 | 504 | 53 | 26 | 6 | 0.41325 | mediumCG | nonCGI | HypoMeth | Hspa12b | FALSE |
| Mouse | M162 | Promoter_HypoMeth_mediumCG | chr10 | 85127693 | 85128270 | - | GGGATAAAGAGATTGAATTATTAGAGT | TAAAACAATCCTCTCAATAAAACCAA | 577 | 479 | 37 | 35 | 9 | 0.40637 | mediumCG | nonCGI | HypoMeth | Mterf2 | FALSE |
| Mouse | M166 | Promoter_HyperMeth_lowCG | chr5 | 30968443 | 30969018 | + | TTTTGTTAGGTGTTAGGGTTAGTTAT | CCTCCTATCAAAATCCTTACCCT | 575 | 515 | 46 | 28 | 3 | 0.20109 | lowCG | nonCGI | HyperMeth | Tcf23 | FALSE |
| Mouse | M167 | Promoter_HyperMeth_lowCG | chr10 | 127759636 | 127760195 | + | TATATTTAAGAGGGAAGAAATGAGTG | TTAATAACATCCAAAATCACTATCTCC | 559 | 471 | 37 | 33 | 4 | 0.17716 | lowCG | nonCGI | HyperMeth | Rdh9 - Rdh1 | FALSE |
| Mouse | M168 | Promoter_HyperMeth_lowCG | chr11 | 100087775 | 100088348 | + | GTAGGATTGGTAGTTAGGATATATGG | AACACTCCCCCATAAAAATACTTT | 573 | 517 | 45 | 32 | 5 | 0.24539 | lowCG | nonCGI | HyperMeth | Krt32 | FALSE |
| Mouse | M169 | Promoter_HyperMeth_lowCG | chr7 | 46240532 | 46241101 | + | TTTTATTGAATAGAGGATGGGGATTT | AATAAACTCAACCCTAAATAAACAAAA | 569 | 513 | 40 | 33 | 2 | 0.06567 | lowCG | nonCGI | HyperMeth | Otog | FALSE |
| Mouse | M170 | Promoter_HyperMeth_lowCG | chr1 | 90014281 | 90014855 | + | TAAGGGTATGTTATTTTGTTGGAATT | CTCAAATAAACAAAATCACCTCTAAAA | 574 | 517 | 44 | 30 | 5 | 0.23041 | lowCG | nonCGI | HyperMeth | Asb18 | FALSE |
| Mouse | M171 | Promoter_HyperMeth_lowCG | chr1 | 36274381 | 36274940 | - | AGGTTTTGTTTGGGGATTATATTATT | ATCCCTTCCTCTTCTTAAAATCTAAT | 559 | 482 | 41 | 29 | 1 | 0.11580 | lowCG | nonCGI | HyperMeth | Neurl3 | FALSE |
| Mouse | M172 | Promoter_HyperMeth_lowCG | chr2 | 26973086 | 26973663 | + | ATAGGAGATTTAGATGAAGTTTGGAA | TAATCCCACCTTCTAAAAATCAAAAA | 577 | 517 | 38 | 30 | 1 | 0.09441 | lowCG | nonCGI | HyperMeth | Adamts13 | FALSE |
| Mouse | M173 | Promoter_HyperMeth_lowCG | chr15 | 78468386 | 78468964 | - | TAGATTTATGGAAGAGAGTGTTAAGT | CAAAAATCCTAATTCTCTATCCCAAA | 578 | 525 | 40 | 34 | 2 | 0.13335 | lowCG | nonCGI | HyperMeth | Tmprss6 | FALSE |
| Mouse | M174 | Promoter_HyperMeth_lowCG | chr2 | 180986352 | 180986916 | - | GTGTGTAGTATATGAAATTAGATTGAAAA | CAATCTTCATCAACATCTCCATAATA | 564 | 481 | 38 | 31 | 1 | 0.06071 | lowCG | nonCGI | HyperMeth | Col20a1 | FALSE |
| Mouse | M175 | Promoter_HyperMeth_lowCG | chr14 | 20268934 | 20269513 | - | GAGGAATTTGAAGTTGTTAAGATTTT | CTTTTATACTCTCACCTATACAAACT | 579 | 517 | 52 | 31 | 1 | 0.10590 | lowCG | nonCGI | HyperMeth | Kcnk16 | FALSE |
| Mouse | M176 | Promoter_HyperMeth_lowCG | chr2 | 27079134 | 27079711 | - | TTTGTGGTTGAATATTTTTATTAGGG | CCATTAAAATCAATCACTCTAAACCT | 577 | 523 | 51 | 35 | 2 | 0.20778 | lowCG | nonCGI | HyperMeth | Adamtsl2 | FALSE |
| Mouse | M177 | Intergenic_HypoMeth_highCG | chr1 | 12996102 | 12996606 | - | GAAGGAGGATTTAGGATTATGAAT | AAAAATTACACACTAAAACTAAAATTT | 504 | 439 | 30 | 33 | 10 | 0.68439 | highCG | CGI | HypoMeth | NA | FALSE |
| Mouse | M178 | Intergenic_HypoMeth_highCG | chr1 | 86434145 | 86434640 | + | GAGGAGGGAGAAGAAAAATATTGT | CTAAAAACAAACACAAATAATTCCCA | 495 | 445 | 42 | 37 | 8 | 0.55668 | highCG | CGI | HypoMeth | NA | TRUE |
| Mouse | M179 | Intergenic_HypoMeth_highCG | chr5 | 33692372 | 33692914 | - | AATATTGAAAGATTTAAGAAGGTTTTT | TATTTTCACCCATAAAATTCCAAAAA | 542 | 470 | 48 | 34 | 8 | 0.50371 | highCG | CGI | HypoMeth | NA | FALSE |
| Mouse | M181 | Intergenic_HypoMeth_highCG | chr1 | 12995973 | 12996491 | + | TTTTTGTATGGAATTAGGTGAAAGG | AATCAATTTATACCACTCCACAATTT | 518 | 463 | 22 | 40 | 2 | 0.59721 | highCG | CGI | HypoMeth | NA | FALSE |
| Mouse | M182 | Intergenic_HypoMeth_highCG | chr11 | 107012309 | 107012868 | + | AGTATTATTTTGTATTTAGAGATTTTGTTT | TCACTACAAAATCTAAATTCTCACTC | 559 | 440 | 28 | 30 | 12 | 0.77778 | highCG | CGI | HypoMeth | NA | FALSE |
| Mouse | M183 | Intergenic_HypoMeth_highCG | chr11 | 20532213 | 20532775 | - | ATAAATAGAAGGTGATGTGTTTTAAT | TCCAAACCTTTAAATTATTATTCTTCC | 562 | 503 | 40 | 33 | 11 | 0.54291 | highCG | CGI | HypoMeth | NA | FALSE |
| Mouse | M184 | Intergenic_HypoMeth_highCG | chr3 | 34894384 | 34894893 | + | TTATTATTGAGGGAAAGGAAATTTTT | TTTCTAACTCTCAAAATCTCCAAAAA | 509 | 420 | 44 | 38 | 8 | 0.61446 | highCG | CGI | HypoMeth | NA | FALSE |
| Mouse | M186 | Intergenic_HypoMeth_highCG | chr5 | 139619055 | 139619561 | + | TTTTTAGATGGAGGTGAGTTAAA | TTATAAAAACACACATTCTTCCTTTC | 506 | 447 | 36 | 36 | 6 | 0.46179 | highCG | CGI | HypoMeth | NA | FALSE |
| Mouse | M187 | Intergenic_HypoMeth_highCG | chr10 | 126928237 | 126928750 | + | TTTTGGGAGAGAAAGAAGATATT | AAATATTTTAAAACACAAAAACCAAT | 513 | 457 | 40 | 36 | 8 | 0.67112 | highCG | CGI | HypoMeth | NA | FALSE |
| Mouse | M188 | Intergenic_HypoMeth_highCG | chr15 | 101246191 | 101246725 | - | GTGTTAGATTTTAAATTTTTGTTATT | AAAAATTAAAAACATTCACCTTTAAT | 534 | 470 | 47 | 39 | 14 | 0.87347 | highCG | CGI | HypoMeth | NA | FALSE |

Supplementary Table 1 (cont.)

| Organism | ID | Class | Chromosome | Start | End | Strand | Fwseq | Rvseq | Full frag. length | GCH inform. length | GCH number | GCH MaxGap | WCG number | obs/exp CG | o/e CG level | CGI class | Methylation class | Promoter SYMBOL | Edges |
|---|---|---|---|---|---|---|---|---|---|---|---|---|---|---|---|---|---|---|---|
| Mouse | M194 | Intergenic_HypoMeth_mediumCG | chr2 | 113893543 | 113894122 | + | AAAGAAAAAGAAAAAGAAGATAGAAA | ACTATTCTAAATACTCTATATAACCTTATA | 579 | 502 | 38 | 30 | 5 | 0.26490 | mediumCG | nonCGI | HypoMeth | NA | FALSE |
| Mouse | M196 | Intergenic_HypoMeth_mediumCG | chr15 | 92840192 | 92840751 | + | TTGAATATAGGATGTTTTGGTTAA | AAACACACTAATCACCATACAAT | 559 | 494 | 34 | 34 | 2 | 0.31189 | mediumCG | nonCGI | HypoMeth | NA | FALSE |
| Mouse | M198 | Intergenic_HypoMeth_mediumCG | chr3 | 17553670 | 17554238 | + | TTTGAAAATGTTATTTATTTGAAAAA | TATTTCTTCTCTACCAACTCTT | 568 | 504 | 34 | 37 | 6 | 0.32158 | mediumCG | nonCGI | HypoMeth | NA | FALSE |
| Mouse | M199 | Intergenic_HypoMeth_mediumCG | chr12 | 71293787 | 71294361 | + | ATTTTTAAAGATATTTATTTATGGAGTT | AAAATTAACTCAATAACCTTTCAAAA | 574 | 487 | 29 | 37 | 5 | 0.39413 | mediumCG | nonCGI | HypoMeth | NA | FALSE |
| Mouse | M200 | Intergenic_HypoMeth_mediumCG | chr15 | 3184797 | 3185375 | + | TGTATTTGGTTTTTAGTGGATTATTTT | CAATTTTCCAATCACCTATCCATTAA | 578 | 495 | 34 | 38 | 8 | 0.28229 | mediumCG | nonCGI | HypoMeth | NA | FALSE |
| Mouse | M201 | Intergenic_HypoMeth_mediumCG | chr4 | 29133373 | 29133846 | + | TGTGAGAGTGTTGAGTTTAAATT | TTACTAATAACTCCACAAACTTATTT | 473 | 407 | 32 | 37 | 5 | 0.40215 | mediumCG | nonCGI | HypoMeth | NA | FALSE |
| Mouse | M202 | Intergenic_HypoMeth_mediumCG | chr16 | 64329560 | 64330129 | + | TGATGAAGAATATAGTAGGGGAAAAA | ATATTATAAACCCAATAAAACTAAACTTTT | 569 | 488 | 32 | 34 | 3 | 0.25752 | mediumCG | nonCGI | HypoMeth | NA | FALSE |
| Mouse | M204 | Intergenic_HypoMeth_mediumCG | chr5 | 118857328 | 118857898 | - | ATTTTGAGTTAAGTTTTTGGATTGTG | TCCTAAAATCAATAATTCTCCTTTCT | 570 | 509 | 36 | 33 | 3 | 0.29307 | mediumCG | nonCGI | HypoMeth | NA | FALSE |
| Mouse | M195 | Intergenic_HypoMeth_lowCG | chr3 | 17616523 | 17617092 | + | TAATTGGTGATATAATTTTTAAGGGG | TTTACATATAAATTCAACCAAATAAAAA | 569 | 432 | 38 | 31 | 1 | 0.03575 | lowCG | nonCGI | HypoMeth | NA | FALSE |
| Mouse | M197 | Intergenic_HypoMeth_lowCG | chr2 | 63559893 | 63560461 | + | GGATAGGAGATAGGATAGTTTTT | TTCTATAATTTAAAACCTATCCCTA | 568 | 518 | 54 | 37 | 2 | 0.18948 | lowCG | nonCGI | HypoMeth | NA | FALSE |
| Mouse | M203 | Intergenic_HypoMeth_lowCG | chr4 | 31363508 | 31364068 | + | TATGTAAGGATTGGAAATGTGGG | AATTTAATCTCAAACTCTCAAAAATCT | 560 | 491 | 39 | 34 | 2 | 0.24166 | lowCG | nonCGI | HypoMeth | NA | FALSE |
| Mouse | M205 | Intergenic_HypoMeth_lowCG | chr15 | 88105418 | 88105979 | + | GTGAGTGGATGATTTTATAAATTAGGA | ATTCCCTCCCTCTTTATACCAAAA | 561 | 463 | 32 | 34 | 2 | 0.09326 | lowCG | nonCGI | HypoMeth | NA | FALSE |
| Mouse | M189 | Intergenic_HyperMeth_highCG | chr2 | 5083412 | 5083934 | - | TTTTTGGAAGTATTTAATGAGATTAGA | AAAATCCTATAAACACAAACCAAAAA | 522 | 469 | 43 | 36 | 27 | 0.64969 | highCG | CGI | HyperMeth | NA | FALSE |
| Mouse | M190 | Intergenic_HyperMeth_highCG | chr3 | 34959111 | 34959644 | + | TTGTGAGTAGAGAGATATTTGTTTTA | AACAATTTTCCCTCCCATTATAAAAA | 533 | 421 | 22 | 32 | 7 | 0.59598 | highCG | CGI | HyperMeth | NA | FALSE |
| Mouse | M191 | Intergenic_HyperMeth_highCG | chr1 | 54957181 | 54957634 | - | TAGAGATTTATAAGGGTGGTGTGATA | CAAATATCTACCACTTACCCACAC | 453 | 394 | 18 | 33 | 6 | 0.60985 | highCG | CGI | HyperMeth | NA | FALSE |
| Mouse | M192 | Intergenic_HyperMeth_highCG | chr2 | 121044505 | 121044973 | - | ATTTGAGAGTTAGTGAATTTTGAAGT | AATCTTTAAAAATCTCCCACCTATTC | 468 | 387 | 31 | 39 | 7 | 0.58927 | highCG | CGI | HyperMeth | NA | FALSE |
| Mouse | M193 | Intergenic_HyperMeth_highCG | chr11 | 6447229 | 6447753 | - | AGTGAATTTAAATAAGGAATAGGTAT | AAACTCCCAAAACTCATCTATAA | 524 | 449 | 34 | 40 | 19 | 0.70313 | highCG | CGI | HyperMeth | NA | FALSE |
| Mouse | M207 | Intergenic_HyperMeth_mediumCG | chr12 | 107648033 | 107648592 | + | TTGAATGGGTGATAAGGAATTGG | ATCCATTCTCTCTCCCACTATATTTT | 559 | 494 | 30 | 31 | 6 | 0.36961 | mediumCG | nonCGI | HyperMeth | NA | FALSE |
| Mouse | M210 | Intergenic_HyperMeth_mediumCG | chr1 | 161233226 | 161233794 | - | TTTGGATTTTAGTTTTTGATAGGTGA | TATTTTTCCTTTAAATTCCACAAACA | 568 | 485 | 34 | 34 | 7 | 0.40289 | mediumCG | nonCGI | HyperMeth | NA | FALSE |
| Mouse | M215 | Intergenic_HyperMeth_mediumCG | chr1 | 193720200 | 193720759 | + | TTTTTGATTATTGAGGAGGTGAGATA | TCCCTTAATCTCTCTCTCCAAAAA | 559 | 455 | 28 | 32 | 6 | 0.29784 | mediumCG | nonCGI | HyperMeth | NA | FALSE |
| Mouse | M206 | Intergenic_HyperMeth_lowCG | chr9 | 98640215 | 98640782 | - | GAGTGGTGGGAAATGATTTTTG | TCAATATCTAAACCCAATCAAAAAC | 567 | 516 | 44 | 31 | 3 | 0.16282 | lowCG | nonCGI | HyperMeth | NA | FALSE |
| Mouse | M208 | Intergenic_HyperMeth_lowCG | chr7 | 36492062 | 36492626 | + | TTTAGGTTTGAAGGTTTGATAAAAGA | ACCATATCTATAATAAAACTCCAAAATT | 564 | 486 | 32 | 34 | 6 | 0.21903 | lowCG | nonCGI | HyperMeth | NA | FALSE |
| Mouse | M209 | Intergenic_HyperMeth_lowCG | chr3 | 31767306 | 31767865 | + | ATTAGGAGATTTTGAAGTGAAGAAAG | AATATAACAAAAATCTACAAATACCACA | 559 | 457 | 36 | 30 | 2 | 0.12655 | lowCG | nonCGI | HyperMeth | NA | FALSE |
| Mouse | M211 | Intergenic_HyperMeth_lowCG | chr12 | 5552296 | 5552855 | + | GGTAATGAATGTGGATAGGGATATTT | CCCATTTATATATACAAATAATAACACCAC | 559 | 458 | 37 | 35 | 2 | 0.16471 | lowCG | nonCGI | HyperMeth | NA | FALSE |
| Mouse | M212 | Intergenic_HyperMeth_lowCG | chr5 | 28136357 | 28136936 | + | TAGAATTTTTGGTGATATTATTGGGG | CCCCACACCATCTCTATAAATTTTT | 579 | 508 | 50 | 32 | 1 | 0.06883 | lowCG | nonCGI | HyperMeth | NA | FALSE |
| Mouse | M213 | Intergenic_HyperMeth_lowCG | chr2 | 178895007 | 178895571 | - | TGAATAGGTGGAAGATGATTGAATAT | CCTAAACTAAAAACACCAATTATCAC | 564 | 475 | 34 | 33 | 0 | 0.06526 | lowCG | nonCGI | HyperMeth | NA | FALSE |
| Mouse | M214 | Intergenic_HyperMeth_lowCG | chr2 | 179281211 | 179281781 | + | ATGTTTGTGTGAAGTTTATATATTGAT | CATAAACCACCCTCTCTACTAAAA | 570 | 469 | 40 | 29 | 2 | 0.16340 | lowCG | nonCGI | HyperMeth | NA | FALSE |
| Mouse | M216 | Intergenic_HyperMeth_lowCG | chr1 | 118788418 | 118788977 | - | TTTGAGTGTGGGAGAAAATATTATTTA | CAAACCCAAAAACATAAAATCTCTTA | 559 | 475 | 39 | 31 | 4 | 0.16867 | lowCG | nonCGI | HyperMeth | NA | FALSE |
| Mouse | M217 | Intergenic_HyperMeth_lowCG | chr5 | 125819402 | 125819973 | + | TGATATTTTATAATTAGGTGTTTGTGTG | TCATCATACCAAAACTAACACCTATA | 571 | 489 | 31 | 34 | 5 | 0.21308 | lowCG | nonCGI | HyperMeth | NA | FALSE |
| Mouse | M218 | Enhancer_HypoMeth_highCG | chr15 | 96709361 | 96709847 | - | TTAGTGGAAAAATTTTAGGAGATTT | TCCTAAAATAAATAAATAATAATAATCCAA | 486 | 420 | 39 | 33 | 8 | 0.71718 | highCG | CGI | HypoMeth | NA | FALSE |
| Mouse | M219 | Enhancer_HypoMeth_highCG | chr15 | 101977097 | 101977671 | + | GGTGTTGTGGATATTTAAGAATTTA | AAACTATTCCAAAAATCCAAATC | 574 | 503 | 52 | 31 | 4 | 0.52707 | highCG | CGI | HypoMeth | NA | TRUE |
| Mouse | M220 | Enhancer_HypoMeth_highCG | chr17 | 83276123 | 83276663 | - | TAATAGTTTGGGAATTAGGTTAAGGA | ATATCCTCAATCAAAAACCAACAA | 540 | 468 | 46 | 34 | 10 | 0.81889 | highCG | CGI | HypoMeth | NA | FALSE |
| Mouse | M227 | Enhancer_HypoMeth_highCG | chr18 | 24466748 | 24467312 | - | TGAGGTGTGGAAGTAAAAATAATTAT | AAACCTTAAAATCCTAATATCTAATTTTT | 564 | 460 | 29 | 36 | 10 | 0.88723 | highCG | CGI | HypoMeth | NA | FALSE |
| Mouse | M228 | Enhancer_HypoMeth_highCG | chr19 | 44388419 | 44388949 | + | TTTGTTTATTGTTTGTTTAGTGTAGG | TTTTTCCTAAATCAAAAACAATATAAAAA | 530 | 468 | 36 | 33 | 9 | 0.63210 | highCG | nonCGI | HypoMeth | NA | FALSE |
| Mouse | M221 | Enhancer_HypoMeth_mediumCG | chr12 | 72844024 | 72844583 | + | GGGTTGGATTTAGGGGTTTTATATAT | AAAATTATAACACAAAAACACTACTCC | 559 | 503 | 41 | 35 | 7 | 0.27840 | mediumCG | nonCGI | HypoMeth | NA | FALSE |
| Mouse | M222 | Enhancer_HypoMeth_mediumCG | chr17 | 50470358 | 50470918 | + | AGTTTGAAAAGATTTAAGTTTGAAAAGA | TTTCTCAACAAAAATCTCAAATTACT | 560 | 476 | 35 | 36 | 5 | 0.49734 | mediumCG | nonCGI | HypoMeth | NA | FALSE |
| Mouse | M223 | Enhancer_HypoMeth_mediumCG | chr11 | 45991522 | 45992099 | - | GTAATGAGGTTTAGGTAGGAAGTAAG | TAAACTCTCCTTCAAAAATCCAAATT | 577 | 469 | 35 | 32 | 8 | 0.47093 | mediumCG | nonCGI | HypoMeth | NA | FALSE |
| Mouse | M224 | Enhancer_HypoMeth_mediumCG | chr13 | 60060126 | 60060673 | - | TTTTAATTTTGGGGATTTATGGAATT | AACCCATTATTACCTCCATAAATTTT | 547 | 487 | 30 | 34 | 5 | 0.37802 | mediumCG | nonCGI | HypoMeth | NA | FALSE |
| Mouse | M225 | Enhancer_HypoMeth_mediumCG | chr18 | 70564976 | 70565484 | + | AAGGAAAGTGATTTTTGGGTTTTT | AAATCACACCCAAATTTTCCTAA | 508 | 408 | 28 | 37 | 2 | 0.32330 | mediumCG | nonCGI | HypoMeth | NA | TRUE |
| Mouse | M226 | Enhancer_HypoMeth_mediumCG | chr13 | 99152453 | 99152954 | - | GTGAATAGTTTGTGTTGGATTAATAT | TAACTCACACCAAAAATAAAAACTCA | 501 | 445 | 40 | 31 | 7 | 0.39824 | mediumCG | nonCGI | HypoMeth | NA | FALSE |
| Mouse | M236 | ICR_Mat_HypoMeth | chr7 | 6729294 | 6729873 | + | ATTAAGATTGAGTGAGGGTTTAG | ACCACCACAATAAACATCAAAAT | 579 | 533 | 49 | 29 | 5 | 0.60469 | highCG | CGI | HypoMeth | NA | FALSE |
| Mouse | M237 | ICR_Mat_HypoMeth | chr7 | 6728328 | 6728788 | + | GGTGGAATTGGTTTTTAATTAATTAT | TTTATAATCCTAAATCACCACAAA | 460 | 401 | 34 | 26 | 1 | 0.33699 | mediumCG | nonCGI | HypoMeth | NA | FALSE |
| Mouse | M238 | ICR_Mat_HypoMeth | chr2 | 174295865 | 174296399 | + | AGAGGGATATGTTAGATTTTAGAGAA | TTTATCCCCTCCTCCTTCTATTATAA | 534 | 474 | 45 | 40 | 5 | 0.73695 | highCG | nonCGI | HypoMeth | Gnasas1 | FALSE |
| Mouse | M239 | ICR_Mat_HypoMeth | chr2 | 174300331 | 174300788 | + | GAAATTTTGTTAGAAGTTTTAGG | ATAACCTATTTTAAAATACAAATATAAAA | 457 | 385 | 29 | 40 | 11 | 0.70621 | highCG | CGI | HypoMeth | NA | FALSE |

Supplementary Table 1 (cont.)

| Organism | ID | Class | Chromosome | Start | End | Strand | Fwseq | Rvseq | Full frag. length | GCH inform. length | GCH number | GCH MaxGap | WCG number | obs/exp CG | o/e CG level | CGI class | Methylation class | Promoter SYMBOL | Edges |
|---|---|---|---|---|---|---|---|---|---|---|---|---|---|---|---|---|---|---|---|
| Mouse | M240 | ICR_Pat_HyperMeth | chr9 | 89872761 | 89873250 | - | TGAGTTTGTTTGGTTAATTTAATTTA | ACTTCTTCAAAAATTTAATCTTTTAT | 489 | 401 | 25 | 37 | 1 | 0.10028 | lowCG | nonCGI | HyperMeth | NA | FALSE |
| Mouse | M241 | ICR_Pat_HyperMeth | chr7 | 142582311 | 142582827 | + | AGATTTAGTATTGTTTGGTTATTAGTT | CTTTCTCCACCACTTATCTAAATTC | 516 | 460 | 32 | 39 | 2 | 0.09836 | lowCG | nonCGI | HyperMeth | NA | FALSE |
| Mouse | M242 | ICR_Pat_HyperMeth | chr12 | 109526490 | 109526991 | + | TATGTGTATTTTTGGGTGGTAGTATA | AATACATCCTATTCCCTAATAAAAATTT | 501 | 425 | 35 | 38 | 9 | 0.57806 | highCG | nonCGI | HyperMeth | NA | FALSE |
| Mouse | M243 | ICR_Pat_HyperMeth | chr12 | 109531102 | 109531599 | - | GGTTTAGTTGGTGATAAAGATGATAG | CACACACACACTAAATTCACAATT | 497 | 445 | 25 | 39 | 3 | 0.10363 | lowCG | nonCGI | HyperMeth | NA | FALSE |
| Mouse | M153 | CTCF_HypoMeth | chr2 | 178407300 | 178407831 | - | GGTGGTGGAGTTTTATAGTGATT | CTCACAAACTCCTCCCTATATAATT | 531 | 483 | 36 | 29 | 7 | 0.64193 | highCG | CGI | HypoMeth | Sycp2 | FALSE |
| Mouse | M180 | CTCF_HypoMeth | chr7 | 79495791 | 79496324 | + | TTTTTGAAATTTGTAGAGATGAGGAG | AACACAAATATTCCTAAAAATCTCCC | 533 | 440 | 33 | 38 | 4 | 0.60200 | highCG | CGI | HypoMeth | NA | TRUE |
| Mouse | M185 | CTCF_HypoMeth | chr4 | 55480237 | 55480759 | - | ATTGAGTGATGGTGAAAGGAAG | CCTCTCAAATCTAAACCTTCAAATC | 522 | 455 | 40 | 36 | 9 | 0.51457 | highCG | CGI | HypoMeth | NA | FALSE |
| Mouse | M229 | CTCF_HypoMeth | chr2 | 119337717 | 119338251 | + | TTAGTTAGGGAATTGTGATTGTTTTT | ATTTAAAATTCCTAAACTAATCCCCA | 534 | 467 | 39 | 36 | 9 | 0.74905 | highCG | CGI | HypoMeth | NA | FALSE |
| Mouse | M230 | CTCF_HypoMeth | chr1 | 52630412 | 52630971 | + | TGAAAGGAGGGATAAATAAATAAATAAGT | CCTAAACAAACCAAAACACTAAAATATC | 559 | 495 | 54 | 25 | 8 | 0.74096 | highCG | CGI | HypoMeth | Nemp2 | FALSE |
| Mouse | M231 | CTCF_HypoMeth | chr7 | 38060959 | 38061522 | + | GAATGAAAAAGAGTTTATAGAGGTGA | CTAAATAAAACAATCCAATCACTCTA | 563 | 502 | 53 | 29 | 4 | 0.68520 | highCG | CGI | HypoMeth | NA | TRUE |
| Mouse | M232 | CTCF_HypoMeth | chr11 | 63925115 | 63925605 | - | TTTTTGGGGAAATAGAGAGGATATT | ACCTATTATAACTCCAAACTTTCTAA | 490 | 427 | 45 | 30 | 3 | 0.63954 | highCG | CGI | HypoMeth | NA | FALSE |
| Mouse | M233 | CTCF_HypoMeth | chr19 | 55218327 | 55218799 | - | TTAGTGGAATTTAGTTTGTTTTTGA | AATTCCAACTCATAACCTTATCCA | 472 | 423 | 29 | 34 | 4 | 0.52066 | highCG | nonCGI | HypoMeth | NA | FALSE |
| Mouse | M234 | CTCF_HypoMeth | chr6 | 128799420 | 128799992 | + | TTTAGGAGAAAATTGGGAATTAAAGA | AAACACCCTAAAACCTTATCACA | 572 | 495 | 41 | 33 | 10 | 0.65126 | highCG | CGI | HypoMeth | NA | FALSE |
| Mouse | M235 | CTCF_HypoMeth | chr11 | 78826415 | 78826989 | + | TGTATTTAAGGTTGTAAGGGAGATAA | CCAAATACTCCTAATAACAAAATATTCC | 574 | 509 | 44 | 33 | 11 | 0.79744 | highCG | CGI | HypoMeth | Lyrm9 | TRUE |
| Mouse | Control locus 1 | Promoter_HypoMeth_highCG | chr1 | 71102603 | 71102954 | + | ATTTAGAAAATTTAGGAGTGGGATA | AATCCTTCCATCTCTAAATTTTAC | 351 | 268 | 28 | 31 | 10 | 0.83150 | highCG | CGI | HypoMeth | Vwc2l | NA |
| Mouse | Control locus 2 | Promoter_HypoMeth_highCG | chr1 | 9299278 | 9299634 | + | GGGATAGGGAGGTTGGGTTTTA | TCACTCTATATATTTCAATTCCCCCA | 356 | 292 | 25 | 36 | 5 | 0.66303 | highCG | CGI | HypoMeth | Rrs1 | NA |
| Lambda | L1 | NA | NC_001416.1 | 7386 | 7755 | + | GATATGAAAATGAGGTGGGATTA | ATTTTCAAAATCATCAAAAACACCA | 369 | 299 | 24 | 46 | 14 | NA | NA | NA | NA | NA | NA |

**Supplementary Table 1. List of mouse and Lambda phage amplicons used in this study and their main characteristics.**

**“Organism”:** mouse or lambda, for which amplicons are designed. **“ID”:** amplicon identifier. **“Class”:** amplicon category. **“Chromosome”**, **“Start”** and **“End”** refer to the coordinates of mouse (mm10) and Lambda phage (NC_001416.1) genome assemblies. **“Strand”**: DNA strand amplified. **“Fwseq”** and **“Rvseq”**: sequences of the forward and reverse primers, respectively. **“Full frag. length”:** the full length of amplicon fragment refers to the total number of base pairs covered by the amplicon. **“GCH inform. length”**: the GCH-informative length refers to the number of base pairs covered from the first to the last GCH trinucleotide. **“GCH number”**: number of GCH trinucleotides contained in the amplicon sequence. **“GCH MaxGap”**: maximum number of base pairs separating adjacent GCH trinucleotides. **“WCG number”**: number of WCG trinucleotides contained in the amplicon sequence. **“obs/exp CG”**: Observed/Expected CG ratio of the amplicon. **“o/e CG level”**: classification of CG levels based on Observed/Expected CG ratio of the amplicon: “highCG”: ≥ 0.5 observed/expected CG; “mediumCG”: <0.5 and ≥ 0.25 observed/expected CG, and “lowCG”: <0.25 observed/expected CG. The classification is based on all CGs within an amplicon irrespective of the identity of adjacent bases. **“CGI class”**: whether amplicon overlaps a CGI (CGI) or not (nonCGI), based on the UCSC Table Browser. **“Methylation class”**: methylation status of the amplicon. **“Promoter SYMBOL”**: name of gene(s) overlapping promoter amplicons (TSS+/- 500bp of reference genes). **“Edges”**: whether NOMe-seq footprints in round spermatids are at the edges of the amplicon. If TRUE, amplicons were not included in footprint occupancy analysis.

### Supplementary Table 2

| Organism | ID | Class | Chromosome | Start | End | Strand | Fwseq | Rvseq | Full frag. length | GCH inform. length | GCH number | GCH MaxGap | WCG number | obs/exp CG | o/e CG level | CGI class | Methylation class | Promoter SYMBOL |
|---|---|---|---|---|---|---|---|---|---|---|---|---|---|---|---|---|---|---|
| Human | H1 | Promoter_HypoMeth_highCG | chr11 | 94473365 | 94473944 | - | TAGATTTTTAGGGAGTGATAAAT | ATTTAAATCTCCTCCTACCTTT | 579 | 531 | 44 | 31 | 44 | 0.99104 | highCG | CGI | HypoMeth | AMOTL1 |
| Human | H2 | Promoter_HypoMeth_highCG | chr8 | 133787403 | 133787975 | - | GAAAGAGAGATATTAGAATGGGG | ATCAAATACTTCACAACTACCAATAA | 572 | 511 | 66 | 28 | 66 | 0.92904 | highCG | CGI | HypoMeth | PHF20L1 |
| Human | H3 | Promoter_HypoMeth_highCG | chr7 | 20370975 | 20371544 | - | AGTTTAAGAATAAGTGAAAATATTTAG | AAATATACATTAAAATAATTTCCCCC | 569 | 513 | 55 | 30 | 55 | 0.89777 | highCG | CGI | HypoMeth | ITGB8 |
| Human | H4 | Promoter_HypoMeth_highCG | chr4 | 109683665 | 109684241 | - | TAGATTTAAAAGGGAGATGATTTTA | CATCTCCCATCCCTAAAATAATAT | 576 | 525 | 56 | 27 | 56 | 0.80485 | highCG | CGI | HypoMeth | ETNPPL |
| Human | H5 | Promoter_HypoMeth_highCG | chr4 | 170678792 | 170679366 | - | TAGTAGATATTTGAAGAGGAAGGTTG | TACTTTCCTAATCCAACCCCCT | 574 | 466 | 43 | 31 | 43 | 0.79647 | highCG | CGI | HypoMeth | HPF1 |
| Human | H6 | Promoter_HypoMeth_highCG | chr8 | 25902825 | 25903404 | + | TAAAAAGAAGGGGAAGATTTGG | ATATCACCATAAAACTCCAAAAA | 579 | 524 | 47 | 33 | 47 | 0.7953 | highCG | CGI | HypoMeth | PPP2R2A |
| Human | H7 | Promoter_HypoMeth_highCG | chr15 | 40226090 | 40226669 | - | GTGGAATTTAAGGGATGAAAAG | TCATTCCTATCTATAATTCATTTCTT | 579 | 509 | 52 | 25 | 52 | 0.77357 | highCG | CGI | HypoMeth | EIF2AK4 |
| Human | H8 | Promoter_HypoMeth_highCG | chr5 | 110847891 | 110848470 | - | AATTTTAGATATTTAGGGAATTTTG | TTCTCTTTCATCTATAAAATCCTC | 579 | 507 | 42 | 33 | 42 | 0.73298 | highCG | CGI | HypoMeth | STARD4-AS1 |
| Human | H9 | Promoter_HypoMeth_highCG | chr6 | 84569166 | 84569736 | - | TTGTAGATAAAGTGGGATTAGAAAGA | AACAAAACTAAAAATTTATTCCCACA | 570 | 506 | 46 | 33 | 46 | 0.72377 | highCG | CGI | HypoMeth | CYB5R4 |
| Human | H10 | Promoter_HypoMeth_highCG | chr6 | 72595990 | 72596557 | + | TTAGAAGAGGGAGGATGATTTAT | AAAACCCAATCTTTCATAATTATTT | 567 | 488 | 54 | 35 | 54 | 0.71634 | highCG | CGI | HypoMeth | RIMS1 |
| Human | H11 | Promoter_HypoMeth_highCG | chr9 | 14993005 | 14993583 | + | TAGTAGATATTTGAAGAGGAAGGTTG | TACTTTCCTAATCCAATCCCCTA | 578 | 463 | 42 | 30 | 42 | 0.7115 | highCG | CGI | HypoMeth | LOC389705 |
| Human | H12 | Promoter_HypoMeth_highCG | chr2 | 208889866 | 208890444 | - | TGTGTATAAGAGGAATTGTATATAAGAA | TACTTTCTCCTCCTATTACATACCTA | 578 | 509 | 46 | 30 | 46 | 0.67625 | highCG | CGI | HypoMeth | PLEKHM3 |
| Human | H13 | Promoter_HypoMeth_highCG | chr5 | 140743666 | 140744233 | + | AGTTATTGGTTAGGATTTTGAG | TTCTCAACCAAAATATTAAAATTCAC | 567 | 503 | 43 | 33 | 43 | 0.67008 | highCG | CGI | HypoMeth | PCDHGB2 |
| Human | H14 | Promoter_HypoMeth_highCG | chr5 | 101631845 | 101632422 | + | TTTTTGGGGGTTAGAGGATAAG | AAACTCCCACAATCTCTATATATTCC | 577 | 525 | 44 | 36 | 44 | 0.66397 | highCG | CGI | HypoMeth | SLCO4C1 |
| Human | H15 | Promoter_HypoMeth_highCG | chr3 | 134082776 | 134083352 | - | TTTTTGAGGAGTTTGGGGAGGA | CTACCAACAAAACCCTAATAAAACCT | 576 | 522 | 51 | 30 | 51 | 0.65526 | highCG | CGI | HypoMeth | AMOTL2 |
| Human | H16 | Promoter_HypoMeth_highCG | chr11 | 4629075 | 4629654 | - | AGAAAAATTTAGGAAGTAGAGAAAAA | AAAACTTCCAATTAAAACCCCA | 579 | 521 | 52 | 29 | 52 | 0.62384 | highCG | CGI | HypoMeth | TRIM68 |
| Human | H17 | Promoter_HypoMeth_highCG | chr20 | 524003 | 524571 | - | AGAATTGAATTTTTAAGGTTATAATAGGA | CTCCTCCAAATTATCCCCTAATC | 568 | 484 | 46 | 36 | 46 | 0.59928 | highCG | CGI | HypoMeth | CSNK2A1 |
| Human | H18 | Promoter_HypoMeth_highCG | chr2 | 85581229 | 85581800 | - | GGAGGGTTAATTATTATTTGAGA | AATAACCACACCCTTAAAACTA | 571 | 511 | 46 | 34 | 46 | 0.53343 | highCG | CGI | HypoMeth | RETSAT |
| Human | H19 | Promoter_HypoMeth_mediumCG | chr7 | 53879281 | 53879859 | + | TAGAGAGTGGTAGTGAAAATTT | AAAACTCAATAACAAATAAATATTCA | 578 | 505 | 44 | 30 | 44 | 0.46394 | mediumCG | nonCGI | HypoMeth | LINC01446 |
| Human | H20 | Promoter_HypoMeth_mediumCG | chr7 | 30978023 | 30978592 | - | GTGAAATTTTTAGGGTTTAGTTTTAT | ATCATTATTAAAAATATCCCCAAAAC | 569 | 485 | 49 | 28 | 49 | 0.34638 | mediumCG | nonCGI | HypoMeth | AQP1 |
| Human | H21 | Promoter_HypoMeth_mediumCG | chr19 | 11649341 | 11649919 | + | AGAGATAAAGAGAGATGATTTTAGAGA | TCTCAATTTCCAAATCACATAAAATAA | 578 | 506 | 44 | 35 | 44 | 0.29239 | mediumCG | nonCGI | HypoMeth | CNN1 |
| Human | H22 | Promoter_HypoMeth_lowCG | chr1 | 8484607 | 8485185 | + | TAGTGTGAGGAGTGATTTTTGTTATT | TCTTTAACTAATCTATCCCTAATCTACA | 578 | 489 | 45 | 37 | 45 | 0.23643 | lowCG | CGI | HypoMeth | RERE |
| Human | H23 | Promoter_HypoMeth_lowCG | chr15 | 40731473 | 40732043 | - | ATAGAGAGAGATGGGAAGTTTTTG | CCCTCCTTTAAATATATCCCCC | 570 | 502 | 44 | 31 | 44 | 0.21509 | lowCG | nonCGI | HypoMeth | BAHD1 |
| Human | H24 | Promoter_HyperMeth_highCG | chr19 | 6389355 | 6389914 | + | GAATTTGAGGTTTAGAGAGGGTATT | ATAATCACCAAATAATCTTTTTCTCTTTA | 559 | 460 | 38 | 36 | 38 | 0.5582 | highCG | nonCGI | HyperMeth | GTF2F1 |
| Human | H25 | Promoter_HyperMeth_mediumCG | chr2 | 109605473 | 109606052 | - | TGAGAATGTGAATAGATATGTTGGAA | CAAATACCATCCAATTCTAAATAACAA | 579 | 526 | 44 | 32 | 44 | 0.32716 | mediumCG | nonCGI | HyperMeth | EDAR |
| Human | H26 | Promoter_HyperMeth_mediumCG | chr22 | 19118386 | 19118951 | + | GGTGATTGTTAGGGATAATTGTTT | ATATAACCCTTCTTCCTTAAAACTAT | 565 | 496 | 41 | 34 | 41 | 0.30695 | mediumCG | CGI | HyperMeth | TSSK2 |
| Human | H27 | Promoter_HyperMeth_mediumCG | chr9 | 130616795 | 130617372 | + | GGGGTTAGGAGAAGTGGATATAG | CCATCTTCTTAAACCAATAATCTCAA | 577 | 493 | 41 | 37 | 41 | 0.29359 | mediumCG | CGI | HyperMeth | ENG |
| Human | H28 | Promoter_HyperMeth_mediumCG | chr12 | 53207498 | 53208061 | - | GATTTTTGTAGGTGTGTTTAGTGATA | TTAATAATAACCTCCTAAATTCCCCC | 563 | 509 | 49 | 31 | 49 | 0.28609 | mediumCG | nonCGI | HyperMeth | KRT4 |
| Human | H29 | Promoter_HyperMeth_lowCG | chr16 | 2513760 | 2514334 | + | AGAGTGAAGATATATTATTTGTTAAAAGGA | CCCATCCTCCTAAATCCTAAAAA | 574 | 494 | 43 | 27 | 43 | 0.24742 | lowCG | nonCGI | HyperMeth | TEDC2 |
| Human | H30 | Promoter_HyperMeth_lowCG | chr10 | 45406387 | 45406954 | - | TAGGGGTGAGGATAAAGTTTTG | AAAACTCATTTCAAAAATTATTTCTACT | 567 | 484 | 54 | 31 | 54 | 0.24343 | lowCG | nonCGI | HyperMeth | TMEM72-AS1 |
| Human | H31 | Promoter_HyperMeth_lowCG | chr22 | 42348019 | 42348585 | + | ATAGTATTAGGGATTTAGAAGAG | AACAAAATAAAAATAAAATTACCAAA | 566 | 493 | 48 | 24 | 48 | 0.22785 | lowCG | nonCGI | HyperMeth | SMIM45 |
| Human | H32 | Promoter_HyperMeth_lowCG | chr1 | 153329847 | 153330421 | - | GTTAGGGAAGGAAAGAAGGAAT | ATCTCACCCTATAAATCTAAAAACAC | 574 | 495 | 47 | 35 | 47 | 0.21443 | lowCG | nonCGI | HyperMeth | S100A9 |
| Human | H33 | Promoter_HyperMeth_lowCG | chr15 | 43558977 | 43559550 | - | ATTTAATGTTTAGGAAGTGATTTTT | TACAATCTCTCTCTATCCTTCTTAT | 573 | 493 | 44 | 31 | 44 | 0.20463 | lowCG | nonCGI | HyperMeth | TGM5 |
| Human | H34 | Promoter_HyperMeth_lowCG | chr15 | 42158434 | 42159002 | + | GGGAGAGAGGGGATATAGGATAT | CCTTTATATAAAAACCTCACCAATATCC | 568 | 511 | 53 | 30 | 53 | 0.17283 | lowCG | nonCGI | HyperMeth | MIR4310 |
| Human | H35 | Promoter_HyperMeth_lowCG | chr9 | 139869292 | 139869859 | + | TTTTAGGAAAAGGTTGGGGTTAG | ACTCCTCAATCCCTATAATCAAAATA | 567 | 510 | 46 | 35 | 46 | 0.16263 | lowCG | nonCGI | HyperMeth | PTGDS |
| Human | H36 | Promoter_HyperMeth_lowCG | chr1 | 20445617 | 20446184 | + | TTTTATTTAGTGGGAGAGGTAGGG | AACAAAACAAAAACAAAACAAAACTC | 567 | 475 | 45 | 27 | 45 | 0.08557 | lowCG | nonCGI | HyperMeth | PLA2G2D |
| Human | H37 | Promoter_HyperMeth_lowCG | chr3 | 50316051 | 50316614 | + | ATTGGGATTTAGTTGTTTATTATTTTT | CCCTCTCCTACCTTCTTAAATC | 563 | 486 | 50 | 27 | 50 | 0.04234 | lowCG | nonCGI | HyperMeth | LSMEM2 |
| Human | H38 | Intragenic_HypoMeth_highCG | chr6 | 34494806 | 34495385 | - | AATGTTTTGTGTTTGGTAGATGG | CCAAATCCAAATTCTAAAAATCCTTC | 579 | 526 | 55 | 33 | 55 | 0.83376 | highCG | CGI | HypoMeth | PACSIN1 |
| Human | H39 | Intragenic_HypoMeth_highCG | chr5 | 115783047 | 115783611 | - | TGTTGTTATTTATTATTGTTATTAATTAAT | TTAATAAAATTCTAATTCCTCTCCC | 564 | 493 | 43 | 34 | 43 | 0.71535 | highCG | CGI | HypoMeth | SEMA6A |
| Human | H40 | Intragenic_HypoMeth_highCG | chr16 | 30543885 | 30544464 | - | TAGAAATAAGGAAGAGGAAAGATAAA | AACAAAAATCAAAATCACAAACA | 579 | 516 | 52 | 36 | 52 | 0.68452 | highCG | CGI | HypoMeth | ZNF747 |
| Human | H41 | Intragenic_HypoMeth_highCG | chr19 | 1789406 | 1789983 | - | ATTTTGGAGATTTATTGGGAAA | ACCACCTAATAATACTTCTTAACC | 577 | 494 | 47 | 36 | 47 | 0.65245 | highCG | CGI | HypoMeth | ATP8B3 |
| Human | H42 | Intragenic_HypoMeth_highCG | chr5 | 77805479 | 77806045 | + | GGAAATTTGAATAGATAGAAAGTGGT | ACATCAATATATATCATATCATTATCACCT | 566 | 486 | 48 | 34 | 48 | 0.65093 | highCG | CGI | HypoMeth | LHFPL2 |
| Human | H43 | Intragenic_HypoMeth_highCG | chr2 | 132431282 | 132431861 | - | TGAATGATTATGTTTTTATGTTTAATTT | CACCCTAAAATCTAATCTTCCTT | 579 | 505 | 54 | 27 | 54 | 0.57924 | highCG | CGI | HypoMeth | LINC03124 |
| Human | H44 | Intragenic_HypoMeth_highCG | chr7 | 158750066 | 158750645 | + | AGTGTGTTGTGGATAGATTATTTTT | CAAAATCCCACAAACACTAAAAA | 579 | 456 | 41 | 33 | 41 | 0.57692 | highCG | CGI | HypoMeth | DYNC2I1 |
| Human | H45 | Intragenic_HypoMeth_mediumCG | chr11 | 134158448 | 134159010 | + | TAGTGTGAGAATTAGAGGGAAATTT | TCCAATCAACAATAAACTATTAATAACT | 562 | 470 | 40 | 37 | 40 | 0.45346 | mediumCG | nonCGI | HypoMeth | GLB1L3 |
| Human | H46 | Intragenic_HypoMeth_mediumCG | chr7 | 157475400 | 157475959 | - | TTAGAAGAAGGGATGGATATTT | AAAAATCTCCTCTTACCTAATAA | 559 | 506 | 40 | 32 | 40 | 0.43348 | mediumCG | nonCGI | HypoMeth | PTPRN2 |

### Supplementary Table 2 (cont.)

| Organism | ID | Class | Chromosome | Start | End | Strand | Fwseq | Rvseq | Full frag. length | GCH inform. length | GCH number | GCH MaxGap | WCG number | obs/exp CG | o/e CG level | CGI class | Methylation class | Promoter SYMBOL |
|---|---|---|---|---|---|---|---|---|---|---|---|---|---|---|---|---|---|---|
| Human | H47 | Intragenic_HyperMeth_highCG | chr11 | 1215508 | 1216087 | - | TTTGGGATAGTTTTTATGGTAT | AAATCCATCATCCTAAAATACC | 579 | 513 | 45 | 34 | 45 | 0.76316 | highCG | CGI | HyperMeth | MUC5B |
| Human | H48 | Intragenic_HyperMeth_highCG | chr19 | 3192207 | 3192786 | + | AAGATTTTTAGGAGGTTTGAGA | AAAAACAATCCATCCTACTTTC | 579 | 514 | 47 | 32 | 47 | 0.65297 | highCG | CGI | HyperMeth | NCLN |
| Human | H49 | Intragenic_HyperMeth_highCG | chr16 | 1250207 | 1250786 | - | GATGAAGTTGTAGAATGAGTGG | TCCCTAACCCTAATTATACCTTTTA | 579 | 500 | 39 | 33 | 39 | 0.62155 | highCG | CGI | HyperMeth | CACNA1H |
| Human | H50 | Intragenic_HyperMeth_highCG | chr11 | 69468643 | 69469222 | + | AAGTTTTGGGTATGTTTAATTTGTTA | TCTAAACTTTCATATTTATCTTTTTATCT | 579 | 475 | 39 | 32 | 39 | 0.61548 | highCG | CGI | HyperMeth | CCND1 |
| Human | H51 | Intragenic_HyperMeth_highCG | chr9 | 96278125 | 96278692 | - | TTATTTGAGTATAGATTGATTATTATTT | ATAAATTCCTAACCTCTTAAATTAAT | 567 | 496 | 41 | 36 | 41 | 0.61481 | highCG | CGI | HyperMeth | FAM120A |
| Human | H52 | Intragenic_HyperMeth_highCG | chr12 | 132837140 | 132837711 | + | TTTATTTTATTTGATTTGAGGTTTTT | AATAAAACTCAAATTCAATCTAAACC | 571 | 479 | 42 | 33 | 42 | 0.60174 | highCG | CGI | HyperMeth | GALNT9 |
| Human | H53 | Intragenic_HyperMeth_highCG | chr3 | 126668048 | 126668627 | + | GGAGAATAGAAGGGATAATTTGATTT | TTCCCTTCATATCCAAAACAATCT | 579 | 516 | 50 | 25 | 50 | 0.59535 | highCG | CGI | HyperMeth | CHCHD6 |
| Human | H54 | Intragenic_HyperMeth_highCG | chr16 | 87742695 | 87743274 | - | AGGGATTTAAGTTTGAAAAGAAG | CTTTCCACAACAAAATCCTATT | 579 | 508 | 53 | 30 | 53 | 0.59337 | highCG | CGI | HyperMeth | KLHDC4 |
| Human | H55 | Intragenic_HyperMeth_highCG | chr10 | 135271644 | 135272220 | - | AAGGGATTTGATAGAGAATAAG | CCCTAACCCTAACCTTAATAAT | 576 | 532 | 53 | 28 | 53 | 0.55994 | highCG | CGI | HyperMeth | SCART1 |
| Human | H56 | Intragenic_HyperMeth_highCG | chr16 | 15026516 | 15027085 | - | TAGGAAGAGATAGATGAGGAGA | AACAATCTAAACATCTTCCAAAT | 569 | 508 | 48 | 35 | 48 | 0.55725 | highCG | CGI | HyperMeth | NPIPA1 |
| Human | H57 | Intragenic_HyperMeth_highCG | chr15 | 93588350 | 93588923 | + | AGTTTGTTTTTGTTGGAGTGGA | TCAAAAACTTCCAAAAATATATAAACCA | 573 | 510 | 42 | 32 | 42 | 0.54729 | highCG | CGI | HyperMeth | RGMA |
| Human | H58 | Intragenic_HyperMeth_highCG | chr11 | 2593875 | 2594442 | + | TTTATTTTGGGGGTAATTTTTATT | TAAAACCATTCAAAACTTCAAAATAA | 567 | 512 | 42 | 35 | 42 | 0.54361 | highCG | CGI | HyperMeth | KCNQ1 |
| Human | H59 | Intragenic_HyperMeth_highCG | chr16 | 15221909 | 15222478 | + | TAGTTATTGATTAGGAAGAAGGT | ATCACAAATTACAATTCTTTCCATAT | 569 | 495 | 48 | 33 | 48 | 0.54147 | highCG | CGI | HyperMeth | PDXDC1 |
| Human | H60 | Intragenic_HyperMeth_highCG | chr7 | 1536576 | 1537154 | - | GTTTTTAGAGAGGGGAATTAGATGT | CTCATAAATCCTCACACAAAACAAA | 578 | 521 | 46 | 30 | 46 | 0.52732 | highCG | CGI | HyperMeth | INTS1 |
| Human | H61 | Intragenic_HyperMeth_highCG | chr1 | 16474710 | 16475285 | + | TTAATAAGGAAATTGATGTTTGGGAA | CTATTCACCAAAATTAACACCATT | 575 | 485 | 43 | 29 | 43 | 0.52525 | highCG | CGI | HyperMeth | EPHA2 |
| Human | H62 | Intragenic_HyperMeth_mediumCG | chr20 | 49225824 | 49226403 | - | TGGATAGGTAGTGATTGTTATTATG | ATTACCTTTCATCCTAATATAAAACT | 579 | 507 | 63 | 28 | 63 | 0.47407 | mediumCG | CGI | HyperMeth | RIPOR3 |
| Human | H63 | Intragenic_HyperMeth_mediumCG | chr21 | 44839961 | 44840538 | + | TTGGGTGATTGAGAATATTTTT | AATTCTCTATATAATAAACACAAAATAT | 577 | 511 | 50 | 24 | 50 | 0.46802 | mediumCG | CGI | HyperMeth | SIK1 |
| Human | H64 | Intragenic_HyperMeth_mediumCG | chr19 | 33878622 | 33879201 | + | TTTGTTTTTATTGAGGGGTGAG | CAAAAACACTCAAAATCCAAAAATAT | 579 | 529 | 49 | 32 | 49 | 0.45523 | mediumCG | CGI | HyperMeth | PEPD |
| Human | H65 | Intragenic_HyperMeth_mediumCG | chr2 | 27372726 | 27373305 | + | TGGAGTAGTAGAGAGAATATAGGATT | CATTCCAATCCCTTTCTCCAAA | 579 | 520 | 53 | 35 | 53 | 0.44347 | mediumCG | CGI | HyperMeth | TCF23 |
| Human | H66 | Intragenic_HyperMeth_mediumCG | chr5 | 1253511 | 1254090 | + | AATTTGGGGATGGATTATTTTTATGT | ATATCTATCCCTTCACTAAAATTCCC | 579 | 512 | 48 | 33 | 48 | 0.43981 | mediumCG | CGI | HyperMeth | TERT |
| Human | H67 | Intragenic_HyperMeth_mediumCG | chr20 | 62165193 | 62165772 | - | AATTGTGGGGATTTAGAAATTTT | ACACAAAAATAAACTCTCCTAAAA | 579 | 530 | 50 | 32 | 50 | 0.43346 | mediumCG | CGI | HyperMeth | PTK6 |
| Human | H68 | Intragenic_HyperMeth_mediumCG | chr19 | 13320030 | 13320609 | - | ATTGGAGAGTGAGGATTAGATTTTT | CCTTTCTTCTTCCTTAATATCTCCTC | 579 | 502 | 42 | 35 | 42 | 0.40392 | mediumCG | CGI | HyperMeth | CACNA1A |
| Human | H69 | Intragenic_HyperMeth_mediumCG | chr19 | 18886231 | 18886806 | - | TTAGGAGTGAGTTAAATTTAGGGG | TCATCCATTCATCCATTCATCC | 575 | 516 | 56 | 29 | 56 | 0.39064 | mediumCG | CGI | HyperMeth | CRTC1 |
| Human | H70 | Intragenic_HyperMeth_mediumCG | chr16 | 70506785 | 70507355 | + | GGAGTTTGTTTTGTTTTTATGAATTGATAG | CCTCATAATCCCCTAATCCCCT | 570 | 506 | 56 | 28 | 56 | 0.38844 | mediumCG | CGI | HyperMeth | FCSK |
| Human | H71 | Intragenic_HyperMeth_mediumCG | chr14 | 76957609 | 76958188 | + | TTTTAAGGGGAGGATAGTGAGTT | AAATTTCAATATTTTCCTTTCATCTCT | 579 | 518 | 54 | 30 | 54 | 0.35985 | mediumCG | CGI | HyperMeth | ESRRB |
| Human | H72 | Intragenic_HyperMeth_mediumCG | chr7 | 150553502 | 150554068 | - | TTGTGGTATTGAGGAAGAATTGAT | CCATAAAACAACTAAATTCATCTCCT | 566 | 511 | 51 | 29 | 51 | 0.3041 | mediumCG | CGI | HyperMeth | AOC1 |
| Human | H73 | Intragenic_HyperMeth_mediumCG | chr17 | 42856947 | 42857526 | - | TTTTAGAGAGGTAAGATTTTGGTTTT | CATATCACCACTATCTAACCTCC | 579 | 512 | 47 | 36 | 47 | 0.25752 | mediumCG | CGI | HyperMeth | ADAM11 |
| Human | H74 | Intragenic_HyperMeth_lowCG | chr11 | 34194679 | 34195248 | - | AGGGGGAAAGGTTTTTAATTTTATATAT | CTCCCTCCTCCCTAATTCATAC | 569 | 506 | 52 | 30 | 52 | 0.22844 | lowCG | nonCGI | HyperMeth | ABTB2 |
| Human | H75 | Intragenic_HyperMeth_lowCG | chr7 | 47596208 | 47596787 | - | TGTTGGTTTTAAGGATTGTTTTGATT | TATAACTTCCTCCCAACAAATTCCTC | 579 | 483 | 46 | 31 | 46 | 0.22612 | lowCG | nonCGI | HyperMeth | TNS3 |
| Human | H76 | Intragenic_HyperMeth_lowCG | chr8 | 142499602 | 142500180 | - | GGGTTATGGAGAGTGATTTAAGAG | CTAAATATCCTTCTTCTAAAATCCTCA | 578 | 521 | 50 | 32 | 50 | 0.22477 | lowCG | nonCGI | HyperMeth | MROH5 |
| Human | H77 | Intragenic_HyperMeth_lowCG | chr15 | 42135720 | 42136293 | + | TTTAGGTGTTAAATTTTAGTTTTG | CTCATACCAAATATCTATCTCC | 573 | 502 | 45 | 28 | 45 | 0.216 | lowCG | nonCGI | HyperMeth | PLA2G4B |
| Human | H78 | Intragenic_HyperMeth_lowCG | chr1 | 36793406 | 36793977 | + | TTTGGGGAAGGTTTTTGAAATTTTT | CTTCCCTCCATCCAAAACCTAT | 571 | 506 | 46 | 33 | 46 | 0.20341 | lowCG | nonCGI | HyperMeth | SH3D21 |
| Human | H79 | Intragenic_HyperMeth_lowCG | chr2 | 101258854 | 101259433 | - | TTTTAGAATTGTGGGAGAAATAAATG | AAAACACCAAATTCTCTTCCTAAC | 579 | 514 | 49 | 33 | 49 | 0.19638 | lowCG | nonCGI | HyperMeth | PDCL3 |
| Human | H80 | Intragenic_HyperMeth_lowCG | chr17 | 60755563 | 60756133 | + | ATAGAAATTGAAGTTTAGGGAGAAGA | CTCCCTAAAATCACCCACATTC | 570 | 502 | 49 | 29 | 49 | 0.18419 | lowCG | nonCGI | HyperMeth | MRC2 |
| Human | H81 | Intragenic_HyperMeth_lowCG | chr1 | 3645737 | 3646316 | + | GAAAGAAGATTAGGGGATTTTGA | TCTAATCTTAATAAATCATAATCACTCT | 579 | 527 | 46 | 31 | 46 | 0.14433 | lowCG | nonCGI | HyperMeth | TP73 |
| Human | H82 | Intragenic_HyperMeth_lowCG | chr14 | 74942819 | 74943393 | - | TTAAGGGTTTTGGAAGAGGGTG | CCATCAAACCTCATCTTTCCTATATT | 574 | 516 | 52 | 32 | 52 | 0.14054 | lowCG | nonCGI | HyperMeth | NPC2 |
| Human | H83 | Intergenic_HypoMeth_highCG | chr5 | 72594738 | 72595311 | + | AAGGGAAGTTTATAAGAATTTTT | AAAACTAATTATAAAAACTCCTATCC | 573 | 498 | 45 | 29 | 45 | 0.9463 | highCG | CGI | HypoMeth | NA |
| Human | H84 | Intergenic_HypoMeth_highCG | chr13 | 50422057 | 50422626 | - | TATTGAGAGGTTTAAATGGGTAAA | AAACAATAAAACTATTCAAAATAAATAAAA | 569 | 503 | 38 | 35 | 38 | 0.78303 | highCG | CGI | HypoMeth | NA |
| Human | H85 | Intergenic_HypoMeth_highCG | chr18 | 76123522 | 76124096 | - | TGTGAAAGGTATTTAATGTATTAA | AAATCTCTTCTTATTATTAAACTCAC | 574 | 456 | 38 | 36 | 38 | 0.76696 | highCG | CGI | HypoMeth | NA |
| Human | H86 | Intergenic_HypoMeth_highCG | chr2 | 74942482 | 74943055 | - | AAGGATATAGAAAAGTTTTTAATTTTT | CAACTAAAAACTCCACCTAAATAA | 573 | 520 | 44 | 34 | 44 | 0.73327 | highCG | CGI | HypoMeth | NA |
| Human | H87 | Intergenic_HypoMeth_highCG | chr7 | 27265119 | 27265695 | - | TGATTTAAGGAAGAGATTAATTAGGT | CCCCATAAATCCAAAACATTCTC | 576 | 494 | 38 | 34 | 38 | 0.72487 | highCG | CGI | HypoMeth | NA |
| Human | H88 | Intergenic_HypoMeth_highCG | chr18 | 76674158 | 76674726 | - | TGTGGTGATAAATATTTAGGAAA | AACATCATTTTTCAATTCACAAAA | 568 | 505 | 48 | 25 | 48 | 0.68389 | highCG | CGI | HypoMeth | NA |
| Human | H89 | Intergenic_HypoMeth_highCG | chr10 | 26680682 | 26681241 | - | GTGGGTGTTTGGAGATAGGAATA | CTCCCCCAAAATCTAAAATACCT | 559 | 499 | 52 | 20 | 52 | 0.68265 | highCG | CGI | HypoMeth | NA |
| Human | H90 | Intergenic_HypoMeth_highCG | chr7 | 152591215 | 152591783 | - | AATTTAAGGGATTTTTGAGAATGAAT | TCAAAATTCCTCTCCCTTATCAATAT | 568 | 515 | 40 | 31 | 40 | 0.66317 | highCG | CGI | HypoMeth | NA |
| Human | H91 | Intergenic_HypoMeth_highCG | chr7 | 28893673 | 28894249 | + | TTGATTTTGGGTTTGAGAGTAAGAG | AAATCTTAAAATCCTCCTTTCTCTTC | 576 | 478 | 45 | 34 | 45 | 0.60387 | highCG | CGI | HypoMeth | NA |
| Human | H92 | Intergenic_HypoMeth_highCG | chr9 | 129484716 | 129485282 | + | TTTGATTTAGGTTTTGGGAATAT | CTAAACCCTCCAAAATAACCTAA | 566 | 518 | 46 | 35 | 46 | 0.59476 | highCG | CGI | HypoMeth | NA |
| Human | H93 | Intergenic_HypoMeth_highCG | chr4 | 8964975 | 8965553 | + | TATGGTGTGAAATTATTTTTGATT | AATTATCCTCCTCCAAATAAAAATAA | 578 | 487 | 43 | 34 | 43 | 0.56023 | highCG | CGI | HypoMeth | NA |

#### Supplementary Table 2 (cont.)

| Organism | ID | Class | Chromosome | Start | End | Strand | Fwseq | Rvseq | Full frag. length | GCH inform. length | GCH number | GCH MaxGap | WCG number | obs/exp CG | o/e CG level | CGI class | Methylation class | Promoter SYMBOL |
|---|---|---|---|---|---|---|---|---|---|---|---|---|---|---|---|---|---|---|
| Human | H94 | Intergenic_HypoMeth_mediumCG | chr2 | 86038046 | 86038624 | + | TTTATAGATGAGGGAATTGAGATTT | AATTATATACACAAAATTTTAATATTTCCT | 578 | 485 | 45 | 36 | 45 | 0.47197 | mediumCG | nonCGI | HypoMeth | NA |
| Human | H95 | Intergenic_HypoMeth_mediumCG | chr11 | 15959880 | 15960458 | - | GAGGATTTAGGGATTATTAGAAAGTA | AACACATAAATATAAATTCAAACCCT | 578 | 526 | 45 | 33 | 45 | 0.46498 | mediumCG | CGI | HypoMeth | NA |
| Human | H96 | Intergenic_HypoMeth_mediumCG | chr10 | 54565153 | 54565728 | + | TTTGTGAAATTATGTTTATGATAGT | ACCAAAACTACCATTTCAAAAA | 575 | 522 | 36 | 36 | 36 | 0.40252 | mediumCG | nonCGI | HypoMeth | NA |
| Human | H97 | Intergenic_HypoMeth_mediumCG | chr1 | 5675486 | 5676054 | + | AGTTTTAATATTAATTTAGTGAGAAAAT | TCATTTTTCTTTCACTATATCTACAT | 568 | 486 | 38 | 35 | 38 | 0.30636 | mediumCG | nonCGI | HypoMeth | NA |
| Human | H98 | Intergenic_HypoMeth_mediumCG | chr8 | 111746821 | 111747395 | + | TTGAGTTTGATTGAGTTTGAGTTTTT | AAAAATCCCCCAACACCTATAA | 574 | 521 | 47 | 29 | 47 | 0.27291 | mediumCG | nonCGI | HypoMeth | NA |
| Human | H99 | Intergenic_HypoMeth_lowCG | chr13 | 90080500 | 90081059 | - | TATTTAGGTTTTTATATTGGGATTGA | TAAATTAACCCAACCACAAAAATA | 559 | 505 | 34 | 36 | 34 | 0.1806 | lowCG | nonCGI | HypoMeth | NA |
| Human | H100 | Intergenic_HypoMeth_lowCG | chr17 | 19627313 | 19627872 | - | GGATAGTGGAGTTAAGAGGGATTTT | TAATTTCCCCTTCCAATCACTATC | 559 | 492 | 38 | 37 | 38 | 0.1563 | lowCG | CGI | HypoMeth | NA |
| Human | H101 | Intergenic_HypoMeth_lowCG | chrX | 149716689 | 149717268 | + | TATTTTTGAATTGTTAATGGAGGT | TTTCCTACCCTTCTCAATAAATA | 579 | 521 | 35 | 36 | 35 | 0.11254 | lowCG | CGI | HypoMeth | NA |
| Human | H102 | Intergenic_HyperMeth_highCG | chr7 | 386395 | 386965 | + | TTTGTTGGTTTGTTTTTGTGGAGAAT | CCACACCCCCTTATCACTAAAA | 570 | 476 | 57 | 28 | 57 | 0.7885 | highCG | CGI | HyperMeth | NA |
| Human | H103 | Intergenic_HyperMeth_highCG | chr6 | 170488003 | 170488575 | - | TGGTGTTTTTAGGATAGAGATAGG | CCTCAATCATCTTAAATCACCCA | 572 | 473 | 42 | 28 | 42 | 0.62829 | highCG | CGI | HyperMeth | NA |
| Human | H104 | Intergenic_HyperMeth_highCG | chr4 | 8910147 | 8910719 | + | AAGATTTTGTTATTTGGGTGGGAT | ATCTATAATTCCTCACCTAAATCACT | 572 | 488 | 45 | 36 | 45 | 0.55908 | highCG | CGI | HyperMeth | NA |
| Human | H105 | Intergenic_HyperMeth_highCG | chr7 | 1641765 | 1642344 | - | GAATTTAGGGGGATAAAATTTAATTT | CTATCTAAATCCCAATCTTCTCAC | 579 | 514 | 37 | 36 | 37 | 0.5375 | highCG | CGI | HyperMeth | NA |
| Human | H106 | Intergenic_HyperMeth_mediumCG | chr9 | 136100752 | 136101331 | + | GGGATTTGAATTTAGAGGTTTTTAG | TCCCACCCTAAAATTCAAAATT | 579 | 526 | 41 | 35 | 41 | 0.49467 | mediumCG | CGI | HyperMeth | NA |
| Human | H107 | Intergenic_HyperMeth_mediumCG | chr16 | 88498556 | 88499132 | - | TTTTTAGGAAATGAAAGTTTTTGGT | ATAAACTAAAAATCCAAAAATTAATCAC | 576 | 504 | 53 | 30 | 53 | 0.43221 | mediumCG | CGI | HyperMeth | ZNF469 |
| Human | H108 | Intergenic_HyperMeth_mediumCG | chr12 | 133018351 | 133018920 | - | ATATTTGTGAGGAATTAGAGAGGA | TTATTATATCCATTTCAAATTCAAAAAC | 569 | 466 | 39 | 29 | 39 | 0.40541 | mediumCG | CGI | HyperMeth | NA |
| Human | H109 | Intergenic_HyperMeth_mediumCG | chr4 | 827579 | 828146 | + | TTTGAGATTGGTGTTTTTAAGGTG | CTAACCCACACCTCTCTATATTAATT | 567 | 493 | 47 | 28 | 47 | 0.33396 | mediumCG | CGI | HyperMeth | NA |
| Human | H110 | Intergenic_HyperMeth_mediumCG | chr9 | 33434731 | 33435292 | - | TGGTTATATTTTTAGGTTAGAGTGAAATA | CCCACCCCAACATAAAAATAAAAA | 561 | 477 | 36 | 31 | 36 | 0.30001 | mediumCG | nonCGI | HyperMeth | NA |
| Human | H111 | Intergenic_HyperMeth_mediumCG | chr22 | 49580153 | 49580727 | - | AGTTTTTATTTGGAGTGAATTTGAGA | CCCCTCACTTCATTTCTCACTT | 574 | 497 | 43 | 30 | 43 | 0.29016 | mediumCG | CGI | HyperMeth | NA |
| Human | H112 | Intergenic_HyperMeth_mediumCG | chr2 | 240649498 | 240650061 | + | AGATGTTTATTTATTTGATTTTTAGGATAA | CAATCATCCTCCTAAAACTCTTTC | 563 | 472 | 52 | 26 | 52 | 0.27634 | mediumCG | CGI | HyperMeth | NA |
| Human | H113 | Intergenic_HyperMeth_mediumCG | chr9 | 130908096 | 130908663 | + | GAAATTAGAATGGGGAGTTAGATTG | CAATAAAAATACACAAATATTCACACT | 567 | 494 | 46 | 32 | 46 | 0.2638 | mediumCG | nonCGI | HyperMeth | NA |
| Human | H114 | Intergenic_HyperMeth_mediumCG | chr1 | 2794277 | 2794836 | + | AGGTTTTGAGGATAGAATTATAGGTT | CTCACTAATCTCCTAAAACCCATC | 559 | 488 | 44 | 31 | 44 | 0.25119 | mediumCG | nonCGI | HyperMeth | NA |
| Human | H115 | Intergenic_HyperMeth_mediumCG | chr10 | 133453020 | 133453587 | - | AGATTTTTATATTGTGTTTTTAAAGGAA | ATAAAACACACCTTAAAACACTCTAA | 567 | 486 | 43 | 30 | 43 | 0.25062 | mediumCG | nonCGI | HyperMeth | NA |
| Human | H116 | Intergenic_HyperMeth_lowCG | chr4 | 1755318 | 1755897 | - | TGTTTTATTTTAGGATTGGTGATAGA | TAAAACCTAAAACTCCAAAATCAAAA | 579 | 518 | 50 | 30 | 50 | 0.24001 | lowCG | nonCGI | HyperMeth | NA |
| Human | H117 | Intergenic_HyperMeth_lowCG | chr11 | 36280346 | 36280925 | + | ATAAAAAGGGTTTGGGAGGAGTATTT | TTCTCCTCTCTTCTATCCCTCTCC | 579 | 518 | 45 | 32 | 45 | 0.22898 | lowCG | nonCGI | HyperMeth | NA |
| Human | H118 | Intergenic_HyperMeth_lowCG | chr9 | 33416393 | 33416964 | - | GAGGTAGTGTGGATGAATATATA | TCTCTCAAATCATAAAATATAATAAAAA | 571 | 519 | 46 | 36 | 46 | 0.22049 | lowCG | nonCGI | HyperMeth | NA |
| Human | H119 | Intergenic_HyperMeth_lowCG | chr5 | 544345 | 544924 | - | TGGAGGAAATATAGTGATAATTTTT | TCCACTTCCTATTTAATTTTCATAA | 579 | 527 | 45 | 30 | 45 | 0.21992 | lowCG | nonCGI | HyperMeth | NA |
| Human | H120 | Intergenic_HyperMeth_lowCG | chr4 | 8897729 | 8898302 | + | TAAGGAGTGGATGGGGTTTGAG | CCTCCTCAACCTCCAAAATATAAAAA | 573 | 505 | 45 | 23 | 45 | 0.1773 | lowCG | nonCGI | HyperMeth | NA |
| Human | H121 | Intergenic_HyperMeth_lowCG | chr2 | 95932515 | 95933087 | - | ATTTTTAGAAGTGGAGTTTAGTGTTT | TAATCCCCAATAATCCCCCAAA | 572 | 511 | 46 | 34 | 46 | 0.17503 | lowCG | nonCGI | HyperMeth | NA |
| Human | H122 | Intergenic_HyperMeth_lowCG | chr10 | 126063779 | 126064354 | - | TTATATAGATTTTATTGTGGATTAAAGG | CCTCTCTTCAATCCATTTTCAA | 575 | 511 | 44 | 33 | 44 | 0.17225 | lowCG | nonCGI | HyperMeth | NA |
| Human | H123 | Intergenic_HyperMeth_lowCG | chr5 | 2191943 | 2192515 | + | GGTTTTATTTGGGATTTTAATAGTTT | AAAAACACTTAAACTTTTCAACA | 572 | 477 | 51 | 29 | 51 | 0.17092 | lowCG | nonCGI | HyperMeth | NA |
| Human | H124 | Intergenic_HyperMeth_lowCG | chr10 | 72939189 | 72939754 | - | ATGAAGGGTGAAATGGGAGGAA | ACAACATCCCCAAATAAAACAAATTC | 565 | 508 | 50 | 34 | 50 | 0.13685 | lowCG | nonCGI | HyperMeth | NA |
| Human | H125 | Intergenic_HyperMeth_lowCG | chr14 | 104776154 | 104776713 | + | TAATTGGGAGATGAAGTTATAGGTG | CTATCCATCTCTCCACACTATCT | 559 | 487 | 52 | 27 | 52 | 0.1354 | lowCG | nonCGI | HyperMeth | NA |
| Human | H126 | Intergenic_HyperMeth_lowCG | chr11 | 97622426 | 97622993 | - | AAGATTGAAAGGGTTGTAATATA | TAATCATAAAATCCTTCAAAAATCAC | 567 | 472 | 49 | 31 | 49 | 0.13515 | lowCG | nonCGI | HyperMeth | NA |
| Human | H127 | Intergenic_HyperMeth_lowCG | chr6 | 91980247 | 91980821 | - | ATTTATTAAATGGTGGAATTGAAAGA | ATTATCATAAAATCCTTAAATTATCACT | 574 | 517 | 49 | 25 | 49 | 0.13295 | lowCG | nonCGI | HyperMeth | NA |
| Human | H128 | Intergenic_HyperMeth_lowCG | chr9 | 99910533 | 99911112 | - | TGTGTGATTTAAGAAATGATTTTAAGT | AACCTAACATATCCCTTCTTTCC | 579 | 527 | 47 | 34 | 47 | 0.11085 | lowCG | nonCGI | HyperMeth | NA |
| Human | H129 | Intergenic_HyperMeth_lowCG | chr4 | 1756073 | 1756649 | + | TTTGAGATTTGAAGGGAGATAGG | CATCTAAAATCCATCTAACAATCACC | 576 | 523 | 57 | 28 | 57 | 0.10374 | lowCG | nonCGI | HyperMeth | NA |
| Human | H130 | Intergenic_HyperMeth_lowCG | chr4 | 176420924 | 176421501 | - | GTTTAGATGTGGGAGATATGGAA | AAAATTCCAAATCCTATACCTAAAAA | 577 | 527 | 56 | 25 | 56 | 0.08558 | lowCG | nonCGI | HyperMeth | NA |
| Human | H131 | Intergenic_HyperMeth_lowCG | chr8 | 63074469 | 63075031 | - | ATTTAAATGTGTAGTAATATTGGGTATAT | ATATTTCCCTCCTCCCCATTAT | 562 | 486 | 42 | 33 | 42 | 0.08446 | lowCG | nonCGI | HyperMeth | NA |
| Human | H132 | Enhancer_HypoMeth_highCG | chr9 | 98075244 | 98075823 | + | TGTTTATAGGGAGGTGTTTTATTATAG | AATAAAATTCACTCACAAAATCATCA | 579 | 472 | 40 | 30 | 40 | 0.65592 | highCG | NA | HypoMeth | FANCC |
| Human | H133 | Enhancer_HypoMeth_mediumCG | chr16 | 57125225 | 57125789 | + | TGGAAAGATAAGAGATAGATTTTGA | ACCAATTCTTAAAACACTTATCTAT | 564 | 507 | 41 | 32 | 41 | 0.40714 | mediumCG | NA | HypoMeth | NA |
| Human | H134 | Enhancer_HypoMeth_mediumCG | chr20 | 4065823 | 4066400 | + | AGGTTAGGTTTTTAAAGTATAGGATT | TCATTCCATAACTAATATTCTTAAAATC | 577 | 473 | 42 | 35 | 42 | 0.37005 | mediumCG | NA | HypoMeth | NA |
| Human | H135 | Enhancer_HypoMeth_mediumCG | chr10 | 60727320 | 60727899 | - | TTAGATAGGATTTTAGGGAGATTTTT | CTAAACTCACACACCTCAAAAA | 579 | 514 | 46 | 35 | 46 | 0.36263 | mediumCG | NA | HypoMeth | NA |
| Human | H136 | Enhancer_HypoMeth_mediumCG | chr4 | 10459838 | 10460397 | - | AATTTGATTTTATTTTTGAGGGAGATTT | ACAATCAATAACTCCCCACAAAATA | 559 | 490 | 45 | 36 | 45 | 0.32298 | mediumCG | NA | HypoMeth | NA |
| Human | H137 | Enhancer_HypoMeth_mediumCG | chr1 | 43471513 | 43472083 | - | AATAAAGATATTTTGGGAGGGAAGT | TCCATCTAATCTTTCTAAACACTAATC | 570 | 496 | 44 | 31 | 44 | 0.25238 | mediumCG | NA | HypoMeth | NA |

Supplementary Table 2 (cont.)

| Organism | ID | Class | Chromosome | Start | End | Strand | Fwseq | Rvseq | Full frag. length | GCH inform. length | GCH number | GCH MaxGap | WCG number | obs/exp CG | o/e CG level | CGI class | Methylation class | Promoter SYMBOL |
|---|---|---|---|---|---|---|---|---|---|---|---|---|---|---|---|---|---|---|
| Human | H138 | Enhancer_HypoMeth_lowCG | chr8 | 124934032 | 124934611 | + | GTTGGATTTTATTGATGATGATG | CTATTTCCTCCAATAACTCAAA | 579 | 511 | 43 | 35 | 43 | 0.24723 | lowCG | NA | HypoMeth | FER1L6 |
| Human | H139 | Enhancer_HypoMeth_lowCG | chr2 | 172962397 | 172962962 | + | TTAGGGAAATTTAGGATTATAGGT | ACATAATAACAACTCACTAAATTATA | 565 | 436 | 32 | 33 | 32 | 0.18588 | lowCG | NA | HypoMeth | NA |
| Human | H140 | Enhancer_HypoMeth_lowCG | chr20 | 42718542 | 42719119 | + | ATTTTTATTAAAGGGATTAGAGGTTT | ACTCTATAACTTAATACAACAACTAT | 577 | 498 | 41 | 33 | 41 | 0.17475 | lowCG | NA | HypoMeth | NA |
| Human | H141 | Enhancer_HypoMeth_lowCG | chrX | 58112122 | 58112687 | - | TTTTATTATTAGAATGAGTTGGGTTG | AAAAATAAACCTAACTAACCTCAA | 565 | 515 | 28 | 35 | 28 | 0.15189 | lowCG | NA | HypoMeth | NA |
| Human | H142 | Enhancer_HypoMeth_lowCG | chr14 | 23357117 | 23357696 | - | AAAGTGTATAAAGTTTTAAGGATAGG | AAACAAAAACCTCCTAAAAATCATT | 579 | 486 | 39 | 32 | 39 | 0.11561 | lowCG | NA | HypoMeth | NA |
| Human | H143 | Enhancer_HyperMeth_highCG | chr20 | 20432904 | 20433480 | - | AATATTTTAAATTATGTTGGATTTGTT | TATAATAACCCAAAAATCACCATTAT | 576 | 493 | 45 | 34 | 45 | 0.66521 | highCG | NA | HyperMeth | RALGAPA2 |
| Human | H144 | Enhancer_HyperMeth_highCG | chr7 | 2185569 | 2186129 | + | TGGTGAAAGGAATATTAAGTTATTA | TATCCATCTTTCCCTAATAATAAA | 560 | 467 | 45 | 34 | 45 | 0.61725 | highCG | NA | HyperMeth | MAD1L1 |
| Human | H145 | Enhancer_HyperMeth_highCG | chr19 | 2064731 | 2065298 | - | TTATGGGATAATGTTTTAGTGTG | AAAAATCACACCTAACTCCATA | 567 | 475 | 49 | 29 | 49 | 0.5388 | highCG | NA | HyperMeth | NA |
| Human | H146 | Enhancer_HyperMeth_highCG | chr20 | 48887107 | 48887666 | - | TTATTTGGAGAAGTGGTAAAGGT | TTATTATTTTTATTTCCCCTTTTCCT | 559 | 503 | 36 | 37 | 36 | 0.53018 | highCG | NA | HyperMeth | NA |
| Human | H147 | Enhancer_HyperMeth_highCG | chr7 | 1140412 | 1140987 | - | GGTTTTAGGTGGAGTTTATTTTAGTA | AAAAACAAAACAAAACCCTTTCTAA | 575 | 519 | 44 | 32 | 44 | 0.51223 | highCG | NA | HyperMeth | C7orf50 |
| Human | H148 | Enhancer_HyperMeth_highCG | chr13 | 114875873 | 114876452 | - | TTTGGATTATGATAGGTGTTTG | ACTCAAATCTCAATATCTAAAATAAA | 579 | 486 | 46 | 28 | 46 | 0.50139 | highCG | NA | HyperMeth | RASA3 |
| Human | H149 | Enhancer_HyperMeth_mediumCG | chr8 | 142286748 | 142287327 | - | GATTTTGATGAAGTTTATTAGTGTTA | ACTCACAAATATCCATAAATCA | 579 | 525 | 45 | 29 | 45 | 0.44126 | mediumCG | NA | HyperMeth | NA |
| Human | H150 | Enhancer_HyperMeth_mediumCG | chr8 | 1910137 | 1910712 | + | AGGTTTAGAAAGTGAAGTAAAAT | CCAATACCAATCTAACTAATACTAA | 575 | 527 | 46 | 31 | 46 | 0.36424 | mediumCG | NA | HyperMeth | NA |
| Human | H151 | Enhancer_HyperMeth_mediumCG | chr14 | 105201730 | 105202309 | - | GAAGGGGTTAGATATATTATAAGGATG | TAATCCCTAATCCCTAATCCCTC | 579 | 525 | 49 | 28 | 49 | 0.32225 | mediumCG | NA | HyperMeth | ADSS1 |
| Human | H152 | Enhancer_HyperMeth_mediumCG | chr14 | 95980752 | 95981331 | + | ATTGATAGATAAGGGGAAATTAAG | TTTCTATCTATCTCCATAAAAACAAA | 579 | 529 | 56 | 29 | 56 | 0.31193 | mediumCG | NA | HyperMeth | NA |
| Human | H153 | Enhancer_HyperMeth_lowCG | chr15 | 90578269 | 90578843 | - | TTAATTTAGATGAGATATGTGAGT | ATAATACTAAAATTTTAAATCCCTTAA | 574 | 511 | 51 | 29 | 51 | 0.2469 | lowCG | NA | HyperMeth | ZNF710 |
| Human | H154 | Enhancer_HyperMeth_lowCG | chr10 | 126386448 | 126387027 | + | GGAAATATTTGGGAAGAATTTAAAATG | TAAAATCCCCCTTAATCTAAAACAT | 579 | 527 | 58 | 32 | 58 | 0.20613 | lowCG | NA | HyperMeth | FAM53B |
| Human | H155 | Enhancer_HyperMeth_lowCG | chr12 | 31126085 | 31126662 | - | GAATTTATAGGATTTTATGGTGTTT | TAAAACTCCTAAATTCTATTCCT | 577 | 512 | 49 | 30 | 49 | 0.19538 | lowCG | NA | HyperMeth | TSPAN11 |
| Human | H156 | Enhancer_HyperMeth_lowCG | chr20 | 31126686 | 31127261 | - | TAAATGAGGGGAGTGAGATTTA | ACCTTCTAACATACACTAAATAATAT | 575 | 523 | 48 | 31 | 48 | 0.18881 | lowCG | NA | HyperMeth | NOL4L |
| Human | H157 | Enhancer_HyperMeth_lowCG | chr11 | 813390 | 813969 | + | TGATTTTAATGGAGTTTGGGTA | TTTATCCTCCCAAAACTAAAATAAT | 579 | 520 | 46 | 29 | 46 | 0.17546 | lowCG | NA | HyperMeth | NA |
| Human | H158 | Enhancer_HyperMeth_lowCG | chr11 | 65069313 | 65069891 | + | TTGGTTATTTTGGGTTTTTAGTTTTT | AAAATCCCCACCCTATTAAACAAA | 578 | 518 | 59 | 30 | 59 | 0.15712 | lowCG | NA | HyperMeth | NA |
| Human | H159 | ICR_Mat_HypoMeth | chr7 | 130132480 | 130132997 | - | TTGTTATTGAAATTTGAAAAATATTTAAG | AATAACCCTAATCTCACCCTAAT | 517 | 462 | 26 | 37 | 26 | 0.91478 | highCG | NA | HypoMeth | MEST |
| Human | H160 | ICR_Mat_HypoMeth | chr15 | 25199837 | 25200291 | - | GTTTTAAAAATTTGGAATATTTGATGA | TTTATTATAAAACCAAAAATTATTTCTTT | 454 | 337 | 37 | 36 | 37 | 0.62455 | highCG | NA | HypoMeth | SNRPN |
| Human | H161 | ICR_Mat_HypoMeth | chr20 | 57425425 | 57425977 | - | GATATGGGTGGGAGGTTTAATAG | AAAATCCCACAAACCCATAAAAATAT | 552 | 448 | 36 | 35 | 36 | 0.45514 | mediumCG | NA | HypoMeth | GNAS-AS1 |
| Human | H162 | ICR_Mat_HypoMeth | chr19 | 57349346 | 57349913 | - | GTGGTTTTATTTTGTGTATGGGG | CCCCTCAATCACTCAAAACAAA | 567 | 519 | 47 | 29 | 47 | 0.421 | mediumCG | NA | HypoMeth | ZIM2 |
| Human | H163 | ICR_Pat_HyperMeth | chr20 | 871766 | 872321 | + | AATGAATTGATGAAAAAGGGGAAATA | AAAATCACCACCTCTCCTAAAAA | 555 | 506 | 39 | 33 | 39 | 0.06985 | lowCG | NA | HyperMeth | ANGPT4 |

**Supplementary Table 2. List of human amplicons used in this study and their main characteristics.**

**“Organism”:** human, for which amplicons are designed. **“ID”:** amplicon identifier. **“Class”:** amplicon category. **“Chromosome”**, **“Start”** and **“End”** refer to the coordinates of human (hg19) genome assemblies. **“strand”**: DNA strand amplified. **“Fwseq”** and **“Rvseq”**: sequences of the forward and reverse primers, respectively. **“Full frag. length”:** the full length of amplicon fragment refers to the total number of base pairs covered by the amplicon. **“GCH inform. length”**: the GCH-informative length refers to the number of base pairs covered from the first to the last GCH trinucleotide. **“GCH number”**: number of GCH trinucleotides contained in the amplicon sequence. **“GCH MaxGap”**: maximum number of base pairs separating adjacent GCH trinucleotides. **“WCG number”**: number of WCG trinucleotides contained in the amplicon sequence. **“obs/exp CG”**: Observed/Expected CG ratio of the amplicon. **“o/e CG level”**: classification of CG levels based on Observed/Expected CG ratio of the amplicon: “highCG”: ≥ 0.5 observed/expected CG; “mediumCG”: <0.5 and ≥ 0.25 observed/expected CG, and “lowCG”: <0.25 observed/expected CG. The classification is based on all CGs within an amplicon irrespective of the identity of adjacent bases. **“CGI class”**: whether amplicon overlaps a CGI (CGI) or not (nonCGI), based on the UCSC Table Browser. **“Methylation class”**: methylation status of the amplicon. **“Promoter SYMBOL”**: name of gene(s) overlapping promoter amplicons (TSS+/- 500bp of reference genes).
